# Longitudinal investigation of spatial memory and retinal parameters in a 5xFAD model of Alzheimer’s disease reveals differences dependent on genotype and sex

**DOI:** 10.1101/2025.05.23.655771

**Authors:** Georg Ladurner, Conrad W. Merkle, Lucas May, Sybren Worm, Yash Patel, Maria Varaka, Magdalena Daurer, Laurenz Jauk, Roland Rabl, Philipp Königshofer, Gerhard Garhöfer, Manuela Prokesch, Bernhard Baumann

## Abstract

**Significance:** The retinal phenotype of Alzheimer’s disease (AD) is poorly understood. The connection between spatial memory and retinal phenotype is poorly investigated. Additionally, the influence of sex on the disease in mouse models is not sufficiently clear and requires further investigation.

**Aim:** To investigate the retina and spatial memory of 5xFAD mouse models of AD by measuring retinal and behavioral parameters.

**Approach:** A custom-built optical coherence tomography (OCT) system is used to image the retina of both eyes of 32 transgenic 5xFAD mice and 32 non-transgenic littermates over the course of 6 months (3-9 months of age) to investigate retinal parameters. The Morris Water Maze (MWM) test was performed to examine correlations between the retinal and spatial memory phenotype of the mouse model.

**Results:** Data were acquired in the form of OCT reflectivity images and OCT angiograms as well as video recordings of the MWM test. Layer thickness and vascular density were calculated from the resulting data. Behavioral data was extracted from the videos acquired from the MWM. Total retinal and inner retinal layer thickness increased slightly over the measurement period, while outer retinal layer and retinal nerve fiber layer thickness showed no significant change. The correlation analysis between MWM and layer thickness data revelated a positive correlation between inner nuclear layer thickness and MWM test day parameters.

**Conclusions:** OCT and MWM data revealed sex-based differences in the retinal phenotype of the 5xFAD mouse model, with changes in retinal thickness in different stages of the study and dissimilar correlations between retinal and spatial memory phenotype.

## 1 Introduction

Alzheimer’s disease (AD) is the most common form of dementia and represents a huge challenge for modern health care systems in an increasingly aging population.^1^ AD-related lesions in the brain include the appearance of amyloid beta plaques, neurofibrillary tangles as well as loss of neurons and synapses on a cellular level.^2^ On a behavioral level, clinical signs of AD include memory impairment, irritability, orientational troubles and in later stages also difficulties with basic body functions.^3^ Diagnosis of the disease is at this point not fully possible, a definitive diagnosis can only be achieved by postmortem neuropathology.^4^ Promising diagnostic approaches include magnetic resonance imaging, positron emission tomography and cerebrospinal fluid assays, in all cases in combination with neurological tests.^5^ Blood tests have recently emerged as a new alternative for the detection of AD biomarkers such as phosphorylated tau protein, amyloid beta or neurofilaments, although distinction between AD and non-related dementias can be challenging due to similar biomarkers.^6^ Treatment options for AD are still limited, due to a lack of methods to stop or reverse the disease progression.^7^ New compounds for AD treatment like donanemab and lecanemab recently received FDA approval, although their efficiency and safety have been disputed.^8^ The National Institute for Health and Care Excellence (NICE) even does reject the use of donanemab due to significant health risk associated with treatment and high costs.^9^ Due to a common embryological origin, the retina and the brain share similar functionalities as well as disease manifestations.^10^ For many neurodegenerative diseases, as for example Parkinson’s disease^11^ or amyotrophic lateral sclerosis (ALS),^12^ retinal pathologies in parallel to lesions in the brain have been reported. In the case of AD, markers such as inflammation, neurodegeneration as well as amyloid beta deposits and hyperphosphorylated tau aggregates have been reported to appear in the retina of patients even at an early stage,^10^ although controversial findings have also been published^13^. Whether the appearance of AD makers in the retina can be exploited for diagnostics purposes is still disputed.^14^

Given the difficulty of AD treatment^7^ and the challenges of diagnosis,^5,6^ it is crucial to increase the understanding of the retina as a potential diagnosis method of the disease.^15^ Mouse models are a central component of drug testing and the investigation of disease mechanisms^16^ and are thus an important subject to increase the understanding of the connection between retinal and cerebral pathologies of AD. The retinal phenotype of different mouse models of AD has been investigated, however reporting inconclusive results.^17^

One candidate technology for retinal diagnostics in AD is optical coherence tomography (OCT). OCT is a non-invasive imaging technique often used for *in vivo* retinal imaging, also in the context of neurodegenerative diseases.^18^ OCT is based on the interference of low-coherent light scattered by the sample with a reference beam to reveal information about the sample. OCT can be used to generate 3D images of tissue, also in real time.^19^ Modern OCT technology yields high-resolution images of vasculature^20^, charts retinal thickness^21^ and can visualize focal retinal lesions.^22^ Several mouse models of familial AD have been investigated using retinal OCT imaging. In an APP-PS1 mouse model, retinal thinning was observed in one study,^23^ whereas no changes could be measured by another group.^22^ Several studies reported thinning of retinal layers for the 3xTg mouse models.^17^ In previous investigations of the 5xFAD mouse model of AD thinning of the retinal nerve fiber layer (RNFL) and thickening of the inner plexiform layer (IPL) was measured with a commercial OCT device (Leica Envisu R2200) in 6–to 17-month-old mice.^24^ In a study by Kim et al., thinning of the total retina, the inner (IRL) and outer retinal layers (ORL) as well as the RNFL were reported with a spectral domain OCT (SD-OCT) system. Additionally, a decrease in capillary density was reported for this study with female mice.^25^ A third study using a commercial SD-OCT system reported thinning for the RNFL and thickening for the OPL and ONL for male transgenic 5xFAD mice and C57BL/6J controls at 3 months of age.^26^ Overall, the retinal phenotype of AD mouse models can be considered very controversial and demands careful clarification to enlighten the current landscape of conflicting results.

In addition, the connection of the retinal phenotype with cognitive impairment of the investigated mouse models is largely unexplored and leaves room for investigation. Moreover, the potential differences between female and male mice are often not investigated in the present literature. Here, by using a high-resolution OCT prototype tailored for multi-contrast retinal imaging in mice and combining the data with the assessment of cognitive impairment through behavioral testing, we aspire to gain new insights on the development of retinal parameters and unveil potential connections between the appearing phenotypes.

## 2 Materials and Methods

### 2.1 Animal Model

The 5xFAD mouse model (The Jackson Laboratory, strain #008730) is a common transgenic mouse model for the investigation of Alzheimer’s disease and has been used in about 10% of studies.^27^ The mouse line is based on a C57BL/6J background and overexpresses human amyloid beta amyloid precursor protein related to the Swedish (K670N, M671L), Florida (I716V), and London (V717I) familial Alzheimer’s disease (FAD) mutations as well as the human presenilin 1 (PS1) protein by harboring two FAD mutations, M146L and L286V. 5xFAD mice have been reported to present amyloid beta plaque formation in the brain as early as two months of age.^27^

### 2.2 Study Design

5xFAD mice (32 transgenic, 32 non-transgenic littermates) at 10 weeks (±1 week) of age were provided by Scantox Neuro GmbH. The animals were housed under controlled light conditions (12 hours dark, 12 hours light) with food and water ad libitum. Overall health status and weights were monitored every week. Animals were longitudinally investigated over the course of 6 months. OCT scans of animals were taken on five occasions at 12, 20, 24 and 36 weeks of age. Spatial memory was tested at 35 weeks of age using the Morris Water Maze (MWM). The exact number of animals and the number of transgenic (tg) and non-transgenic (ntg) animals involved in each test is listed in Table 1. Note that the animal numbers decrease over the course of the study because some animals were used for another investigation, which is not part of the research described in this work, and thus were extracted from the study at 12 weeks and 24 weeks of age. All experiments were performed in accordance with the ARVO Statement for the Use of Animals in Ophthalmic and Vision Research and Directive 2010/63/EU. All experimental procedures and protocols were approved by the ethics committee of the Medical University of Vienna and the Austrian Federal Ministry of Education, Science, and Research (GZ 2024-0.044.300).

**Table 1.** Number of animals used for each test.

| Age [weeks] |  | 12 |  | 20 |  | 24 |  | 35 |  | 36 |  |
| --- | --- | --- | --- | --- | --- | --- | --- | --- | --- | --- | --- |
| Number of mice examined with OCT |  | 64 |  | 48 |  | 48 |  | / |  | 27 |  |
| Transgenic | Non-transgenic | 32 | 32 | 24 | 24 | 24 | 24 | / |  | 14 | 13 |
| Number of mice tested in the MWM |  | / |  | / |  | / |  | 27 |  | / |  |
| Transgenic | Non-transgenic | / |  | / |  | / |  | 14 | 13 |  |  |

### 2.3 OCT System

The polarization-sensitive OCT (PS-OCT) system first presented by Fialová et al., was used to perform the study.^28^ The system was based on a super-luminescent diode with a central wavelength of 840 nm and a bandwidth of 100 nm, resulting in an axial resolution of ∼3.8 µm in tissue (n=1.35). The spectrometer line scan cameras acquired the spectral data with 3072 pixels for the co-and cross polarized channels with an A-scan rate of 80 kHz.^28^ By aligning the mouse placed on a mount providing three axes of translation and two axes of rotation, the imaged field of view was centered at the optic nerve head (ONH) and measured approximately 1mm x 1mm. Five repeated B-scans were acquired in 400 positions, resulting in volumes of 2000 B-scans consisting of 512 A-scans. The recorded volumes contained 512x400x3072 spectral voxels.

### 2.4 Anesthesia and Imaging

Mice were placed in an anesthesia induction chamber, which was afterwards filled with 4% isoflurane (IsoFlo, Zoetis Österreich GmbH) in oxygen for 4 minutes prior to imaging. Tropicamid drops (0.5%, Agepha Pharma s.r.o.) were applied to dilate the pupils of the animals. For imaging, the mice were transferred to a home-built animal mount and kept under anesthesia using a nose cone applying 2% isoflurane. In some cases, individuals developed resistance to the isoflurane during the study, requiring an increased concentration of 2.5% at later imaging timepoints to ensure proper sedation. To prevent hypothermia, the mouse was blanketed with a heating pad. Oculotect eye drops (Théa Pharma GmbH) were frequently applied to keep the eyes of the animals hydrated during the entire anesthesia sessions. Shortly prior to imaging, excessive eye drop liquid was carefully removed from the mouse eye using a cotton swab to avoid additional lensing effects. Both eyes were imaged with the ONH in the center of the field of view. Several male animals (n(tg) = 3, n(ntg) = 3) deceased during or after anesthesia and could not be investigated for all time points, but no female animals were affected.

### 2.5 Image Processing

Image processing was performed to provide images displaying reflectivity, motion and polarization-based contrast, using the pipeline described previously (Augustin et al., 2016). Retinal image data were flattened with respect to the retinal pigment epithelium (RPE) detected via its depolarizing properties in the PS-OCT images.^29^ Data sets that could not be processed in this first step due to low signal intensity, strong vignetting or acquisition errors, among others, were excluded from the study.

#### 2.5.1 Layer Thickness Analysis

To measure retinal layer thickness, the algorithm described by Augustin et al. was used (Augustin et al., 2018). The distance between RPE and inner limiting membrane (ILM) was considered as the total retinal thickness.^30^ Additionally the thicknesses of the outer retinal layers (ORL) and inner retinal layers (IRL) were measured, using the posterior surface of the outer plexiform layer as the boundary. The thickness of the retinal nerve fiber layer/ganglion cell layer complex (RNFL/GCL), the inner (INL), the inner (IPL) and outer plexiform layer (OPL), the photoreceptor complex (PRC) as well as the depolarizing RPE complex were also measured. Layer thickness data were stored as 2D en-face maps. To align measurements between all animals, the center of the ONH in each 3D volume was manually annotated and the central circle with 200 µm in diameter was removed from further analysis. The rest of the volume was divided into four sectors (superior, nasal and inferior, temporal), using the diagonals of the 1x1 mm² square as borders and into radial zones equidistant from the center. Then a zone with an inner radius of 200 µm and an outer radius of 600 µm distance from the ONH is selected. The average thickness of the total retinal thickness and the thickness of the sublayers was calculated in this annular zone. Thickness maps were manually screened during post processing and volumes with insufficient quality to produce a reliable segmentation of the retinal layers were excluded from the study. When two or more acquisitions of the same eye and timepoint were available, the scan with best signal quality or most centered ONH was used. When neither of the scans was significantly better than their counterparts, the resulting thickness measurements were averaged between both scans. This resulted in the number of thickness maps listed in Table 2.

**Table 2.** Number of individual volumetric OCT scans used for retinal layer thickness analysis.

| Age [weeks] |  | 12 |  | 20 |  | 24 |  | 36 |  |
| --- | --- | --- | --- | --- | --- | --- | --- | --- | --- |
| Number of analyzed volume scans |  | 125 |  | 87 |  | 86 |  | 50 |  |
| Transgenic | Non-transgenic | 61 | 64 | 41 | 46 | 44 | 42 | 25 | 25 |

#### 2.5.2 Angiography

OCT angiography (OCTA) data were computed as described in reference ^22^. Volumetric OCTA data were divided into three slabs using the layer segmentation described in section 2.5.1: the superior vascular plexus (SVP) in the RNFL/GCL complex, the intermediate capillary plexus (ICP) between the RNFL and the INL, and the deep capillary plexus (DCP) between the INL and the ONL. Using thresholding at 10 dB above the noise floor and a Frangi filter, binarized en-face images of the vascular plexuses were created, and a margin measuring 10 pixels at the border was set to zero to avoid border artifacts. Using the manually selected ONH position previously used for the quantification of retinal layer thickness, a circle around the ONH was cut out from the binarized images. The signal-to-noise ratio (SNR) was calculated for the regions visible in Figure 1(D), and the image was divided into four regions along the diagonals and then again into rings with 100 µm thickness, each of the zones has an individual SNR value. The binarized images were multiplied with the SNR map of the scan shown in Figure 1(D), thus assigning the values to the binarized image as exemplified in Figure 1(E). The regions whose SNR was below a predefined threshold were not used for further evaluation and were thus set to zero (see Figure 1(F)). For SVP and ICP, the threshold was set to 20 dB, while for the DCP, the threshold was set to 15 dB since the signal in this region was generally lower. In the case displayed in Figure 1, the regions in the corners and around the ONH did not display above threshold SNR and were removed. By inverting only the pixels in the regions with above-threshold SNR, a negative image was created. By counting the positive pixels in the regions with above-threshold SNR (Figure 1(G)) and the negative pixels in the same regions (Figure 1(F)), the area vessel density was calculated for each vascular plexus.

**Fig 1:**
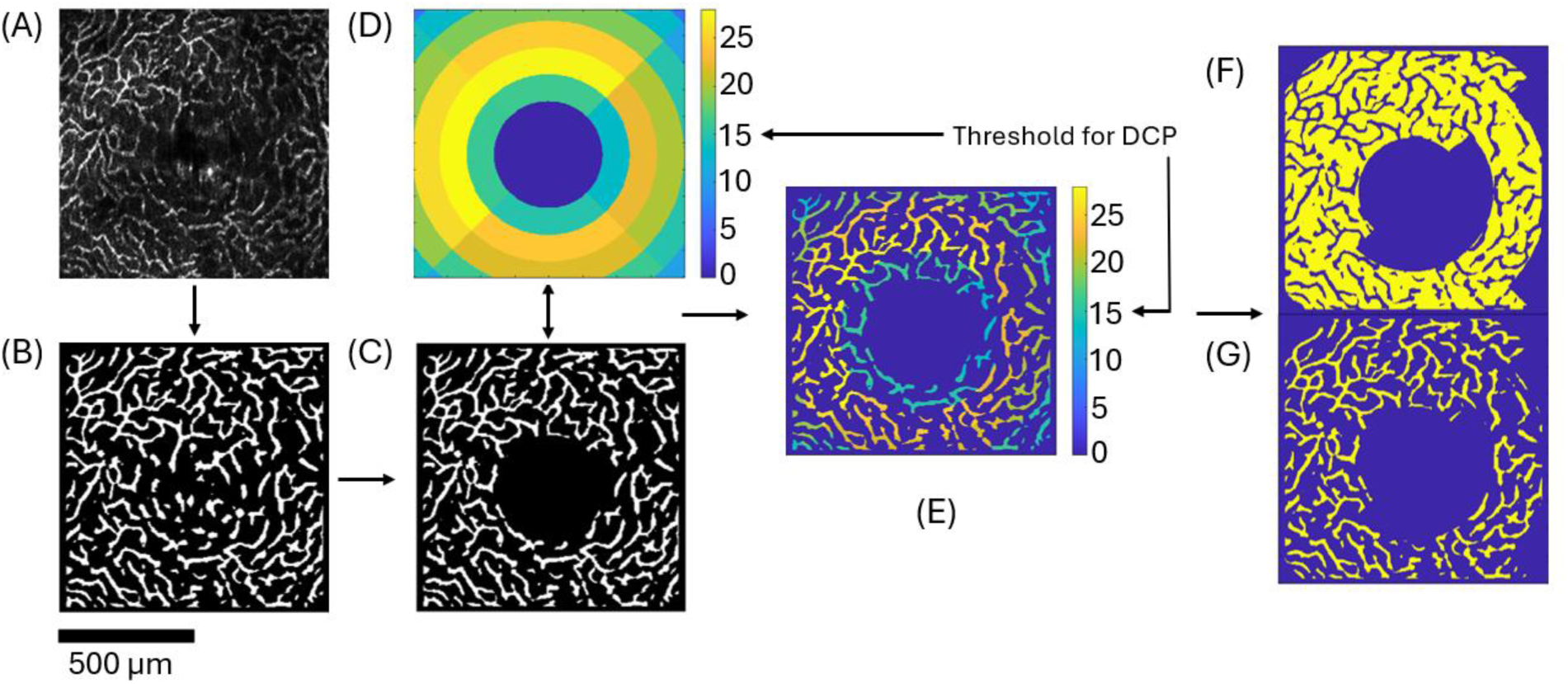
Example for the calculation of vessel density in the DCP. (A) Raw OCTA data after layer segmentation of the 3D stack, (B) binarized image with borders set to zero (C) removal of the circular region centered at the ONH from the binarized image, (D) SNR sector map, rings have a thickness of 100 µm (E) application of the SNR on the binarized image, (F) removal of the regions with SNR below threshold and creation of the negative for counting pixels.

The OCTA data was sorted similar to the retinal layer analysis. Additionally the data was screened manually, to only include the best available datasets. Table 3 presents the number of datasets that yielded at least one measurement for the 3 different vascular regions and thus were used for the longitudinal OCTA analysis.

**Table 3.**
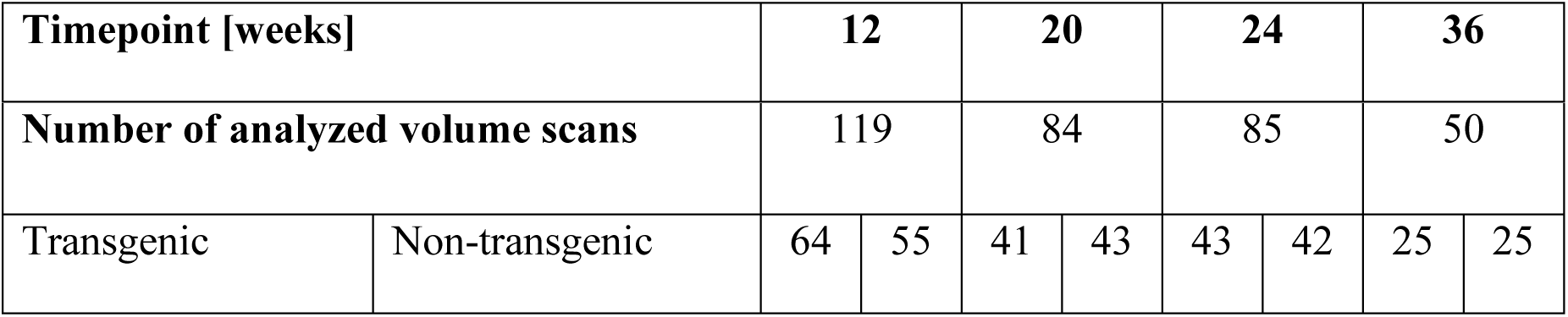
Number of individual volumetric OCTA datasets used for angiography.

### 2.6 Morris Water Maze

Spatial memory capabilities of the animals were tested in a custom-built Morris Water Maze (MWM) setup.^31^ The maze consisted of a water filled pool, one meter in diameter and surrounded by opaque curtains, a translucent platform (8 cm diameter) and four distinct landmarks as illustrated in Figure 2(A). A monochrome camera with 5 megapixels resolution and 14 fps (Basler Ace Classic, acA2500-14uc) was mounted 1.5 m above the pool and the landmarks placed in the middle of each of the four sides surrounding the pool, to avoid any additional orientation points for the mice. Movie data were acquired by using the Basler Video Recording software (version 1.3). The northeastern quadrant contained a translucent platform (target zone) about 1 cm below water level, such that it was invisible for the mice. The pool temperature was kept between 21-22°C, and the light intensity at the water surface level was controlled to be around 50 lux. The pool was divided into four quadrants to randomize the starting position (see Figure 2(B)). The starting positions were defined as per Table 4 and were applied to every mouse to assure an equal distribution of starting positions.

**Fig. 2:**
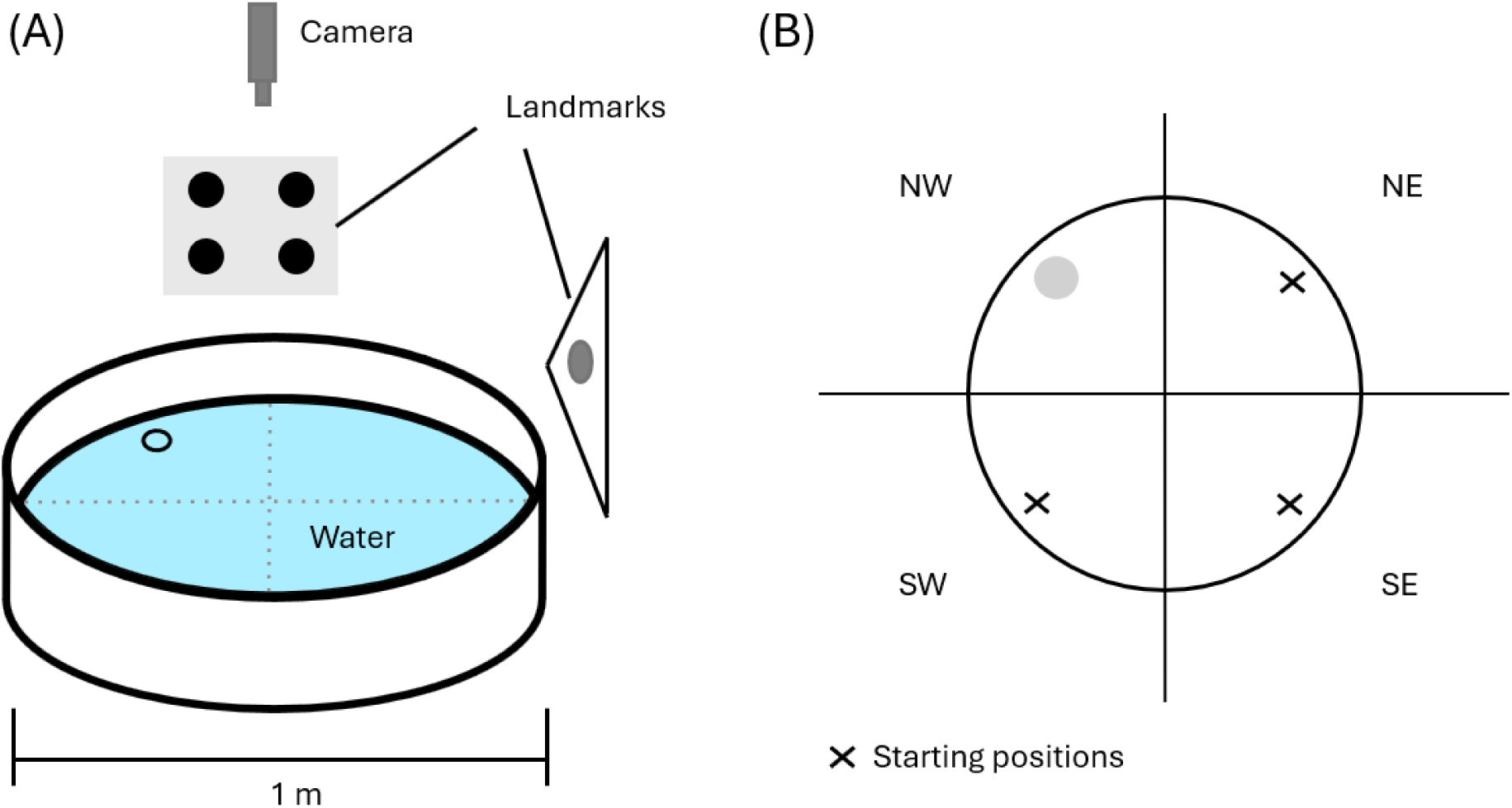
Schematic of the Morris Water Maze set-up. (A) Division of the pool into quadrants for the randomization of starting positions. (B) Schematic of the pool set up with landmarks and camera placement.

**Fig. 3:**
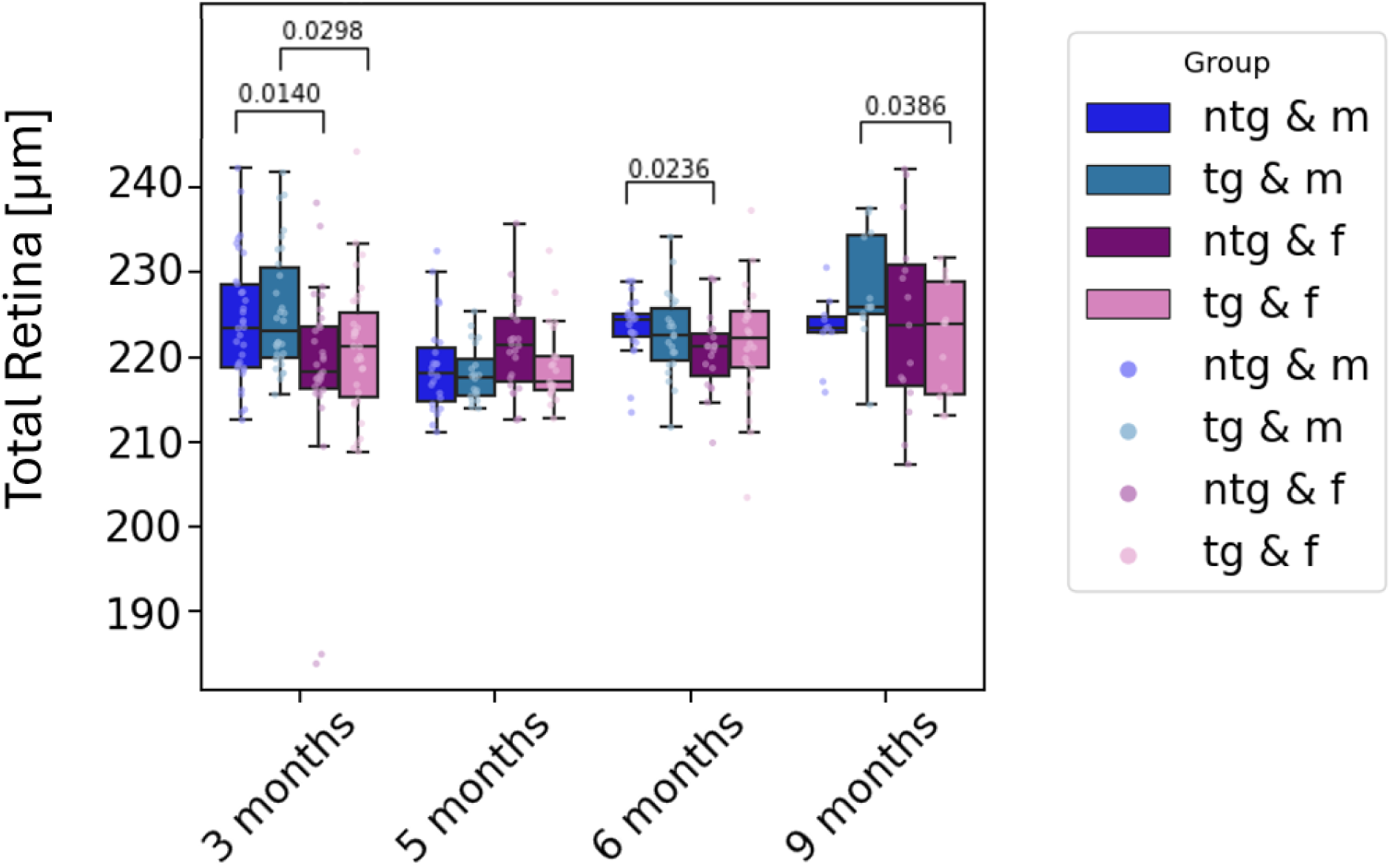
Longitudinal measurement of the total retinal thickness for all mice divided by genotype and sex. Individual points represent singular measurements for one mouse eye. Significant differences between groups are marked with brackets above the boxes and annotated with the corresponding p-values.

**Table 4.** Starting positions for each training and the test day. Quadrants: NE = northeast, SE = southeast, SW = southwest, NW = northwest.

| Trial | 1 | 2 | 3 | 4 | 5 |  |
| --- | --- | --- | --- | --- | --- | --- |
| Task | Training | Training | Training | Training | Test day | Platform position (for all trials) |
| Day 1 | NE | SE | SW | SE | / | NW |
| Day 2 | SW | NE | SE | SW | / | NW |
| Day 3 | SE | SW | NE | SW | / | NW |
| Day 4 | NE | SE | SW | NE | / | NW |
| Day 5 | / | / | / | / | SE | / |

The testing protocol included four consecutive training sessions and one test day. A training day consisted of four trials, where each mouse was placed in the pool (in varying quadrants as outlined in Table 4) for a maximum of 60 seconds and had to find the hidden platform. When a mouse was not able to find the platform within this time, it was placed on the platform and left there for ten seconds to memorize the position. In case the mouse managed to find the platform, the recording was stopped, and the mouse was removed from the pool. The time between daily trials was 10 minutes. On the test day, the hidden platform was removed, and the mice were placed in the pool once for one minute. Each trial was recorded using the top-down mounted camera.

For data evaluation an automatic tracking system, namely Noldus EthoVision XT 14, was used. For the training days, latency to find the platform [s], distance traversed [m], as well as the percentage of time spent floating and thigmotaxis (time spent close to the pool walls) was measured. ^32^ For the test day, the abidance in each of the of the quadrants and the number of crossings of the target zone were evaluated. The resulting data were analysed in GraphPad Prism^TM^ 10.

### 2.7 Statistical Analysis

For the longitudinal analysis of the retinal parameters, a mixed intercepts model was applied to investigate age dependent effects and account for multiple measurements per mouse. For comparison between groups at each timepoint, a t-test was applied. P-values smaller than 0.05 were considered significant.

To assess whether retinal layer thickness and vessel density data were correlated with spatial memory impairment, the OCT and MWM parameters were analyzed using linear regression. If data for both eyes of a mouse was available, the average of the two values was used as one single datapoint for the analysis. Additionally, the weight of the animals was also included in the analysis. Linear correlation coefficients, R-squared, mean squared error (MSE) intercept and the linear regression coefficient were calculated using a custom Python script. Pearsons’s correlation analysis was used to calculate the correlation coefficients and the corresponding p-values.

## 3 Results

### 3.1 Longitudinal Development of Retinal Layer Thickness

The longitudinal analysis of the total retinal thickness reveals an overall thickening of the retina for male animals by 5.7 µm (p=0.0029) from the beginning to the end of the study. Overall retinal thickness increases significantly by 5.8 µm (p=0.0051) for all animals in the study duration. Significant differences between groups were observed between tg male and female animals, as well as for ntg male and female animals at 3 months of age. Tg male animals (225.4±7.1µm) had significantly thicker retinas than tg female animals (221.1±7.8µm) (p=0.014), also for ntg animals, male mice (224.4±7.3µm) displayed a higher total retinal thickness than female animals (218.5±10.7µm) (p=0.0298). Other significant differences were found between male and female ntg animals at 6 months of age, where male animals (223.6±3.6µm) showed significantly higher total retinal thickness than females (220.5±4.5µm) (p=0.0236), and between male and female tg mice at 9 months of age: Male tg mice had significantly thicker retinas (228.0±6.4µm) than female tg mice (220.0±6.5µm) (p=0.0386) for the latest measurement point.

Following the analysis of total retinal thickness, the retinas were segmented into IRL (RNFL, IPL, INL) and ORL (PRC, RPE and OPL) to evaluate the thickness of these retinal sublayers. Figure 4 shows the time development of the IRL (A) and the ORL (B). The mixed effects model yielded a significant thickness increase of the ORL of 3.8 µm (p=0.0048) for all mice over the course of the investigation. Significant thickening of the IRL in comparison to the measurements at 3 months of age could not be detected (p=0.0503). Significant group differences for the IRL measurements were observed at 3, 5 and 9 months of age. At 3 months of age male, tg mice (97.3±3.7µm) showed a significantly higher IRL thickness than female tg mice (94.7±5.0µm) (p=0.0215). This was also observed for ntg mice, where male mice (96.7±4.1µm) had thicker IRL than female ntg mice (94.4±3.9µm) (p=0.0228). For 9 months old mouse models, the IRL thickness of tg male mice (98.16±3.19µm) was significantly larger than for ntg male mice (95.4±2.1µm) (p=0.0363), and also in comparison with female tg mice (94.7±4.0µm), male tg mice had significantly (p=0.0331) thicker IRL. For the measurements of the ORL, no significant changes were observed for the last measurement at 9 months of age.

**Fig. 4:**
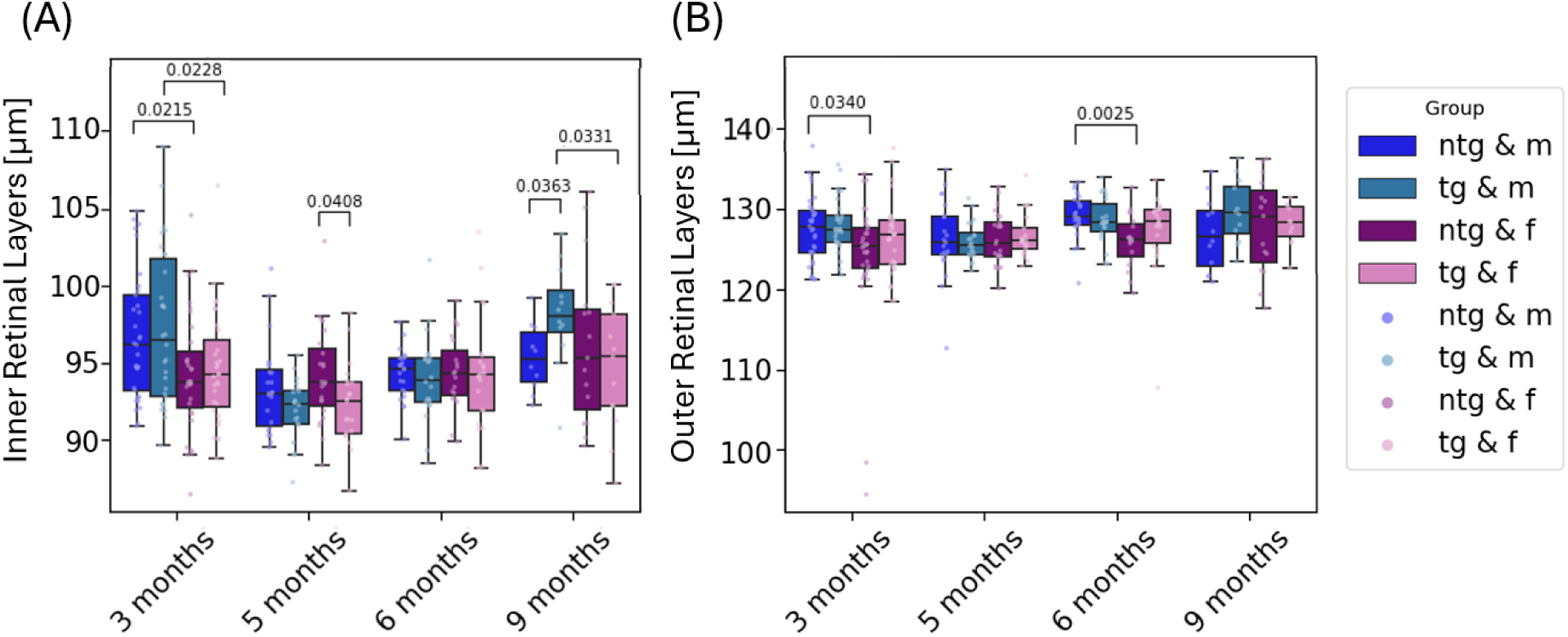
Longitudinal measurement of the IRL (A) and ORL (B) for all mice divided by genotype and sex. Individual points represent singular measurements for one mouse eye. Significant differences between groups are marked with brackets above the boxes and annotated with the corresponding p-values.

**Fig. 5:**
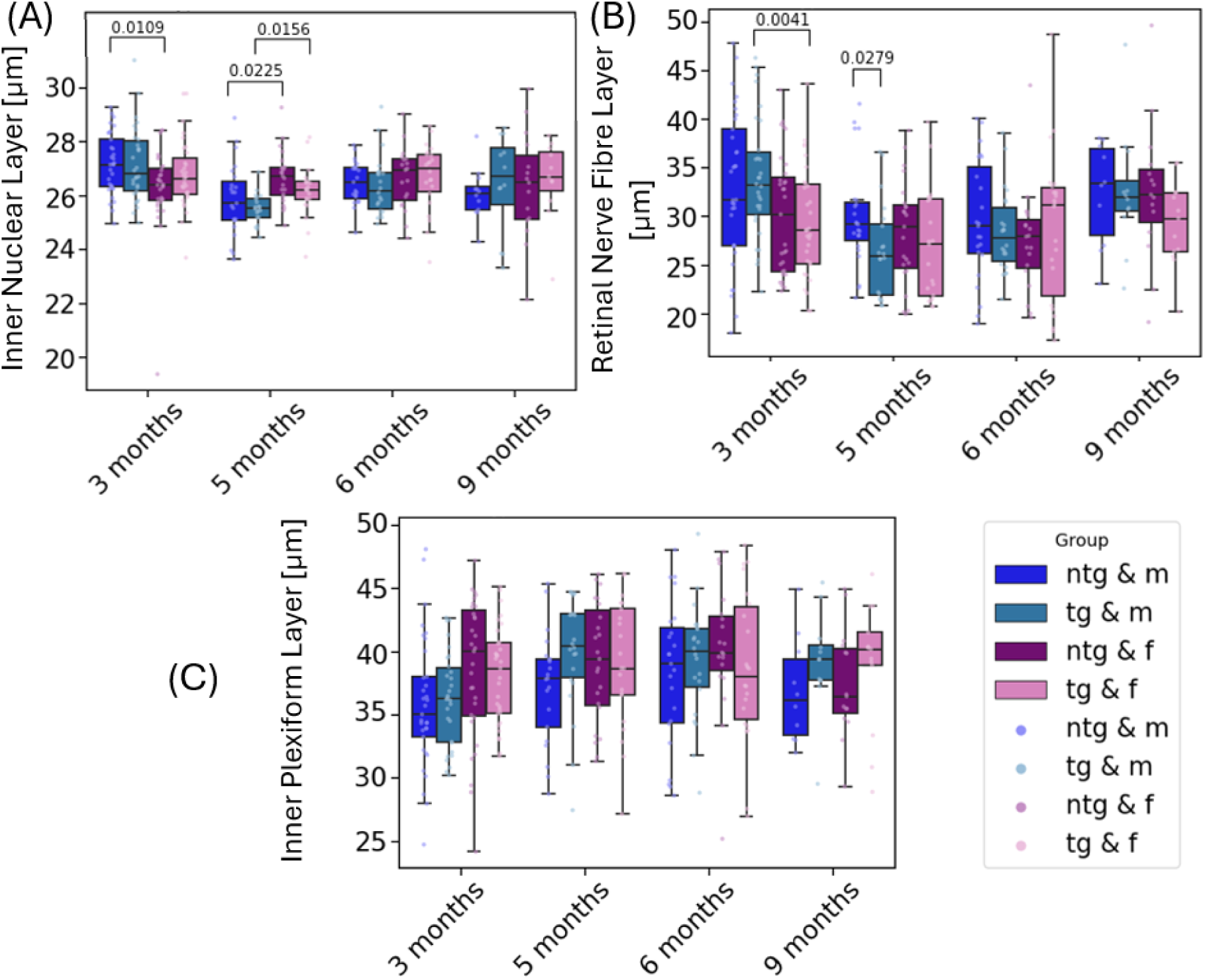
Longitudinal measurement of the INL (A), RNFL (B) and IPL (C) for all mice divided by genotype and sex. Individual points represent singular measurements for each one mouse eye. Significant differences between groups are marked with brackets above the boxes and annotated with the corresponding p-values.

As a next step, the sublayers of the IRL (INL, RNFL and IPL) were individually investigated to assess potential changes over time and between groups. Neither of these retinal sublayers showed significant changes in thickness over the study time of 6 months. For the INL measurements, the analysis yielded significantly different thicknesses for male and female tg animals, where male tg animals displayed a thickness of 26.8 ±1.3µm, significantly thicker than female tg animals with 27.2 ±1.4µm (p=0.0109). Significant thickness differences also appeared between ntg male and female, as well as between tg male and female mice at 5 months of age. The investigation of RNFL thickness showed significantly higher values for male tg animals (33.9±6.2µm) than for female tg mice (29.4±5.6µm) (p=0.0041). For 5 months of age, a thicker RNFL for ntg male animals than for tg male animals was observed as well. For the final measurement with 9-month-old animals, no significant differences were observed for the three inner retinal layers.

The ORL consists of the PRC, the OPL and the RPE. Figure 6 shows the development of thickness over time for these layers. An overall thickening of the PRC for all animals by 2.6±1.3µm (p=0.0350) over the study duration was observed for. Significant differences between groups were found for ntg male and female animals for 3 and 6 months of age. However, similar changes in layer thickness were not observed for the measurements at 5 and 9 months of age. For the OPL, an overall thickening of the layer was measured (1.1±0.3µm) (p=0.0003) for all animals. At the last measured timepoint, female ntg animals showed a significantly thicker OPL (12.7±1.2µm) compared to female tg animals (11.7±0.6µm) (p=0.0128).

**Fig. 6:**
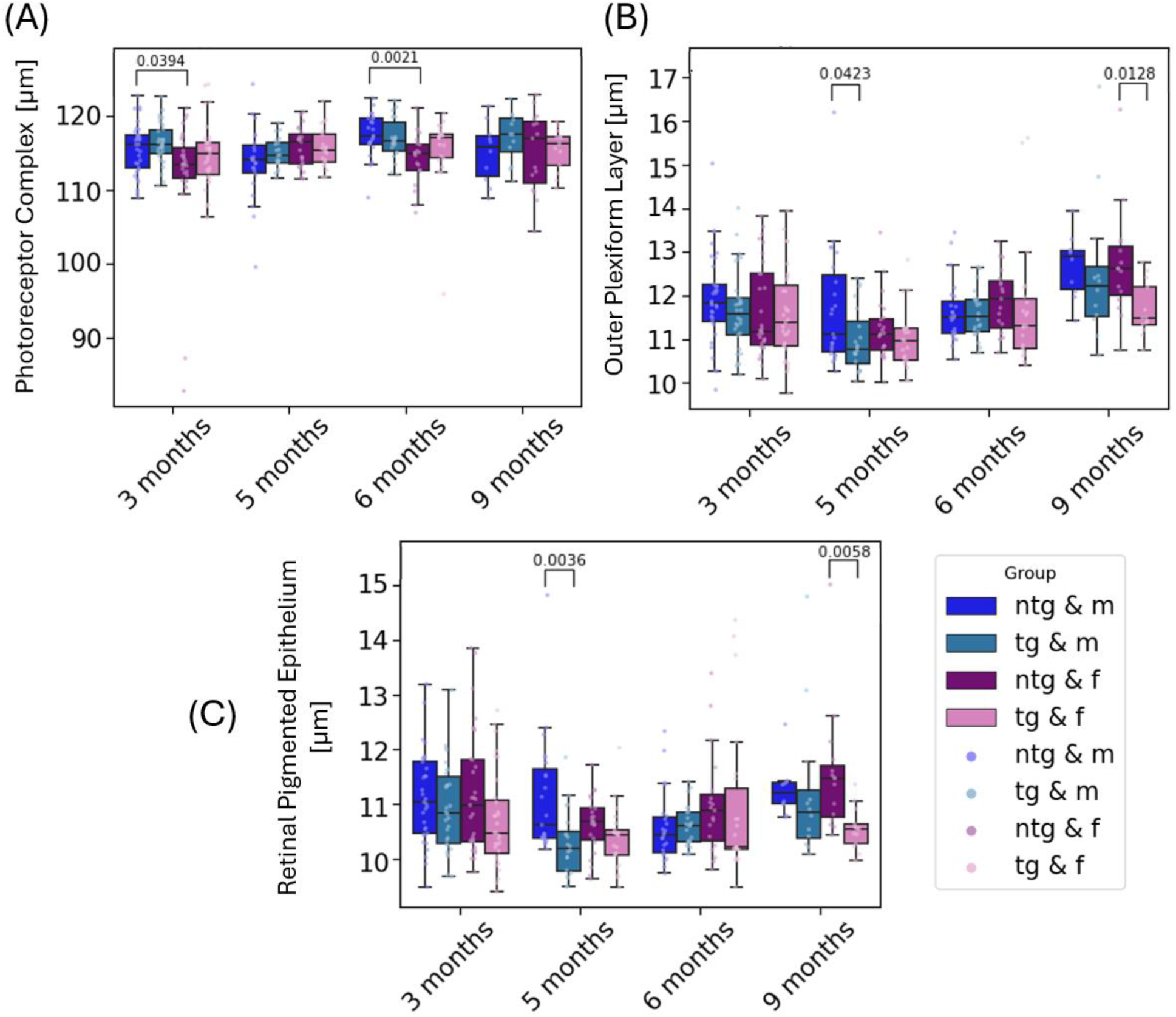
Longitudinal measurement of the PRC (A), OPL (B) and RPE (C) for all mice divided by genotype and sex. Individual points represent singular measurements for one mouse eye. Significant differences between groups are marked with brackets above the boxes and annotated with the corresponding p-values.

For the RPE data, the mixed effects models showed an overall lower RPE thickness for tg animals of about 0.5±0.24µm (p=0.036). Changes over time for the RPE were not identified. Similar to the OPL, female ntg animals had a different RPE thickness than tg female animals at 9 months of age. With 11.6±1.1µm, the female ntg animals display a significantly thicker RPE than the tg animals with 10.6 ±0.4µm (p=0.0058). Note that the RPE thickness measurements provided here reflect the segmented thickness of the depolarizing layer.^28^ Further significant changes were measured for PRC thickness between male (115.7±3.5µm) and female (112.5±7.8µm) ntg animals at 3 months of age (p=0.0394), between tg (11.0±0.7µm) and ntg (11.7±1.3µm) male animals at 5 months of age for OPL (p=0.0423) and for tg (10.3±0.6µm) and ntg (11.1±1.0µm) male animals at 5 months of age for the RPE thickness (p=0.0036).

### 3.2 Angiography

Significant differences were observed between male ntg and tg mice at 9 months of age for the SVP and ICP measurements. The SVP density for ntg male mice (19.8±1.6%) was significantly lower (p=0.0063) than for male tg mice (22.1±1.5%). The mixed intercepts model revealed a significant (p=0.025) density decrease of about 2% for male mice over the course of the study from 3 months until 9 months of age. Significant differences were also observed between ntg male and ntg female mice with lower values for ntg male mice (p=0.0047) at 9 months of age and for female ntg and female tg mice at 5 months of age (p=0.0365, see Figure 7(A)). Male ntg mice also experienced an ICP density decrease over the course of the longitudinal investigation. The mixed effects model yielded a significant decrease of 3.6% in density for all male mice (p=0.039) and 5.4% for male ntg mice (p=0.025) from the baseline measurement to the endpoint. Significant differences between ntg male and tg male mice can be observed at 9 months of age. With 6.8±1.4%, ntg male mice showed significantly lower density than tg male mice with 9.9±2.5% (p=0.0023). Male ntg mice also had significantly lower densities than female ntg mice (p=0.0292). Ntg male mice also showed significantly lower ICP density than female ntg mice at 5 months of age (p=0.0207, see Figure 7(B)). In contrast to these findings in the SVP and ICP, no significant changes over time or between individual groups were observed for the measurement of the DCP density (Figure 7(C)).

**Fig. 7:**
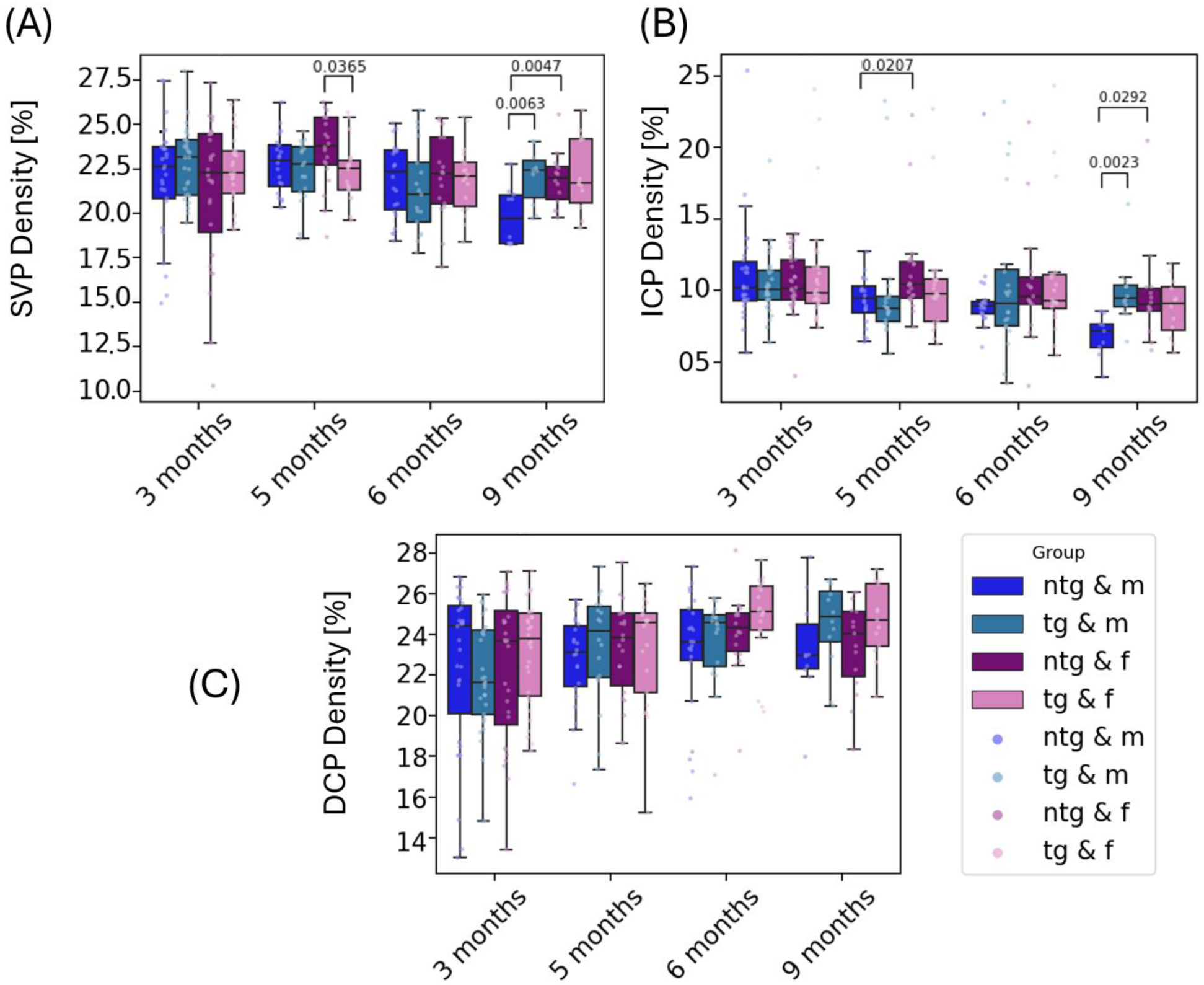
Longitudinal measurements for vascular density in percentage of area (A) SVP density, (B) ICP density, (C) DCP density. Significant differences between groups are marked with brackets above the boxes and annotated with the corresponding p-values.

### 3.3 Spatial Memory Testing

#### 3.3.1 MWM at 35 weeks of age

All remaining animals (n=27) were retested in the same MWM set-up at the age of 35 weeks. As shown in Figure 8, no significant differences between the tg and ntg animals can be observed for the experiment. For female animals, significant differences were observed for the latency to find the platform on the fourth training day (Figure 9 (A)). For the other parameters, no changes can be observed.

**Fig. 8:**
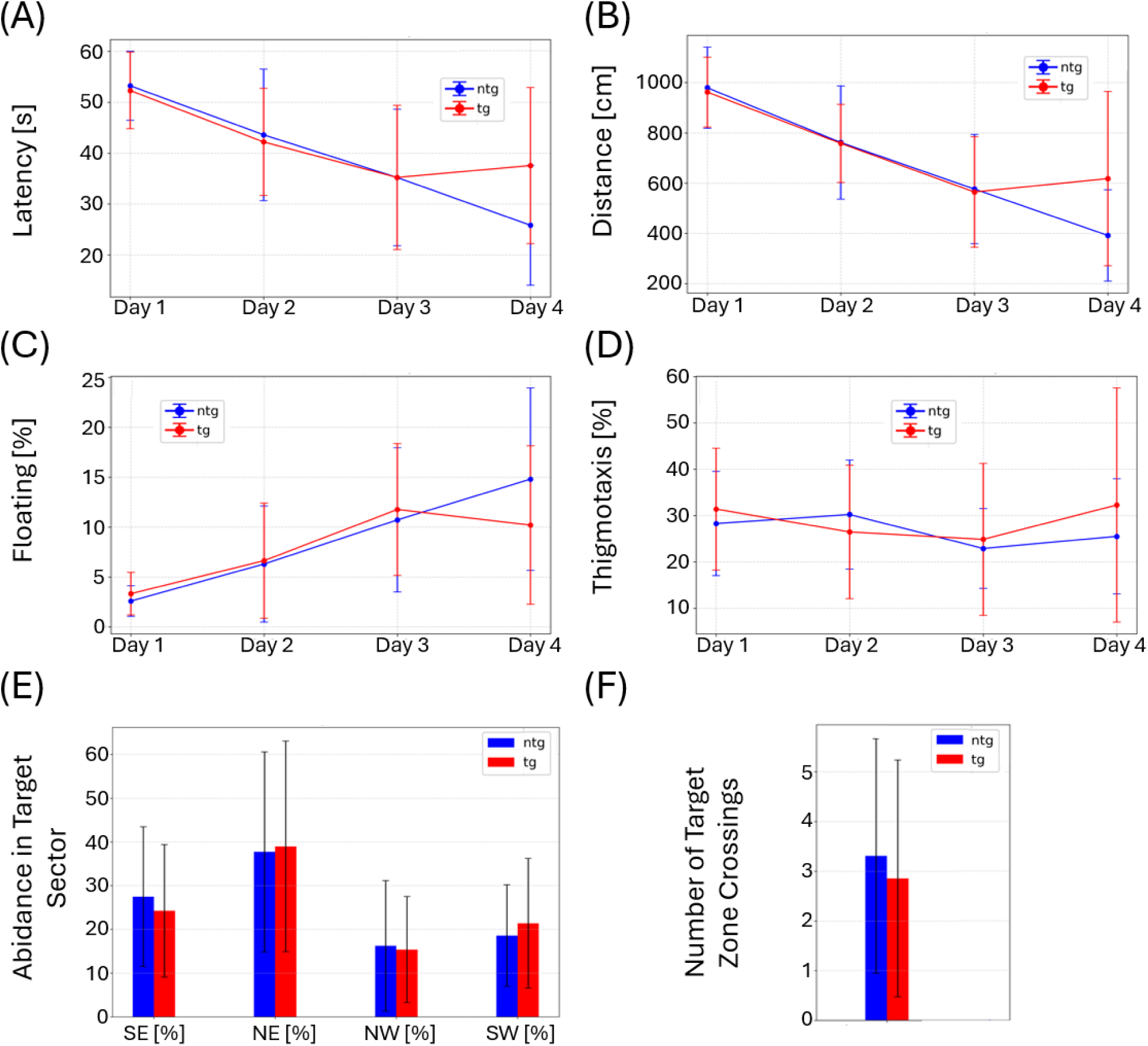
MWM results for animals at 35 weeks of age. (A) Latency to find the target platform, (B) distance traversed, (C) floating behavior, and (D) thigmotaxis over the four training days. (E) Abidance in target quadrant and (F) number of target zone crossings on the test day. Whiskers indicate ± the standard deviation (SD).

**Fig. 9:**
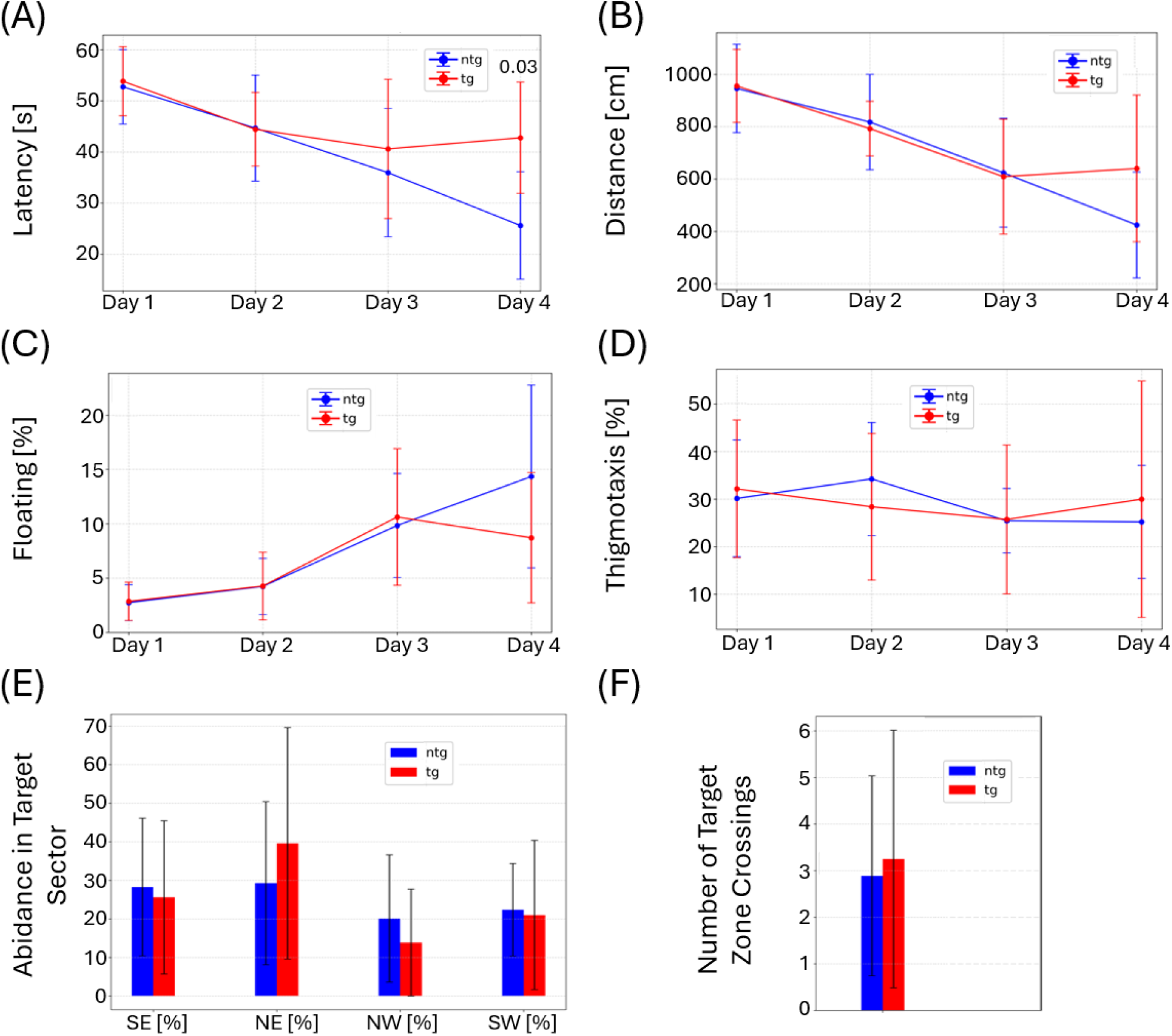
MWM results for female animals at 35 weeks of age. (A) Latency to find the target platform, (B) distance traversed, (C) floating behavior, and (D) thigmotaxis over the four training days. (E) Abidance in target quadrant and (F) number of target zone crossings on the test day. Whiskers indicate ± the standard deviation (SD).

**Fig. 10:**
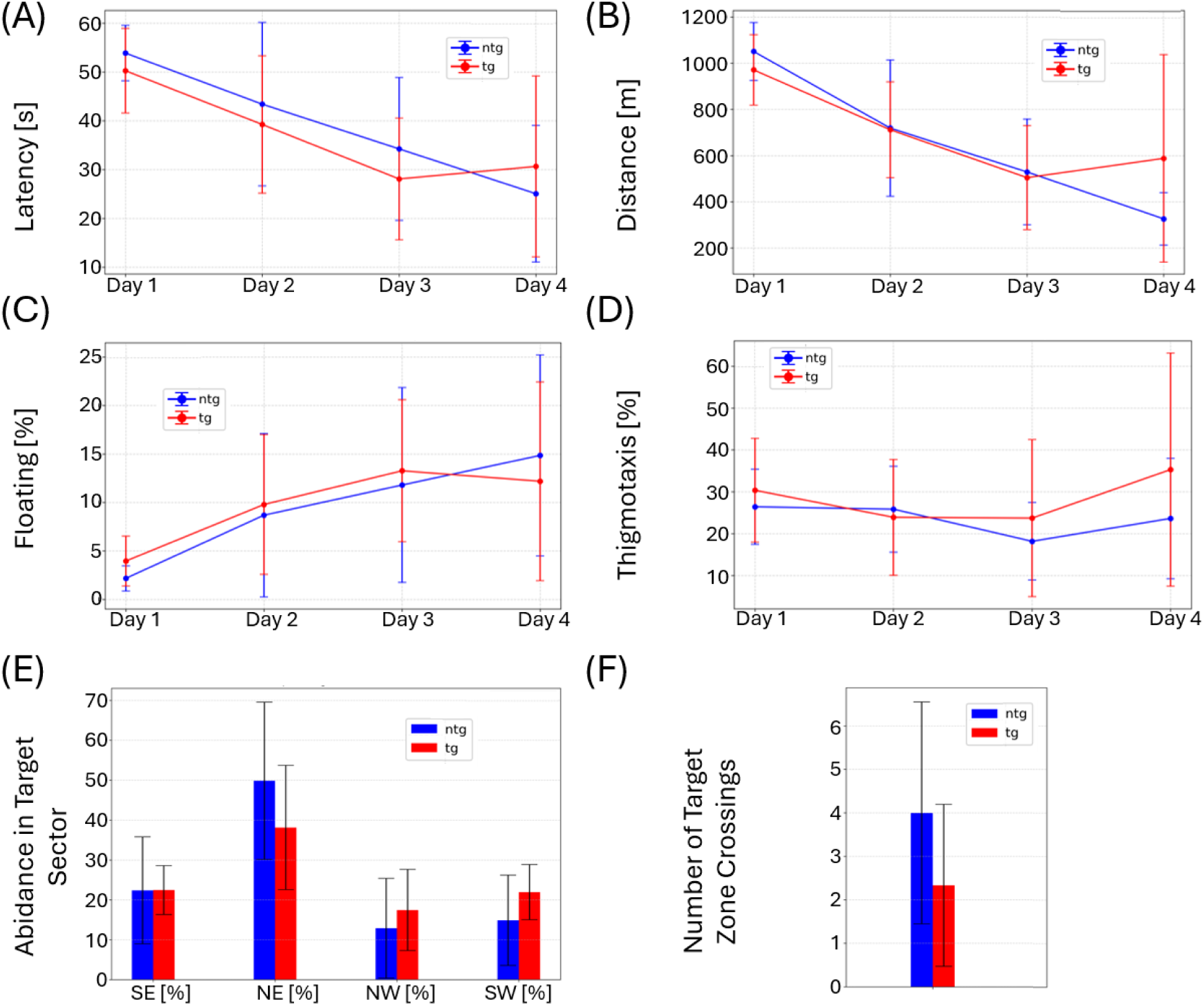
MWM results for male animals at 35 weeks of age. (A) Latency to find the target platform, (B) distance traversed, (C) floating behavior, and (D) thigmotaxis over the four training days. (E) Abidance in target quadrant and (F) number of target zone crossings on the test day. Whiskers indicate ± the standard deviation (SD).

Male animals displayed no significant differences in the comparison with ntg animals in the MWM test.

### 3.4 Correlation between retinal parameters and spatial memory

The correlation analysis between retinal parameters and values obtained from the spatial memory testing revealed a significant association between the INL thickness and the MWM parameters of the test day (day 5) for female transgenic mice. The number of target zone crossings (Figure 11 (D-F)) as well as the percentage of time spent in the target sector (Figure 12 (D-F)) correlated strongly with the thickness of the INL (0.87, 0.86), the IRL (0.77, 0.74) and the total retina (0.76, 0.69). Pearson’s correlation coefficient was the highest for the INL with 0.87 (p=0.0054) for the number of target zone crossings and 0.86 (p=0.0060) for the time spent in the target sector. These correlations were only observed for female transgenic mice. For male transgenic mice, no correlation between layer thickness measurements and MWM parameters was observed for the test day. Combining the two groups and looking at the correlation for all transgenic animals revealed a statistically significant correlation between the two investigated MWM parameters and the IRL (0.54) and INL (0.62), respectively, as shown in Figures 11 (B, C) and 12 (B, C).

**Fig. 11:**
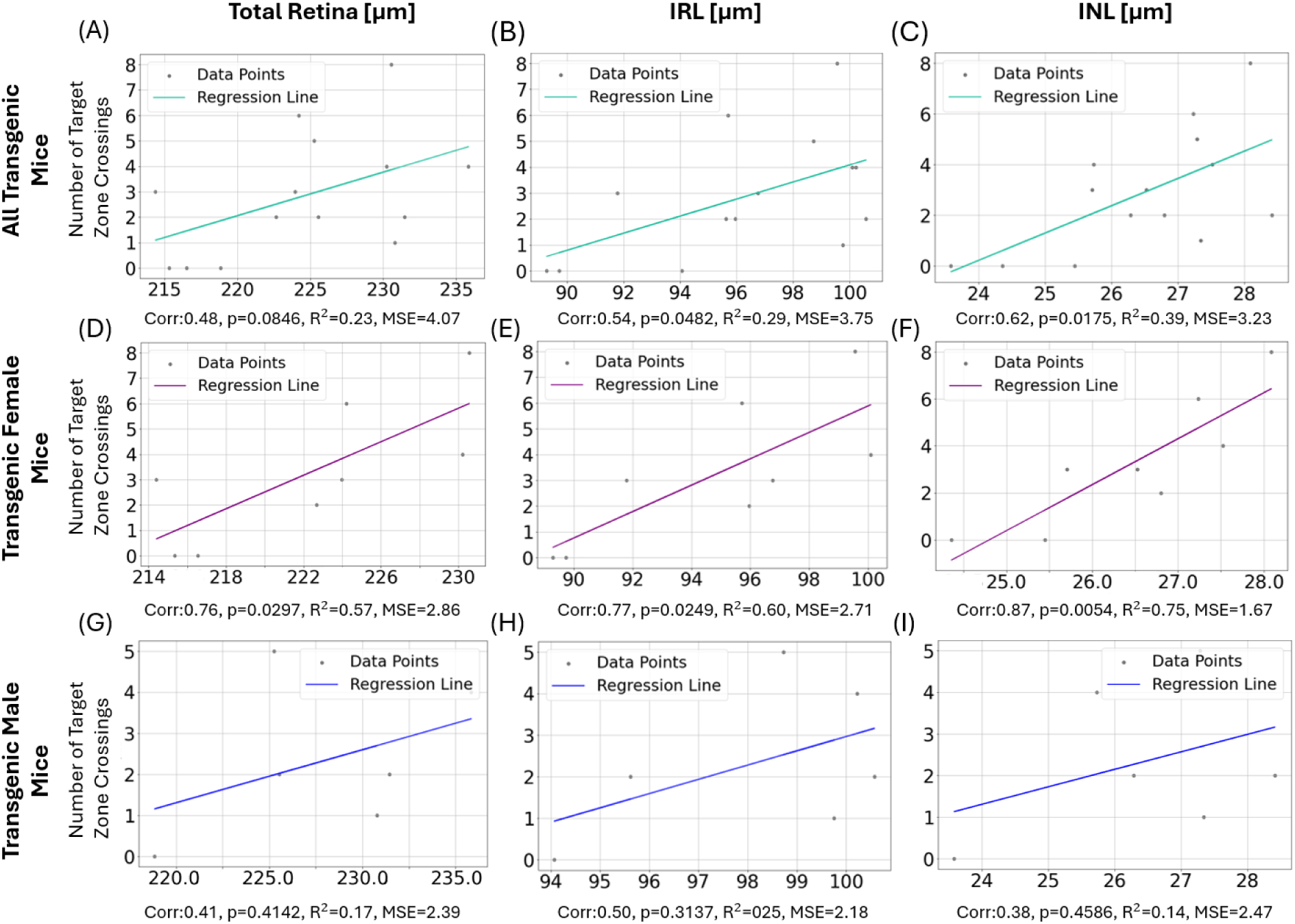
Correlation and linear regression between total retinal (left), IRL (middle) and INL thickness (right) with number of target zone crossings for all transgenic mice (A-C), female transgenic mice (D-F) and male transgenic mice (G-I). The respective Pearson correlation coefficient (Corr) and p-value as well as R^2^ and mean squared error (MSE) resulting from the linear regression are indicated below each graph. For each plot, the colored line represents the linear regression.

**Fig. 12:**
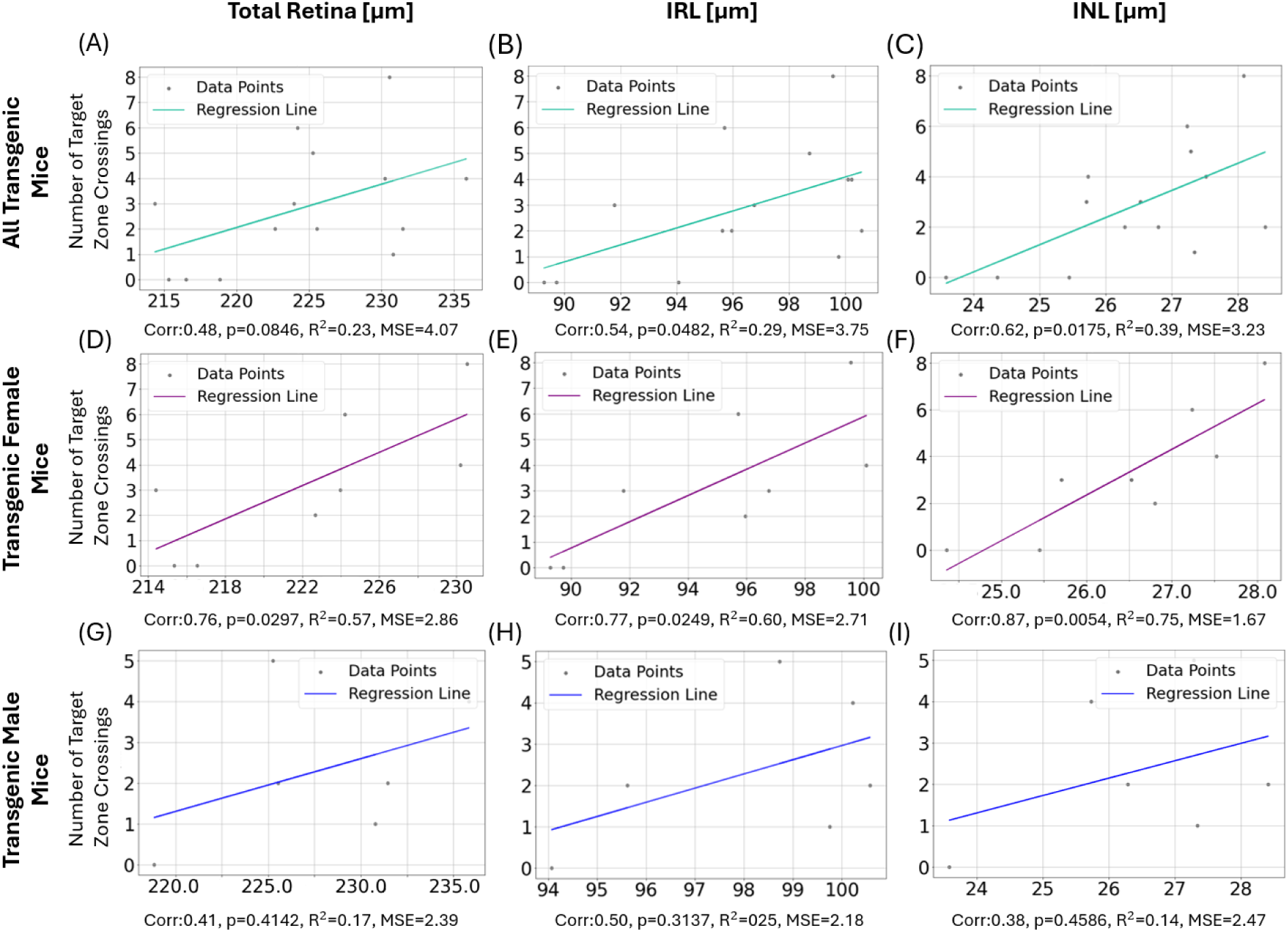
Correlation and linear regression between total retinal (left), IRL (middle) and INL thickness (right) with % of time spent in the target sector for all transgenic mice (A-C), female transgenic mice (D-F) and male transgenic mice (G-I). Below each graph, the respective Pearson correlation coefficient and p-value as well as R^2^ and MSE for the linear regression are indicated. The colored line represents the respective linear regression.

Additionally, the ICP density correlated with thigmotactic behavior for female transgenic mice (Corr=0.76, p=0.047) and the PRC thickness correlated negatively with floating behavior (Corr= -0.82, p=0.0126). For male transgenic animals, no correlations between MWM and retinal parameters were found. Furthermore, the weight of the male transgenic animals strongly correlated with the SVP density (Corr=0.96, p=0.008). Tables with all compared parameters for all animal groups (tg, tg females, tg males, ntg, ntg females and ntg males) including correlation coefficients, p-values, R^2^ and MSE are provided in the supplementary information (Table 1-12).

## 4 Discussion

In this investigation, longitudinal OCT imaging of a 5xFAD mouse model was performed in parallel to spatial memory testing and uncovered pronounced differences in retinal parameters dependent on both genotype and sex. Our analysis of retinal layer thickness revealed subtle but significant thickening of the total retina over the course of the study. Thickening was also observed for the ORL, where in particular the PRC and the OPL thickness increased over time. In the longitudinal investigation, consistent thinning was not detectable for any retinal layers from 3 months to 9 months of age. However, thickness differences were observed for group comparison at individual measurement timepoints. Our comparison between the groups at the last measurement point at 9 months of age resulted in the detection of several parameters with statistically significant differences. Here the total retina and the IRL of tg male mice was significantly thicker than for tg female mice. The IRL layers for tg male mice were also significantly thicker than for the ntg male mice indicating a swelling of the IRL for male mice. For female mice, OPL and RPE thickness significantly thin in comparison to ntg female mice at 9 months of age. The longitudinal analysis of OCT angiography data did not show changes for neither ntg nor tg female mice. However, for male ntg mice, a significant density loss in the SVP and ICP was observed over time, leading to significant differences when comparing the group to tg male mice. The observed changes in retinal layer thickness and vascular density differed strongly between male and female mice indicating a dependence of the phenotype on sex. Changes in angiographic parameters depended more on the sex of animals than on their genotype. The differences in retinal layer thickness also appeared very differently dependent on the sex of animals.

The correlation analysis of the MWM data and the retinal parameters revealed an association between the thickness of the total retina, the IRL and the INL with the number of target zone crossings (NTZC) and the time spent in the target sector (TSTS) on the test day. Both parameters for the MWM – although not completely independent of each other – are measures of spatial memory capabilities. The strongest correlation was observed between these parameters and the INL thickness of female tg mice. The correlation between the NTZC and TSTS with the IRL as well as the total retinal thickness for female mice is likely due to their strong correlation with the INL. For male tg mice, no significant positive correlation between total retina, IRL and INL layer thickness was measured. With a correlation coefficient of 0.65 for INL and TSTS, correlation between the two parameters can still be observed. Combining this information with the sex differences in layer thickness, we conclude a strong sex influence on the retinal phenotype of the 5xFAD mouse model and its connection to spatial memory.

Our results partly differ from data reported in other studies on 5xFAD mice. Lim and coworkers observed RNFL thinning and OPL thickening in the investigation of 32 tg and 38 ntg mice with unspecified sex.^24^ In contrast, we did not observe RNFL thinning, although our investigation did indicate (non-significant) thickening of the OPL. Given that the sex specifications were missing in that publication, it was not possible to perform a clear comparison of our data with this previous measurement. We do want to stress that the gender composition of the investigated animals is especially relevant given the strong impact of sex on the results measured in this study and in AD mouse models in general.^33^ The inclusion of more female or male tg or ntg control animals could shift the outcome of the measured results considerably. In another study, Matei et al. performed OCT on 16 male tg 5xFAD mice in comparison with 10 C57BL/6J and 6 C57BL/6 mice. In 3-months-old tg mice, the authors do not report changes in the total retinal thickness measured with OCT.^26^ Our investigation confirms these findings for male animals. Of note, this study did not use littermates, but C57BL/6J and C57BL/6 as controls, using two different control strains may bias the results.^26^

In their 2021 study, Kim et al. performed an OCT investigation in a female-only cohort of 5 tg and 6 ntg animals at 6 months of age.^25^ The chosen control animals were of the associated background strain (B6SJLF1/J), hence a comparison with our data on female 5xFAD mice should be straight-forward. Still, we cannot confirm their findings of retinal thinning in the total retina, the RNFL, the IRL and ORL at 6 months of age. On the other hand, our measurements confirm the absence of changes in vascular density for SVP, ICP and DCP.^25^ Given that we investigated 44 tg (22 female, n=12) and 42 ntg (19 female, n=12) volume scans for mice at 6 months of age, the lack of overlap with the previously reported results is surprising and can probably be attributed to a number of differences in the study protocols. One potential reason for this is the use of differing anaesthesia agents in the mentioned studies, as ketamine/xylazine was used by Kim et al., Lim et at., and Matei et al., whereas our study used isoflurane. ^24,25,26^ Additionally, genetic drift, breeding, body temperature of animals, anaesthesia time, blood pressure, time of imaging, eye drops as well as diet and housing conditions (e.g., single vs. group housing) could potentially – and in some cases do – differ between the studies and can influence the outcome of experiments. For this reason, we want to stress that there is a lack of standardization in the research field that makes the replication and confirmation of results extremely difficult. Without a description of every detail in the study protocol, results might considerably differ between research groups. This also applies to the comparison between mouse models. Retinal changes were reported for several other mouse models, some with partially overlapping knock-in genes (3xTg) or modelling similar disease aspects (APP/PS1 mouse for amyloid pathology) as the 5xFAD mouse model.^17^ Another factor that should be considered is the regional dependency of retinal thinning, as for example measured in 3xTg mice, where the longitudinal development of sub-layer thickness differs depending on the distance to the ONH.^34^ Given the variety in imaging protocols and aforementioned factors, as well as varying data analysis approaches, a reliable comparison of OCT findings of retinal pathologies in mouse models seems currently only possible if done in the same research group, leaving a significant knowledge gap in the investigation of the retinal phenotype of the AD disease. Since most AD mouse models only model one or few aspects of the disease pattern, comparability between studies and thus models is critical for the research of the retinal pathology of AD and the translatability of the results to human AD diagnostics.

Gender-specific medicine is crucial for drug development, diagnostics and treatment. It is therefore also increasingly important to discover sex-based differences in mouse models of disease and take these into account for preclinical studies. The study presented in this work represents not only the most extensive longitudinal OCT investigation of the 5xFAD mouse model to date, but also reveals sex-based differences in comparisons of retinal parameters with the spatial memory phenotype of the mouse models, thus adding more knowledge to further improve targeted research in the field of AD.

## 5 Conclusion

The presented study investigated retinal parameters in the 5xFAD mouse model of AD over the course of 6 months providing insights into the progression of retinal pathologies, especially the respective retinal thickness changes for male and female mice, as well as the decay in vascular density for ntg male mice. By correlating retinal parameters with the spatial memory phenotype of the investigated animals, a connection between INL thickness and MWM performance was observed for female tg animals, expanding the knowledge on the interaction between phenotypes for this particular model of AD. Our investigation proves that a multidisciplinary approach, with retinal imaging and behavior analysis, can provide more comprehensive and complementary information, which may help understanding the concurrent development of different aspects of AD. Additionally, this study shows that a separate analysis of female and male mice, as well as the minute control of experimental conditions are crucial for the generation of unbiased results in the investigation of the retinal phenotype in mouse models of AD.

## Disclosure

Magdalena Daurer, Laurenz Jauk, Roland Rabl and Manuela Prokesch are employees of Scantox Neuro GmbH. All other authors declare that there are no financial interests, commercial affiliations, or other potential conflicts of interest that could have influenced the objectivity of this research or the writing of this paper.

## Acknowledgments

The authors want to thank Andreas Hodul for his help in constructing the MWM set-up and the mouse imaging stage. We are grateful to Sonja Reynoso-De-Leon, Christian Schönauer, Jasmin Rezek and the team in the animal facility for their irreplaceable assistance with the mice. Additionally, we want to thank Patrick Bilic and Eva Fuchs for their support and guidance on animal welfare related issues, and Robin Ristl (Center for Medical Data Science, Medical University of Vienna) for assistance with the statistical analysis. Funding for this project was provided by Scantox Neuro GmbH, the FFG grant 900435, the ERC Proof of Concept grant 101069344 OPTIMEYEZ and the Austrian Science Fund grant I6092-B.

## Code, Data, and Materials Availability

All data in support of the findings of this paper are available within the article and as supplementary material. Raw data can be requested from the author at.

**Georg Ladurner** has been a PhD student at the Medical University of Vienna since March 2023. He received his BS and MS degrees in physics from the LMU Munich in 2020 and 2021, respectively. His current research interests include optical coherence tomography, Alzheimer’s disease, retinal diseases and retinal imaging techniques. He is a member of SPIE.

Biographies and photographs for the other authors are not available.

