## Supplementary Information for "Longitudinal investigation of spatial memory and retinal parameters in a 5xFAD model of Alzheimer’s disease reveals differences dependent on genotype and sex"

| Correlation Coefficient | Weight | SVP Density | ICP Density | DCP Density | Total Retina | IRL | RPE | RNFL | IPL | INL | ORL | PRC | OPL | Number of Target Zone Crossings | % of time in sector with platform | Latency on Day 3 | Latency on Day 4 | Distance swam (m) | Velocity (m/s) | Thigmotaxis % | Floating % |
|---|---|---|---|---|---|---|---|---|---|---|---|---|---|---|---|---|---|---|---|---|---|
| Weight | | 0.4691 | 0.4559 | 0.1468 | 0.5539 | 0.6622 | 0.3746 | 0.2949 | 0.2008 | 0.2965 | 0.3379 | 0.2580 | 0.4342 | 0.0662 | 0.1680 | -0.1660 | -0.3125 | -0.0326 | 0.2966 | 0.1352 | 0.1759 |
| SVP Density | 0.4691 | | -0.3331 | 0.3683 | -0.1106 | 0.0498 | -0.2922 | -0.1968 | 0.1824 | 0.3125 | -0.2897 | -0.2467 | -0.3246 | -0.0086 | 0.0656 | 0.2185 | -0.1113 | -0.0652 | 0.0637 | 0.0873 | -0.0749 |
| ICP Density | 0.4559 | -0.3331 | | -0.0330 | 0.6091 | 0.5695 | 0.5694 | 0.6591 | -0.2693 | 0.1428 | 0.5562 | 0.4435 | 0.7159 | 0.4801 | 0.3075 | -0.5473 | -0.1948 | 0.0514 | 0.3050 | 0.2867 | 0.3654 |
| DCP Density | 0.1468 | 0.3683 | -0.0330 | | -0.0457 | 0.0846 | -0.0688 | -0.4394 | 0.5011 | 0.2060 | -0.1925 | -0.1855 | -0.1299 | 0.3408 | 0.2487 | 0.6155 | 0.2802 | 0.0953 | -0.0547 | 0.0882 | -0.2940 |
| Total Retina | 0.5539 | -0.1106 | 0.6091 | -0.0457 | | 0.9390 | 0.5162 | 0.4975 | 0.1211 | 0.6429 | 0.9136 | 0.8664 | 0.6593 | 0.4771 | 0.5064 | -0.1828 | -0.3014 | -0.0879 | 0.0716 | 0.0237 | 0.1558 |
| IRL | 0.6622 | 0.0498 | 0.5695 | 0.0846 | 0.9390 | | 0.4275 | 0.3654 | 0.2993 | 0.7235 | 0.7180 | 0.6635 | 0.5713 | 0.5359 | 0.6156 | -0.1142 | -0.2953 | -0.0919 | 0.0985 | 0.0360 | 0.1632 |
| RPE | 0.3746 | -0.2922 | 0.5694 | -0.0688 | 0.5162 | 0.4275 | | 0.6623 | -0.2680 | -0.1900 | 0.5392 | 0.3279 | 0.9484 | -0.0065 | 0.0598 | -0.3602 | -0.3865 | -0.2407 | 0.0782 | -0.2154 | 0.2483 |
| RNFL | 0.2949 | -0.1968 | 0.6591 | -0.4394 | 0.4975 | 0.3654 | 0.6623 | | -0.7577 | -0.0306 | 0.5749 | 0.4524 | 0.6979 | 0.1095 | 0.1424 | -0.5604 | -0.2934 | -0.0766 | 0.1457 | 0.0294 | 0.3725 |
| IPL | 0.2008 | 0.1824 | -0.2693 | 0.5011 | 0.1211 | 0.2993 | -0.2680 | -0.7577 | | 0.3847 | -0.1088 | -0.0576 | -0.2176 | 0.1775 | 0.1858 | 0.4417 | 0.0551 | -0.0123 | -0.0563 | -0.0457 | -0.2389 |
| INL | 0.2965 | 0.3125 | 0.1428 | 0.2060 | 0.6429 | 0.7235 | -0.1900 | -0.0306 | 0.3847 | | 0.4456 | 0.5452 | -0.0520 | 0.6221 | 0.7133 | 0.2217 | -0.0257 | 0.0310 | -0.0371 | 0.1395 | -0.0632 |
| ORL | 0.3379 | -0.2897 | 0.5562 | -0.1925 | 0.9136 | 0.7180 | 0.5392 | 0.5749 | -0.1088 | 0.4456 | | 0.9690 | 0.6589 | 0.3318 | 0.2970 | -0.2349 | -0.2609 | -0.0692 | 0.0285 | 0.0054 | 0.1223 |
| PRC | 0.2580 | -0.2467 | 0.4435 | -0.1855 | 0.8664 | 0.6635 | 0.3279 | 0.4524 | -0.0576 | 0.5452 | 0.9690 | | 0.4528 | 0.3606 | 0.3130 | -0.1637 | -0.1614 | 0.0049 | 0.0116 | 0.0693 | 0.0338 |
| OPL | 0.4342 | -0.3246 | 0.7159 | -0.1299 | 0.6593 | 0.5713 | 0.9484 | 0.6979 | -0.2176 | -0.0520 | 0.6589 | 0.4528 | | 0.0995 | 0.1191 | -0.3496 | -0.4507 | -0.2649 | 0.0674 | -0.1917 | 0.3389 |
| Number of Target Zone Crossing | 0.0662 | -0.0086 | 0.4801 | 0.3408 | 0.4771 | 0.5359 | -0.0065 | 0.1095 | 0.1775 | 0.6221 | 0.3318 | 0.3606 | 0.0995 | | 0.8193 | 0.0878 | 0.2531 | 0.2756 | 0.2147 | 0.4503 | -0.1840 |
| % of time in sector with platform | 0.1680 | 0.0656 | 0.3075 | 0.2487 | 0.5064 | 0.6156 | 0.0598 | 0.1424 | 0.1858 | 0.7133 | 0.2970 | 0.3130 | 0.1191 | 0.8193 | | 0.0316 | 0.0699 | 0.1192 | 0.1138 | 0.3242 | -0.1524 |
| Latency on Day 3 | -0.1660 | 0.2185 | -0.5473 | 0.6155 | -0.1828 | -0.1142 | -0.3602 | -0.5604 | 0.4417 | 0.2217 | -0.2349 | -0.1637 | -0.3496 | 0.0878 | 0.0316 | | 0.4163 | 0.1043 | -0.2631 | -0.1299 | -0.4141 |
| Latency on Day 4 | -0.3125 | -0.1113 | -0.1948 | 0.2802 | -0.3014 | -0.2953 | -0.3865 | -0.2934 | 0.0551 | -0.0257 | -0.2609 | -0.1614 | -0.4507 | 0.2531 | 0.0699 | 0.4163 | | 0.8990 | 0.5430 | 0.6638 | -0.8409 |
| Distance swam (m) | -0.0326 | -0.0652 | 0.0514 | 0.0953 | -0.0879 | -0.0919 | -0.2407 | -0.0766 | -0.0123 | 0.0310 | -0.0692 | 0.0049 | -0.2649 | 0.2756 | 0.1192 | 0.1043 | 0.8990 | | 0.8338 | 0.8775 | -0.8141 |
| Velocity (m/s) | 0.2966 | 0.0637 | 0.3050 | -0.0547 | 0.0716 | 0.0985 | 0.0782 | 0.1457 | -0.0563 | -0.0371 | 0.0285 | 0.0116 | 0.0674 | 0.2147 | 0.1138 | -0.2631 | 0.5430 | 0.8338 | | 0.8738 | -0.5950 |
| Thigmotaxis % | 0.1352 | 0.0873 | 0.2867 | 0.0882 | 0.0237 | 0.0360 | -0.2154 | 0.0294 | -0.0457 | 0.1395 | 0.0054 | 0.0693 | -0.1917 | 0.4503 | 0.3242 | -0.1299 | 0.6638 | 0.8775 | 0.8738 | | -0.6370 |
| Floating % | 0.1759 | -0.0749 | 0.3654 | -0.2940 | 0.1558 | 0.1632 | 0.2483 | 0.3725 | -0.2389 | -0.0632 | 0.1223 | 0.0338 | 0.3389 | -0.1840 | -0.1524 | -0.4141 | -0.8409 | -0.8141 | -0.5950 | -0.6370 | |

| P-Value | Weight | SVP Density | ICP Density | DCP Density | Total Retina | IRL | RPE | RNFL | IPL | INL | ORL | PRC | OPL | Number of Target Zone Crossings | % of time in sector with platform | Latency on Day 3 | Latency on Day 4 | Distance swam (m) | Velocity (m/s) | Thigmotaxis % | Floating % |
|---|---|---|---|---|---|---|---|---|---|---|---|---|---|---|---|---|---|---|---|---|---|
| Weight | | 0.1058 | 0.1364 | 0.6166 | 0.0399 | 0.0099 | 0.1869 | 0.3061 | 0.4912 | 0.3033 | 0.2374 | 0.3731 | 0.1209 | 0.8221 | 0.5658 | 0.5706 | 0.2766 | 0.9118 | 0.3031 | 0.6450 | 0.5475 |
| SVP Density | 0.1058 | | 0.2900 | 0.2156 | 0.7192 | 0.8717 | 0.3327 | 0.5193 | 0.5510 | 0.2985 | 0.3370 | 0.4165 | 0.2791 | 0.9779 | 0.8313 | 0.4733 | 0.7174 | 0.8324 | 0.8361 | 0.7766 | 0.8080 |
| ICP Density | 0.1364 | 0.2900 | | 0.9189 | 0.0355 | 0.0532 | 0.0533 | 0.0197 | 0.3972 | 0.6580 | 0.0604 | 0.1487 | 0.0088 | 0.1142 | 0.3310 | 0.0655 | 0.5441 | 0.8739 | 0.3351 | 0.3662 | 0.2428 |
| DCP Density | 0.6166 | 0.2156 | 0.9189 | | 0.8768 | 0.7737 | 0.8152 | 0.1159 | 0.0680 | 0.4799 | 0.5097 | 0.5254 | 0.6579 | 0.2330 | 0.3912 | 0.0191 | 0.3320 | 0.7458 | 0.8527 | 0.7643 | 0.3077 |
| Total Retina | 0.0399 | 0.7192 | 0.0355 | 0.8768 | | 0.0000 | 0.0588 | 0.0703 | 0.6800 | 0.0131 | 0.0000 | 0.0001 | 0.0103 | 0.0846 | 0.0646 | 0.5317 | 0.2949 | 0.7652 | 0.8078 | 0.9360 | 0.5947 |
| IRL | 0.0099 | 0.8717 | 0.0532 | 0.7737 | 0.0000 | | 0.1273 | 0.1990 | 0.2985 | 0.0034 | 0.0038 | 0.0097 | 0.0328 | 0.0482 | 0.0191 | 0.6975 | 0.3054 | 0.7548 | 0.7377 | 0.9029 | 0.5771 |
| RPE | 0.1869 | 0.3327 | 0.0533 | 0.8152 | 0.0588 | 0.1273 | | 0.0099 | 0.3542 | 0.5152 | 0.0466 | 0.2525 | 0.0000 | 0.9823 | 0.8391 | 0.2058 | 0.1722 | 0.4070 | 0.7904 | 0.4595 | 0.3920 |
| RNFL | 0.3061 | 0.5193 | 0.0197 | 0.1159 | 0.0703 | 0.1990 | 0.0099 | | 0.0017 | 0.9172 | 0.0315 | 0.1044 | 0.0055 | 0.7093 | 0.6273 | 0.0371 | 0.3087 | 0.7945 | 0.6193 | 0.9206 | 0.1896 |
| IPL | 0.4912 | 0.5510 | 0.3972 | 0.0680 | 0.6800 | 0.2985 | 0.3542 | 0.0017 | | 0.1744 | 0.7111 | 0.8450 | 0.4548 | 0.5438 | 0.5247 | 0.1139 | 0.8516 | 0.9666 | 0.8484 | 0.8768 | 0.4107 |
| INL | 0.3033 | 0.2985 | 0.6580 | 0.4799 | 0.0131 | 0.0034 | 0.5152 | 0.9172 | 0.1744 | | 0.1103 | 0.0438 | 0.8598 | 0.0175 | 0.0042 | 0.4463 | 0.9304 | 0.9162 | 0.8998 | 0.6344 | 0.8301 |
| ORL | 0.2374 | 0.3370 | 0.0604 | 0.5097 | 0.0000 | 0.0038 | 0.0466 | 0.0315 | 0.7111 | 0.1103 | | 0.0000 | 0.0104 | 0.2465 | 0.3025 | 0.4190 | 0.3676 | 0.8142 | 0.9230 | 0.9854 | 0.6769 |
| PRC | 0.3731 | 0.4165 | 0.1487 | 0.5254 | 0.0001 | 0.0097 | 0.2525 | 0.1044 | 0.8450 | 0.0438 | 0.0000 | | 0.1040 | 0.2053 | 0.2760 | 0.5761 | 0.5816 | 0.9867 | 0.9685 | 0.8139 | 0.9087 |
| OPL | 0.1209 | 0.2791 | 0.0088 | 0.6579 | 0.0103 | 0.0328 | 0.0000 | 0.0055 | 0.4548 | 0.8598 | 0.0104 | 0.1040 | | 0.7349 | 0.6852 | 0.2205 | 0.1058 | 0.3601 | 0.8188 | 0.5116 | 0.2360 |
| Number of Target Zone Crossing | 0.8221 | 0.9779 | 0.1142 | 0.2330 | 0.0846 | 0.0482 | 0.9823 | 0.7093 | 0.5438 | 0.0175 | 0.2465 | 0.2053 | 0.7349 | | 0.0003 | 0.7653 | 0.3826 | 0.3401 | 0.4610 | 0.1062 | 0.5288 |
| % of time in sector with platform | 0.5658 | 0.8313 | 0.3310 | 0.3912 | 0.0646 | 0.0191 | 0.8391 | 0.6273 | 0.5247 | 0.0042 | 0.3025 | 0.2760 | 0.6852 | 0.0003 | | 0.9145 | 0.8124 | 0.6848 | 0.6984 | 0.2581 | 0.6029 |
| Latency on Day 3 | 0.5706 | 0.4733 | 0.0655 | 0.0191 | 0.5317 | 0.6975 | 0.2058 | 0.0371 | 0.1139 | 0.4463 | 0.4190 | 0.5761 | 0.2205 | 0.7653 | 0.9145 | | 0.1387 | 0.7228 | 0.3634 | 0.6581 | 0.1411 |
| Latency on Day 4 | 0.2766 | 0.7174 | 0.5441 | 0.3320 | 0.2949 | 0.3054 | 0.1722 | 0.3087 | 0.8516 | 0.9304 | 0.3676 | 0.5816 | 0.1058 | 0.3826 | 0.8124 | 0.1387 | | 0.0000 | 0.0448 | 0.0096 | 0.0002 |
| Distance swam (m) | 0.9118 | 0.8324 | 0.8739 | 0.7458 | 0.7652 | 0.7548 | 0.4070 | 0.7945 | 0.9666 | 0.9162 | 0.8142 | 0.9867 | 0.3601 | 0.3401 | 0.6848 | 0.7228 | 0.0000 | | 0.0002 | 0.0000 | 0.0004 |
| Velocity (m/s) | 0.3031 | 0.8361 | 0.3351 | 0.8527 | 0.8078 | 0.7377 | 0.7904 | 0.6193 | 0.8484 | 0.8998 | 0.9230 | 0.9685 | 0.8188 | 0.4610 | 0.6984 | 0.3634 | 0.0448 | 0.0002 | | 0.0000 | 0.0248 |
| Thigmotaxis % | 0.6450 | 0.7766 | 0.3662 | 0.7643 | 0.9360 | 0.9029 | 0.4595 | 0.9206 | 0.8768 | 0.6344 | 0.9854 | 0.8139 | 0.5116 | 0.1062 | 0.2581 | 0.6581 | 0.0096 | 0.0000 | 0.0000 | | 0.0143 |
| Floating % | 0.5475 | 0.8080 | 0.2428 | 0.3077 | 0.5947 | 0.5771 | 0.3920 | 0.1896 | 0.4107 | 0.8301 | 0.6769 | 0.9087 | 0.2360 | 0.5288 | 0.6029 | 0.1411 | 0.0002 | 0.0004 | 0.0248 | 0.0143 | |

Table 1: Correlation Coefficient and corresponding p-value for or all tested parameters for all transgenic animals. For the Correlation coefficient red indicates values from 0-0.3, yellow =0.3-0.5 and green 0.5-1. For the p-value red indicates values above 0.15, yellow values from 0.15-0.1 and green values from below 0.05.

| R-Squared | Weight | SVP Density | ICP Density | DCP Density | Total Retina | IRL | RPE | RNFL | IPL | INL | ORL | PRC | OPL | Number of Target Zone Crossings | % of time in sector with platform | Latency on Day 3 | Latency on Day 4 | Distance swam (m) | Velocity (m/s) | Thigmotaxis % | Floating % |
|---|---|---|---|---|---|---|---|---|---|---|---|---|---|---|---|---|---|---|---|---|---|
| Weight | | 0.2201 | 0.2078 | 0.0215 | 0.3068 | 0.4386 | 0.1403 | 0.0870 | 0.0403 | 0.0879 | 0.1142 | 0.0666 | 0.1885 | 0.0044 | 0.0282 | 0.0276 | 0.0977 | 0.0011 | 0.0880 | 0.0183 | 0.0309 |
| SVP Density | 0.2201 | | 0.1110 | 0.1357 | 0.0122 | 0.0025 | 0.0854 | 0.0387 | 0.0333 | 0.0977 | 0.0839 | 0.0609 | 0.1054 | 0.0001 | 0.0043 | 0.0477 | 0.0124 | 0.0043 | 0.0041 | 0.0076 | 0.0056 |
| ICP Density | 0.2078 | 0.1110 | | 0.0011 | 0.3710 | 0.3244 | 0.3242 | 0.4344 | 0.0725 | 0.0204 | 0.3093 | 0.1967 | 0.5125 | 0.2305 | 0.0945 | 0.2995 | 0.0379 | 0.0026 | 0.0930 | 0.0822 | 0.1335 |
| DCP Density | 0.0215 | 0.1357 | 0.0011 | | 0.0021 | 0.0072 | 0.0047 | 0.1931 | 0.2511 | 0.0424 | 0.0371 | 0.0344 | 0.0169 | 0.1162 | 0.0619 | 0.3788 | 0.0785 | 0.0091 | 0.0030 | 0.0078 | 0.0864 |
| Total Retina | 0.3068 | 0.0122 | 0.3710 | 0.0021 | | 0.8817 | 0.2665 | 0.2475 | 0.0147 | 0.4133 | 0.8346 | 0.7507 | 0.4347 | 0.2276 | 0.2564 | 0.0334 | 0.0909 | 0.0077 | 0.0051 | 0.0006 | 0.0243 |
| IRL | 0.4386 | 0.0025 | 0.3244 | 0.0072 | 0.8817 | | 0.1828 | 0.1335 | 0.0896 | 0.5235 | 0.5155 | 0.4402 | 0.3264 | 0.2872 | 0.3789 | 0.0130 | 0.0872 | 0.0084 | 0.0097 | 0.0013 | 0.0267 |
| RPE | 0.1403 | 0.0854 | 0.3242 | 0.0047 | 0.2665 | 0.1828 | | 0.4387 | 0.0718 | 0.0361 | 0.2907 | 0.1075 | 0.8994 | 0.0000 | 0.0036 | 0.1298 | 0.1494 | 0.0580 | 0.0061 | 0.0464 | 0.0617 |
| RNFL | 0.0870 | 0.0387 | 0.4344 | 0.1931 | 0.2475 | 0.1335 | 0.4387 | | 0.5741 | 0.0009 | 0.3305 | 0.2046 | 0.4871 | 0.0120 | 0.0203 | 0.3140 | 0.0861 | 0.0059 | 0.0212 | 0.0009 | 0.1388 |
| IPL | 0.0403 | 0.0333 | 0.0725 | 0.2511 | 0.0147 | 0.0896 | 0.0718 | 0.5741 | | 0.1480 | 0.0118 | 0.0033 | 0.0474 | 0.0315 | 0.0345 | 0.1951 | 0.0030 | 0.0002 | 0.0032 | 0.0021 | 0.0571 |
| INL | 0.0879 | 0.0977 | 0.0204 | 0.0424 | 0.4133 | 0.5235 | 0.0361 | 0.0009 | 0.1480 | | 0.1985 | 0.2972 | 0.0027 | 0.3871 | 0.5088 | 0.0491 | 0.0007 | 0.0010 | 0.0014 | 0.0195 | 0.0040 |
| ORL | 0.1142 | 0.0839 | 0.3093 | 0.0371 | 0.8346 | 0.5155 | 0.2907 | 0.3305 | 0.0118 | 0.1985 | | 0.9391 | 0.4341 | 0.1101 | 0.0882 | 0.0552 | 0.0681 | 0.0048 | 0.0008 | 0.0000 | 0.0150 |
| PRC | 0.0666 | 0.0609 | 0.1967 | 0.0344 | 0.7507 | 0.4402 | 0.1075 | 0.2046 | 0.0033 | 0.2972 | 0.9391 | | 0.2050 | 0.1300 | 0.0979 | 0.0268 | 0.0260 | 0.0000 | 0.0001 | 0.0048 | 0.0011 |
| OPL | 0.1885 | 0.1054 | 0.5125 | 0.0169 | 0.4347 | 0.3264 | 0.8994 | 0.4871 | 0.0474 | 0.0027 | 0.4341 | 0.2050 | | 0.0099 | 0.0142 | 0.1222 | 0.2031 | 0.0702 | 0.0045 | 0.0367 | 0.1148 |
| Number of Target Zone Crossing | 0.0044 | 0.0001 | 0.2305 | 0.1162 | 0.2276 | 0.2872 | 0.0000 | 0.0120 | 0.0315 | 0.3871 | 0.1101 | 0.1300 | 0.0099 | | 0.6712 | 0.0077 | 0.0641 | 0.0760 | 0.0461 | 0.2027 | 0.0339 |
| % of time in sector with platform | 0.0282 | 0.0043 | 0.0945 | 0.0619 | 0.2564 | 0.3789 | 0.0036 | 0.0203 | 0.0345 | 0.5088 | 0.0882 | 0.0979 | 0.0142 | 0.6712 | | 0.0010 | 0.0049 | 0.0142 | 0.0130 | 0.1051 | 0.0232 |
| Latency on Day 3 | 0.0276 | 0.0477 | 0.2995 | 0.3788 | 0.0334 | 0.0130 | 0.1298 | 0.3140 | 0.1951 | 0.0491 | 0.0552 | 0.0268 | 0.1222 | 0.0077 | 0.0010 | | 0.1733 | 0.0109 | 0.0692 | 0.0169 | 0.1715 |
| Latency on Day 4 | 0.0977 | 0.0124 | 0.0379 | 0.0785 | 0.0909 | 0.0872 | 0.1494 | 0.0861 | 0.0030 | 0.0007 | 0.0681 | 0.0260 | 0.2031 | 0.0641 | 0.0049 | 0.1733 | | 0.8081 | 0.2948 | 0.4407 | 0.7071 |
| Distance swam (m) | 0.0011 | 0.0043 | 0.0026 | 0.0091 | 0.0077 | 0.0084 | 0.0580 | 0.0059 | 0.0002 | 0.0010 | 0.0048 | 0.0000 | 0.0702 | 0.0760 | 0.0142 | 0.0109 | 0.8081 | | 0.6953 | 0.7700 | 0.6627 |
| Velocity (m/s) | 0.0880 | 0.0041 | 0.0930 | 0.0030 | 0.0051 | 0.0097 | 0.0061 | 0.0212 | 0.0032 | 0.0014 | 0.0008 | 0.0001 | 0.0045 | 0.0461 | 0.0130 | 0.0692 | 0.2948 | 0.6953 | | 0.7636 | 0.3540 |
| Thigmotaxis % | 0.0183 | 0.0076 | 0.0822 | 0.0078 | 0.0006 | 0.0013 | 0.0464 | 0.0009 | 0.0021 | 0.0195 | 0.0000 | 0.0048 | 0.0367 | 0.2027 | 0.1051 | 0.0169 | 0.4407 | 0.7700 | 0.7636 | | 0.4057 |
| Floating % | 0.0309 | 0.0056 | 0.1335 | 0.0864 | 0.0243 | 0.0267 | 0.0617 | 0.1388 | 0.0571 | 0.0040 | 0.0150 | 0.0011 | 0.1148 | 0.0339 | 0.0232 | 0.1715 | 0.7071 | 0.6627 | 0.3540 | 0.4057 | |

| Mean Squared Error | Weight | SVP Density | ICP Density | DCP Density | Total Retina | IRL | RPE | RNFL | IPL | INL | ORL | PRC | OPL | Number of Target Zone Crossings | % of time in sector with platform | Latency on Day 3 | Latency on Day 4 | Distance swam (m) | Velocity (m/s) | Thigmotaxis % | Floating % |
|---|---|---|---|---|---|---|---|---|---|---|---|---|---|---|---|---|---|---|---|---|---|
| Weight | | 0.000109 | 0.000221 | 0.000223 | 28.33047001 | 7.8337661 | 0.4083889 | 17.523208 | 14.801248 | 1.6035146 | 8.8361361 | 6.6275976 | 0.5522975 | 5.242241001 | 522.1335214 | 181.6059109 | 196.7953113 | 111640.432 | 10.805108 | 582.464451 | 56.68741 |
| SVP Density | 0.0001086 | | 0.000248 | 0.000212 | 40.69152267 | 14.589362 | 0.426639 | 19.848042 | 15.947988 | 1.0933545 | 8.8397616 | 5.7669023 | 0.6293698 | 4.993716707 | 560.995439 | 178.571698 | 230.572394 | 119842.1416 | 12.078296 | 633.6155311 | 62.28985 |
| ICP Density | 0.0002207 | 0.000248 | | 0.000266 | 26.60232438 | 9.8911133 | 0.336993 | 11.920597 | 12.937778 | 1.2222928 | 7.0924075 | 5.3114027 | 0.3489194 | 4.10402001 | 537.2653195 | 111.4725105 | 241.6857196 | 130013.5723 | 11.899779 | 623.0352754 | 56.975535 |
| DCP Density | 0.0002234 | 0.000212 | 0.000266 | | 40.7837621 | 13.85341 | 0.4728084 | 15.486448 | 11.550476 | 1.6834773 | 9.6054183 | 6.8559301 | 0.6690901 | 4.653591305 | 504.0703405 | 116.008315 | 200.981531 | 110743.4323 | 11.812237 | 588.6869658 | 53.442391 |
| Total Retina | 28.33047 | 40.69152 | 26.60232 | 40.78376 | | 1.6501146 | 0.3484704 | 14.44214 | 15.197019 | 1.0314674 | 1.6501146 | 1.7699878 | 0.3847396 | 4.06704223 | 399.5141468 | 180.5145242 | 198.2830746 | 110896.824 | 11.786889 | 592.9710936 | 57.076885 |
| IRL | 7.8337661 | 14.58936 | 9.891113 | 13.85341 | 1.650114591 | | 0.3882219 | 16.630455 | 14.04148 | 0.8377632 | 4.8331766 | 3.9744671 | 0.4584688 | 3.753161458 | 333.7106177 | 184.3164611 | 199.0855728 | 110816.4459 | 11.732773 | 592.5363362 | 56.938432 |
| RPE | 0.4083889 | 0.426639 | 0.336993 | 0.472808 | 0.348470354 | 0.3882219 | | 10.773578 | 14.31551 | 1.6945804 | 7.0748756 | 6.3370349 | 0.0684525 | 5.26508214 | 535.3845135 | 162.5184306 | 185.5180549 | 105282.017 | 11.775186 | 565.7697742 | 54.890654 |
| RNFL | 17.523208 | 19.84804 | 11.9206 | 15.48645 | 14.44213978 | 16.630455 | 10.773578 | | 6.5693994 | 1.7564137 | 6.6783269 | 5.6474363 | 0.349049 | 5.202139641 | 526.4128989 | 128.1029067 | 199.3291587 | 111102.9297 | 11.596236 | 592.7916048 | 50.380598 |
| IPL | 14.801248 | 15.94799 | 12.93778 | 11.55048 | 15.19701889 | 14.04148 | 14.31551 | 6.5693994 | | 1.4978586 | 9.8569421 | 7.0768017 | 0.6483465 | 5.099432739 | 518.7508196 | 150.3213503 | 217.4381782 | 111742.3735 | 11.810086 | 592.0668773 | 55.158252 |
| INL | 1.6035146 | 1.093355 | 1.222293 | 1.683477 | 1.031467414 | 0.8377632 | 1.6945804 | 1.7564137 | 1.4978586 | | 7.9947991 | 4.9899343 | 0.6787405 | 3.22726988 | 263.9114308 | 177.5742545 | 217.9559385 | 111651.9379 | 11.831331 | 581.7611105 | 58.263854 |
| ORL | 8.8361361 | 8.839762 | 7.092407 | 9.605418 | 1.650114605 | 4.8331766 | 7.0748756 | 6.6783269 | 9.8569421 | 7.9947991 | | 0.432734 | 0.3851316 | 4.685658183 | 489.9101649 | 176.4510124 | 203.2516101 | 111224.4127 | 11.838031 | 593.2864127 | 57.621778 |
| PRC | 6.6275976 | 5.766902 | 5.311403 | 6.85593 | 1.769987756 | 3.9744671 | 6.3370349 | 5.6474363 | 7.0768017 | 4.9899343 | 0.432734 | | 0.5410623 | 4.580638482 | 484.6818465 | 181.750216 | 212.4218537 | 111756.6966 | 11.846045 | 590.4553656 | 58.430547 |
| OPL | 0.5522975 | 0.62937 | 0.348919 | 0.66909 | 0.384739607 | 0.4584688 | 0.0684525 | 0.349049 | 0.6483465 | 0.6787405 | 0.3851316 | 0.5410623 | | 5.213135863 | 529.6888359 | 163.9320019 | 173.7966516 | 103918.966 | 11.793778 | 571.5077459 | 51.780376 |
| Number of Target Zone Crossing | 5.242241 | 4.993717 | 4.10402 | 4.653591 | 4.06704223 | 3.7531615 | 5.2650821 | 5.2021396 | 5.0994327 | 3.2272699 | 4.6856582 | 4.5806385 | 5.2131359 | | 176.6528912 | 185.3112828 | 204.1249067 | 103267.6684 | 11.301365 | 473.0206523 | 56.516184 |
| % of time in sector with platform | 522.13352 | 560.9954 | 537.2653 | 504.0703 | 399.5141468 | 333.71062 | 535.38451 | 526.4129 | 518.75082 | 263.91143 | 489.91016 | 484.68185 | 529.68884 | 176.6528912 | | 186.5650406 | 217.0363155 | 110171.2219 | 11.69418 | 530.9375604 | 57.138521 |
| Latency on Day 3 | 181.60591 | 178.5717 | 111.4725 | 116.0083 | 180.5145242 | 184.31646 | 162.51843 | 128.10291 | 150.32135 | 177.57425 | 176.45101 | 181.75022 | 163.932 | 185.3112828 | 186.5650406 | | 180.3057746 | 110544.7872 | 11.02726 | 583.2954033 | 48.467994 |
| Latency on Day 4 | 196.79531 | 230.5724 | 241.6857 | 200.9815 | 198.2830746 | 199.08557 | 185.51805 | 199.32916 | 217.43818 | 217.95594 | 203.25161 | 212.42185 | 173.79665 | 204.1249067 | 217.0363155 | 180.3057746 | | 21441.46272 | 8.3549825 | 331.8634129 | 17.135592 |
| Distance swam (m) | 111640.43 | 119842.1 | 130013.6 | 110743.4 | 110896.824 | 110816.45 | 105282.02 | 111102.93 | 111742.37 | 111651.94 | 111224.41 | 111756.7 | 103918.97 | 103267.6684 | 110171.2219 | 110544.7872 | 21441.46272 | | 3.6103339 | 136.4316766 | 19.730773 |
| Velocity (m/s) | 10.805108 | 12.0783 | 11.89978 | 11.81224 | 11.78688873 | 11.732773 | 11.775186 | 11.596236 | 11.810086 | 11.831331 | 11.838031 | 11.846045 | 11.793778 | 11.30136511 | 11.69418023 | 11.02726003 | 8.354982496 | 3.610333879 | | 140.2529121 | 37.787632 |
| Thigmotaxis % | 582.46445 | 633.6155 | 623.0353 | 588.687 | 592.9710936 | 592.53634 | 565.76977 | 592.7916 | 592.06688 | 581.76111 | 593.28641 | 590.45537 | 571.50775 | 473.0206523 | 530.9375604 | 583.2954033 | 331.8634129 | 136.4316766 | 140.25291 | | 34.764487 |
| Floating % | 56.68741 | 62.28985 | 56.97553 | 53.44239 | 57.07688479 | 56.938432 | 54.890654 | 50.380598 | 55.158252 | 58.263854 | 57.621778 | 58.430547 | 51.780376 | 56.51618371 | 57.13852079 | 48.46799435 | 17.1355921 | 19.73077285 | 37.787632 | 34.76448702 | |

Table 2: R-squared and corresponding MSE for or all tested parameters for all transgenic animals. For the R-squared values red indicates values from 0-0.3, yellow =0.4-0.6 and green 0.6-1.

| Correlation Coefficient | Weight | SVP Density | ICP Density | DCP Density | Total Retina | IRL | RPE | RNFL | IPL | INL | ORL | PRC | OPL | Number of Target Zone Crossings | % of time in sector with platform | Latency on Day 3 | Latency on Day 4 | Distance swam (m) | Velocity (m/s) | Thigmotaxis % | Floating % |
|---|---|---|---|---|---|---|---|---|---|---|---|---|---|---|---|---|---|---|---|---|---|
| Weight | | -0.5352 | 0.5715 | 0.5770 | 0.6478 | 0.7642 | 0.2464 | -0.1911 | 0.6432 | 0.6528 | 0.3549 | 0.2916 | 0.4125 | 0.5264 | 0.3967 | 0.3810 | 0.4841 | 0.4328 | 0.2766 | 0.2089 | -0.2112 |
| SVP Density | -0.5352 | | -0.7469 | 0.2163 | -0.4732 | -0.5541 | -0.7167 | -0.2330 | -0.1754 | -0.3341 | -0.2659 | -0.0736 | -0.8647 | -0.2266 | -0.3171 | -0.1850 | 0.0069 | -0.0948 | -0.2308 | 0.0176 | -0.1153 |
| ICP Density | 0.5715 | -0.7469 | | 0.1723 | 0.4042 | 0.5150 | 0.0413 | 0.4730 | -0.1680 | 0.4729 | 0.1519 | 0.0792 | 0.4412 | 0.6762 | 0.6046 | -0.3230 | 0.4516 | 0.6341 | 0.6650 | 0.7603 | 0.0350 |
| DCP Density | 0.5770 | 0.2163 | 0.1723 | | 0.0665 | 0.1386 | -0.6039 | -0.6842 | 0.6425 | 0.2913 | -0.0601 | 0.0510 | -0.4545 | 0.2489 | 0.0860 | 0.5218 | 0.5099 | 0.2882 | -0.0537 | 0.2386 | -0.3164 |
| Total Retina | 0.6478 | -0.4732 | 0.4042 | 0.0665 | | 0.9616 | 0.5047 | 0.0921 | 0.5004 | 0.8953 | 0.8977 | 0.8365 | 0.6655 | 0.7568 | 0.6931 | 0.0293 | 0.2459 | 0.4256 | 0.4775 | 0.3599 | -0.5005 |
| IRL | 0.7642 | -0.5541 | 0.5150 | 0.1386 | 0.9616 | | 0.5303 | 0.0667 | 0.5472 | 0.9254 | 0.7423 | 0.6636 | 0.6576 | 0.7716 | 0.7403 | -0.0103 | 0.1745 | 0.3319 | 0.3984 | 0.2855 | -0.2923 |
| RPE | 0.2464 | -0.7167 | 0.0413 | -0.6039 | 0.5047 | 0.5303 | | 0.3945 | 0.0358 | 0.2461 | 0.3810 | 0.1930 | 0.9031 | 0.0227 | 0.1246 | -0.1941 | -0.3027 | -0.1001 | 0.2050 | -0.2916 | 0.1554 |
| RNFL | -0.1911 | -0.2330 | 0.4730 | -0.6842 | 0.0921 | 0.0667 | 0.3945 | | -0.7945 | -0.1009 | 0.1178 | 0.0231 | 0.4193 | 0.2131 | 0.1912 | -0.5477 | 0.0972 | 0.2914 | 0.4266 | 0.2891 | 0.2473 |
| IPL | 0.6432 | -0.1754 | -0.1680 | 0.6425 | 0.5004 | 0.5472 | 0.0358 | -0.7945 | | 0.6135 | 0.3432 | 0.3648 | 0.0819 | 0.2438 | 0.2382 | 0.4899 | 0.0316 | -0.0496 | -0.1296 | -0.1185 | -0.3738 |
| INL | 0.6528 | -0.3341 | 0.4729 | 0.2913 | 0.8953 | 0.9254 | 0.2461 | -0.1009 | 0.6135 | | 0.7003 | 0.6832 | 0.4017 | 0.8660 | 0.8610 | -0.0797 | 0.1166 | 0.2899 | 0.3592 | 0.4224 | -0.3501 |
| ORL | 0.3549 | -0.2659 | 0.1519 | -0.0601 | 0.8977 | 0.7423 | 0.3810 | 0.1178 | 0.3432 | 0.7003 | | 0.9767 | 0.5691 | 0.6091 | 0.5038 | 0.0881 | 0.3203 | 0.5062 | 0.5261 | 0.4204 | -0.7526 |
| PRC | 0.2916 | -0.0736 | 0.0792 | 0.0510 | 0.8365 | 0.6636 | 0.1930 | 0.0231 | 0.3648 | 0.6832 | 0.9767 | | 0.3794 | 0.6184 | 0.4888 | 0.1098 | 0.3615 | 0.5101 | 0.4711 | 0.4581 | -0.8203 |
| OPL | 0.4125 | -0.8647 | 0.4412 | -0.4545 | 0.6655 | 0.6576 | 0.9031 | 0.4193 | 0.0819 | 0.4017 | 0.5691 | 0.3794 | | 0.2565 | 0.2995 | -0.0407 | -0.0044 | 0.2278 | 0.4633 | 0.0570 | -0.1016 |
| Number of Target Zone Crossings | 0.5264 | -0.2266 | 0.6762 | 0.2489 | 0.7568 | 0.7716 | 0.0227 | 0.2131 | 0.2438 | 0.8660 | 0.6091 | 0.6184 | 0.2565 | | 0.9371 | -0.1516 | 0.3414 | 0.4746 | 0.4227 | 0.6626 | -0.2902 |
| % of time in sector with platform | 0.3967 | -0.3171 | 0.6046 | 0.0860 | 0.6931 | 0.7403 | 0.1246 | 0.1912 | 0.2382 | 0.8610 | 0.5038 | 0.4888 | 0.2995 | 0.9371 | | -0.2771 | 0.0147 | 0.1914 | 0.2709 | 0.5050 | -0.0793 |
| Latency on Day 3 | 0.3810 | -0.1850 | -0.3230 | 0.5218 | 0.0293 | -0.0103 | -0.1941 | -0.5477 | 0.4899 | -0.0797 | 0.0881 | 0.1098 | -0.0407 | -0.1516 | -0.2771 | | 0.4560 | 0.1149 | -0.2940 | -0.1959 | -0.4057 |
| Latency on Day 4 | 0.4841 | 0.0069 | 0.4516 | 0.5099 | 0.2459 | 0.1745 | -0.3027 | 0.0972 | 0.0316 | 0.1166 | 0.3203 | 0.3615 | -0.0044 | 0.3414 | 0.0147 | 0.4560 | | 0.8865 | 0.4994 | 0.6078 | -0.5941 |
| Distance swam (m) | 0.4328 | -0.0948 | 0.6341 | 0.2882 | 0.4256 | 0.3319 | -0.1001 | 0.2914 | -0.0496 | 0.2899 | 0.5062 | 0.5101 | 0.2278 | 0.4746 | 0.1914 | 0.1149 | 0.8865 | | 0.8399 | 0.8298 | -0.6491 |
| Velocity (m/s) | 0.2766 | -0.2308 | 0.6650 | -0.0537 | 0.4775 | 0.3984 | 0.2050 | 0.4266 | -0.1296 | 0.3592 | 0.5261 | 0.4711 | 0.4633 | 0.4227 | 0.2709 | -0.2940 | 0.4994 | 0.8399 | | 0.8065 | -0.4891 |
| Thigmotaxis % | 0.2089 | 0.0176 | 0.7603 | 0.2386 | 0.3599 | 0.2855 | -0.2916 | 0.2891 | -0.1185 | 0.4224 | 0.4204 | 0.4581 | 0.0570 | 0.6626 | 0.5050 | -0.1959 | 0.6078 | 0.8298 | 0.8065 | | -0.5092 |
| Floating % | -0.2112 | -0.1153 | 0.0350 | -0.3164 | -0.5005 | -0.2923 | 0.1554 | 0.2473 | -0.3738 | -0.3501 | -0.7526 | -0.8203 | -0.1016 | -0.2902 | -0.0793 | -0.4057 | -0.5941 | -0.6491 | -0.4891 | -0.5092 | |

| P-Value | Weight | SVP Density | ICP Density | DCP Density | Total Retina | IRL | RPE | RNFL | IPL | INL | ORL | PRC | OPL | Number of Target Zone Crossings | % of time in sector with platform | Latency on Day 3 | Latency on Day 4 | Distance swam (m) | Velocity (m/s) | Thigmotaxis % | Floating % |
|---|---|---|---|---|---|---|---|---|---|---|---|---|---|---|---|---|---|---|---|---|---|
| Weight | | 0.1717 | 0.1801 | 0.1343 | 0.0824 | 0.0272 | 0.5563 | 0.6503 | 0.0854 | 0.0793 | 0.3884 | 0.4834 | 0.3099 | 0.1802 | 0.3306 | 0.3517 | 0.2242 | 0.2842 | 0.5073 | 0.6196 | 0.6156 |
| SVP Density | 0.1717 | | 0.0537 | 0.6069 | 0.2363 | 0.1541 | 0.0455 | 0.5787 | 0.6779 | 0.4187 | 0.5244 | 0.8625 | 0.0056 | 0.5895 | 0.4440 | 0.6610 | 0.9871 | 0.8234 | 0.5824 | 0.9669 | 0.7857 |
| ICP Density | 0.1801 | 0.0537 | | 0.7118 | 0.3685 | 0.2369 | 0.9300 | 0.2837 | 0.7189 | 0.2838 | 0.7451 | 0.8659 | 0.3218 | 0.0953 | 0.1504 | 0.4799 | 0.3091 | 0.1262 | 0.1031 | 0.0472 | 0.9405 |
| DCP Density | 0.1343 | 0.6069 | 0.7118 | | 0.8756 | 0.7433 | 0.1129 | 0.0613 | 0.0858 | 0.4840 | 0.8876 | 0.9045 | 0.2579 | 0.5523 | 0.8396 | 0.1847 | 0.1968 | 0.4888 | 0.8996 | 0.5692 | 0.4452 |
| Total Retina | 0.0824 | 0.2363 | 0.3685 | 0.8756 | | 0.0001 | 0.2021 | 0.8283 | 0.2066 | 0.0026 | 0.0025 | 0.0096 | 0.0717 | 0.0297 | 0.0566 | 0.9450 | 0.5571 | 0.2931 | 0.2315 | 0.3811 | 0.2065 |
| IRL | 0.0272 | 0.1541 | 0.2369 | 0.7433 | 0.0001 | | 0.1764 | 0.8753 | 0.1604 | 0.0010 | 0.0349 | 0.0728 | 0.0763 | 0.0249 | 0.0357 | 0.9807 | 0.6794 | 0.4219 | 0.3283 | 0.4931 | 0.4824 |
| RPE | 0.5563 | 0.0455 | 0.9300 | 0.1129 | 0.2021 | 0.1764 | | 0.3335 | 0.9330 | 0.5569 | 0.3518 | 0.6470 | 0.0021 | 0.9575 | 0.7687 | 0.6451 | 0.4662 | 0.8136 | 0.6263 | 0.4834 | 0.7133 |
| RNFL | 0.6503 | 0.5787 | 0.2837 | 0.0613 | 0.8283 | 0.8753 | 0.3335 | | 0.0185 | 0.8121 | 0.7812 | 0.9567 | 0.3010 | 0.6123 | 0.6502 | 0.1599 | 0.8188 | 0.4838 | 0.2918 | 0.4874 | 0.5549 |
| IPL | 0.0854 | 0.6779 | 0.7189 | 0.0858 | 0.2066 | 0.1604 | 0.9330 | 0.0185 | | 0.1057 | 0.4052 | 0.3742 | 0.8470 | 0.5607 | 0.5700 | 0.2178 | 0.9408 | 0.9071 | 0.7598 | 0.7799 | 0.3616 |
| INL | 0.0793 | 0.4187 | 0.2838 | 0.4840 | 0.0026 | 0.0010 | 0.5569 | 0.8121 | 0.1057 | | 0.0531 | 0.0618 | 0.3239 | 0.0054 | 0.0060 | 0.8511 | 0.7833 | 0.4862 | 0.3822 | 0.2972 | 0.3953 |
| ORL | 0.3884 | 0.5244 | 0.7451 | 0.8876 | 0.0025 | 0.0349 | 0.3518 | 0.7812 | 0.4052 | 0.0531 | | 0.0000 | 0.1409 | 0.1090 | 0.2030 | 0.8357 | 0.4393 | 0.2005 | 0.1804 | 0.2997 | 0.0312 |
| PRC | 0.4834 | 0.8625 | 0.8659 | 0.9045 | 0.0096 | 0.0728 | 0.6470 | 0.9567 | 0.3742 | 0.0618 | 0.0000 | | 0.3539 | 0.1022 | 0.2191 | 0.7958 | 0.3789 | 0.1965 | 0.2386 | 0.2536 | 0.0126 |
| OPL | 0.3099 | 0.0056 | 0.3218 | 0.2579 | 0.0717 | 0.0763 | 0.0021 | 0.3010 | 0.8470 | 0.3239 | 0.1409 | 0.3539 | | 0.5397 | 0.4711 | 0.9237 | 0.9918 | 0.5874 | 0.2476 | 0.8933 | 0.8108 |
| Number of Target Zone Crossings | 0.1802 | 0.5895 | 0.0953 | 0.5523 | 0.0297 | 0.0249 | 0.9575 | 0.6123 | 0.5607 | 0.0054 | 0.1090 | 0.1022 | 0.5397 | | 0.0006 | 0.7200 | 0.4078 | 0.2347 | 0.2967 | 0.0734 | 0.4857 |
| % of time in sector with platform | 0.3306 | 0.4440 | 0.1504 | 0.8396 | 0.0566 | 0.0357 | 0.7687 | 0.6502 | 0.5700 | 0.0060 | 0.2030 | 0.2191 | 0.4711 | 0.0006 | | 0.5064 | 0.9725 | 0.6497 | 0.5164 | 0.2017 | 0.8519 |
| Latency on Day 3 | 0.3517 | 0.6610 | 0.4799 | 0.1847 | 0.9450 | 0.9807 | 0.6451 | 0.1599 | 0.2178 | 0.8511 | 0.8357 | 0.7958 | 0.9237 | 0.7200 | 0.5064 | | 0.2562 | 0.7865 | 0.4797 | 0.6420 | 0.3187 |
| Latency on Day 4 | 0.2242 | 0.9871 | 0.3091 | 0.1968 | 0.5571 | 0.6794 | 0.4662 | 0.8188 | 0.9408 | 0.7833 | 0.4393 | 0.3789 | 0.9918 | 0.4078 | 0.9725 | 0.2562 | | 0.0033 | 0.2076 | 0.1100 | 0.1204 |
| Distance swam (m) | 0.2842 | 0.8234 | 0.1262 | 0.4888 | 0.2931 | 0.4219 | 0.8136 | 0.4838 | 0.9071 | 0.4862 | 0.2005 | 0.1965 | 0.5874 | 0.2347 | 0.6497 | 0.7865 | 0.0033 | | 0.0091 | 0.0108 | 0.0816 |
| Velocity (m/s) | 0.5073 | 0.5824 | 0.1031 | 0.8996 | 0.2315 | 0.3283 | 0.6263 | 0.2918 | 0.7598 | 0.3822 | 0.1804 | 0.2386 | 0.2476 | 0.2967 | 0.5164 | 0.4797 | 0.2076 | 0.0091 | | 0.0156 | 0.2187 |
| Thigmotaxis % | 0.6196 | 0.9669 | 0.0472 | 0.5692 | 0.3811 | 0.4931 | 0.4834 | 0.4874 | 0.7799 | 0.2972 | 0.2997 | 0.2536 | 0.8933 | 0.0734 | 0.2017 | 0.6420 | 0.1100 | 0.0108 | 0.0156 | | 0.1974 |
| Floating % | 0.6156 | 0.7857 | 0.9405 | 0.4452 | 0.2065 | 0.4824 | 0.7133 | 0.5549 | 0.3616 | 0.3953 | 0.0312 | 0.0126 | 0.8108 | 0.4857 | 0.8519 | 0.3187 | 0.1204 | 0.0816 | 0.2187 | 0.1974 | |

Table 3: Correlation Coefficient and corresponding p-value for or all tested parameters for transgenic female animals. For the Correlation coefficient red indicates values from 0-0.3, yellow =0.3-0.5 and green 0.5-1. For the p-value red indicates values above 0.15, yellow values from 0.15-0.1 and green values from below 0.05.

| R-Squared | Weight | SVP Density | ICP Density | DCP Density | Total Retina | IRL | RPE | RNFL | IPL | INL | ORL | PRC | OPL | Number of Target Zone Crossings | % of time in sector with platform | Latency on Day 3 | Latency on Day 4 | Distance swam (m) | Velocity (m/s) | Thigmotaxis % | Floating % |
|---|---|---|---|---|---|---|---|---|---|---|---|---|---|---|---|---|---|---|---|---|---|
| Weight | | 0.2864 | 0.3266 | 0.3329 | 0.4197 | 0.5840 | 0.0607 | 0.0365 | 0.4137 | 0.4262 | 0.1259 | 0.0851 | 0.1701 | 0.2771 | 0.1574 | 0.1452 | 0.2343 | 0.1873 | 0.0765 | 0.0436 | 0.0446 |
| SVP Density | 0.2864 | | 0.5578 | 0.0468 | 0.2240 | 0.3070 | 0.5136 | 0.0543 | 0.0307 | 0.1116 | 0.0707 | 0.0054 | 0.7476 | 0.0513 | 0.1006 | 0.0342 | 0.0000 | 0.0090 | 0.0533 | 0.0003 | 0.0133 |
| ICP Density | 0.3266 | 0.5578 | | 0.0297 | 0.1634 | 0.2652 | 0.0017 | 0.2237 | 0.0282 | 0.2236 | 0.0231 | 0.0063 | 0.1946 | 0.4573 | 0.3655 | 0.1043 | 0.2039 | 0.4021 | 0.4423 | 0.5781 | 0.0012 |
| DCP Density | 0.3329 | 0.0468 | 0.0297 | | 0.0044 | 0.0192 | 0.3647 | 0.4681 | 0.4128 | 0.0848 | 0.0036 | 0.0026 | 0.2066 | 0.0619 | 0.0074 | 0.2723 | 0.2600 | 0.0830 | 0.0029 | 0.0570 | 0.1001 |
| Total Retina | 0.4197 | 0.2240 | 0.1634 | 0.0044 | | 0.9246 | 0.2547 | 0.0085 | 0.2504 | 0.8016 | 0.8059 | 0.6997 | 0.4429 | 0.5728 | 0.4804 | 0.0009 | 0.0605 | 0.1811 | 0.2280 | 0.1296 | 0.2505 |
| IRL | 0.5840 | 0.3070 | 0.2652 | 0.0192 | 0.9246 | | 0.2812 | 0.0045 | 0.2994 | 0.8564 | 0.5510 | 0.4404 | 0.4324 | 0.5954 | 0.5480 | 0.0001 | 0.0304 | 0.1102 | 0.1587 | 0.0815 | 0.0854 |
| RPE | 0.0607 | 0.5136 | 0.0017 | 0.3647 | 0.2547 | 0.2812 | | 0.1556 | 0.0013 | 0.0605 | 0.1452 | 0.0373 | 0.8156 | 0.0005 | 0.0155 | 0.0377 | 0.0916 | 0.0100 | 0.0420 | 0.0850 | 0.0241 |
| RNFL | 0.0365 | 0.0543 | 0.2237 | 0.4681 | 0.0085 | 0.0045 | 0.1556 | | 0.6312 | 0.0102 | 0.0139 | 0.0005 | 0.1759 | 0.0454 | 0.0365 | 0.3000 | 0.0095 | 0.0849 | 0.1820 | 0.0836 | 0.0612 |
| IPL | 0.4137 | 0.0307 | 0.0282 | 0.4128 | 0.2504 | 0.2994 | 0.0013 | 0.6312 | | 0.3764 | 0.1178 | 0.1331 | 0.0067 | 0.0594 | 0.0567 | 0.2400 | 0.0010 | 0.0025 | 0.0168 | 0.0140 | 0.1397 |
| INL | 0.4262 | 0.1116 | 0.2236 | 0.0848 | 0.8016 | 0.8564 | 0.0605 | 0.0102 | 0.3764 | | 0.4905 | 0.4668 | 0.1613 | 0.7499 | 0.7414 | 0.0064 | 0.0136 | 0.0840 | 0.1290 | 0.1784 | 0.1225 |
| ORL | 0.1259 | 0.0707 | 0.0231 | 0.0036 | 0.8059 | 0.5510 | 0.1452 | 0.0139 | 0.1178 | 0.4905 | | 0.9540 | 0.3239 | 0.3710 | 0.2538 | 0.0078 | 0.1026 | 0.2562 | 0.2768 | 0.1767 | 0.5664 |
| PRC | 0.0851 | 0.0054 | 0.0063 | 0.0026 | 0.6997 | 0.4404 | 0.0373 | 0.0005 | 0.1331 | 0.4668 | 0.9540 | | 0.1440 | 0.3824 | 0.2389 | 0.0120 | 0.1307 | 0.2602 | 0.2220 | 0.2099 | 0.6729 |
| OPL | 0.1701 | 0.7476 | 0.1946 | 0.2066 | 0.4429 | 0.4324 | 0.8156 | 0.1759 | 0.0067 | 0.1613 | 0.3239 | 0.1440 | | 0.0658 | 0.0897 | 0.0017 | 0.0000 | 0.0519 | 0.2146 | 0.0032 | 0.0103 |
| Number of Target Zone Crossings | 0.2771 | 0.0513 | 0.4573 | 0.0619 | 0.5728 | 0.5954 | 0.0005 | 0.0454 | 0.0594 | 0.7499 | 0.3710 | 0.3824 | 0.0658 | | 0.8782 | 0.0230 | 0.1166 | 0.2252 | 0.1787 | 0.4391 | 0.0842 |
| % of time in sector with platform | 0.1574 | 0.1006 | 0.3655 | 0.0074 | 0.4804 | 0.5480 | 0.0155 | 0.0365 | 0.0567 | 0.7414 | 0.2538 | 0.2389 | 0.0897 | 0.8782 | | 0.0768 | 0.0002 | 0.0366 | 0.0734 | 0.2551 | 0.0063 |
| Latency on Day 3 | 0.1452 | 0.0342 | 0.1043 | 0.2723 | 0.0009 | 0.0001 | 0.0377 | 0.3000 | 0.2400 | 0.0064 | 0.0078 | 0.0120 | 0.0017 | 0.0230 | 0.0768 | | 0.2079 | 0.0132 | 0.0864 | 0.0384 | 0.1646 |
| Latency on Day 4 | 0.2343 | 0.0000 | 0.2039 | 0.2600 | 0.0605 | 0.0304 | 0.0916 | 0.0095 | 0.0010 | 0.0136 | 0.1026 | 0.1307 | 0.0000 | 0.1166 | 0.0002 | 0.2079 | | 0.7860 | 0.2494 | 0.3694 | 0.3529 |
| Distance swam (m) | 0.1873 | 0.0090 | 0.4021 | 0.0830 | 0.1811 | 0.1102 | 0.0100 | 0.0849 | 0.0025 | 0.0840 | 0.2562 | 0.2602 | 0.0519 | 0.2252 | 0.0366 | 0.0132 | 0.7860 | | 0.7055 | 0.6885 | 0.4213 |
| Velocity (m/s) | 0.0765 | 0.0533 | 0.4423 | 0.0029 | 0.2280 | 0.1587 | 0.0420 | 0.1820 | 0.0168 | 0.1290 | 0.2768 | 0.2220 | 0.2146 | 0.1787 | 0.0734 | 0.0864 | 0.2494 | 0.7055 | | 0.6504 | 0.2392 |
| Thigmotaxis % | 0.0436 | 0.0003 | 0.5781 | 0.0570 | 0.1296 | 0.0815 | 0.0850 | 0.0836 | 0.0140 | 0.1784 | 0.1767 | 0.2099 | 0.0032 | 0.4391 | 0.2551 | 0.0384 | 0.3694 | 0.6885 | 0.6504 | | 0.2593 |
| Floating % | 0.0446 | 0.0133 | 0.0012 | 0.1001 | 0.2505 | 0.0854 | 0.0241 | 0.0612 | 0.1397 | 0.1225 | 0.5664 | 0.6729 | 0.0103 | 0.0842 | 0.0063 | 0.1646 | 0.3529 | 0.4213 | 0.2392 | 0.2593 | |

| Mean Squared Error | Weight | SVP Density | ICP Density | DCP Density | Total Retina | IRL | RPE | RNFL | IPL | INL | ORL | PRC | OPL | Number of Target Zone Crossings | % of time in sector with platform | Latency on Day 3 | Latency on Day 4 | Distance swam (m) | Velocity (m/s) | Thigmotaxis % | Floating % |
|---|---|---|---|---|---|---|---|---|---|---|---|---|---|---|---|---|---|---|---|---|---|
| Weight | | 2.5E-05 | 0.00015 | 0.00015 | 20.44 | 6.33229 | 0.10762 | 16.2326 | 12.91629 | 0.752343 | 5.16847 | 4.27288 | 0.2349 | 4.834440228 | 663.696694 | 138.840028 | 79.4472081 | 55822.08828 | 6.874115 | 517.4121637 | 30.022995 |
| SVP Density | 2.52E-05 | | 9.9E-05 | 0.00022 | 27.33383 | 10.5499 | 0.05572 | 15.9331 | 21.35128 | 1.164792 | 5.49483 | 4.64481 | 0.07143 | 6.344224871 | 708.4117567 | 156.863957 | 103.753507 | 68069.60648 | 7.047 | 540.8494704 | 31.007236 |
| ICP Density | 0.000151 | 9.9E-05 | | 0.00025 | 24.97269 | 9.49204 | 0.08741 | 14.4528 | 18.46218 | 1.020539 | 5.40026 | 4.85852 | 0.12199 | 4.097966629 | 550.4214534 | 120.405659 | 94.3608178 | 46933.0212 | 4.738972 | 254.1960718 | 34.221001 |
| DCP Density | 0.000151 | 0.00022 | 0.00025 | | 35.06596 | 14.931 | 0.07279 | 8.96152 | 12.93424 | 1.199863 | 5.89168 | 4.65796 | 0.22459 | 6.273354004 | 781.8119765 | 118.198407 | 76.7863436 | 62982.25457 | 7.422007 | 510.2066824 | 28.279017 |
| Total Retina | 20.44 | 27.3338 | 24.9727 | 35.066 | | 1.14758 | 0.08539 | 16.7048 | 16.51306 | 0.260099 | 1.14758 | 1.40245 | 0.15769 | 2.856941366 | 409.2282346 | 162.281898 | 97.4824336 | 56243.87753 | 5.746362 | 470.9243754 | 23.55293 |
| IRL | 6.332288 | 10.5499 | 9.49204 | 14.931 | 1.14758 | | 0.08236 | 16.7727 | 15.43253 | 0.188337 | 2.65507 | 2.61339 | 0.16065 | 2.705921384 | 355.9805403 | 162.404517 | 100.599445 | 61119.45683 | 6.262087 | 496.9173903 | 28.740839 |
| RPE | 0.107617 | 0.05572 | 0.08741 | 0.07279 | 0.085389 | 0.08236 | | 14.2259 | 22.0004 | 1.231716 | 5.05472 | 4.49609 | 0.05221 | 6.684059693 | 775.4015023 | 156.30116 | 94.2532587 | 67998.18084 | 7.130734 | 495.0114719 | 30.666276 |
| RNFL | 16.23255 | 15.9331 | 14.4528 | 8.96152 | 16.70475 | 16.7727 | 14.2259 | | 8.124881 | 1.297763 | 5.83102 | 4.66762 | 0.23328 | 6.383710706 | 758.8527523 | 113.697651 | 102.777549 | 62854.42543 | 6.08861 | 495.8126219 | 29.503177 |
| IPL | 12.91629 | 21.3513 | 18.4622 | 12.9342 | 16.51306 | 15.4325 | 22.0004 | 8.12488 | | 0.817556 | 5.21646 | 4.04854 | 0.28116 | 6.290025214 | 742.9512242 | 123.440902 | 103.654941 | 68517.23901 | 7.318519 | 533.4222064 | 27.033446 |
| INL | 0.752343 | 1.16479 | 1.02054 | 1.19986 | 0.260099 | 0.18834 | 1.23172 | 1.29776 | 0.817556 | | 3.01296 | 2.49026 | 0.23739 | 1.672713718 | 203.717232 | 161.389098 | 102.347512 | 62915.44673 | 6.483088 | 444.5053991 | 27.574258 |
| ORL | 5.168471 | 5.49483 | 5.40026 | 5.89168 | 1.14758 | 2.65507 | 5.05472 | 5.83102 | 5.216462 | 3.012961 | | 0.21503 | 0.19137 | 4.20676334 | 587.6953119 | 161.161121 | 93.1152807 | 51086.40764 | 5.382901 | 445.4122402 | 13.625804 |
| PRC | 4.272876 | 4.64481 | 4.85852 | 4.65796 | 1.402449 | 2.61339 | 4.49609 | 4.66762 | 4.048542 | 2.490264 | 0.21503 | | 0.24231 | 4.130184751 | 599.4770359 | 160.464892 | 90.1973081 | 50811.68071 | 5.791254 | 427.4618796 | 10.277708 |
| OPL | 0.234902 | 0.07143 | 0.12199 | 0.22459 | 0.157687 | 0.16065 | 0.05221 | 0.23328 | 0.281159 | 0.237389 | 0.19137 | 0.24231 | | 6.247431646 | 716.972355 | 162.152076 | 103.756411 | 65121.00457 | 5.845865 | 539.259629 | 31.100617 |
| Number of Target Zone Crossings | 4.83444 | 6.34422 | 4.09797 | 6.27335 | 2.856941 | 2.70592 | 6.68406 | 6.38371 | 6.290025 | 1.672714 | 4.20676 | 4.13018 | 6.24743 | | 95.97227527 | 158.687267 | 91.662074 | 53216.20793 | 6.113215 | 303.4833116 | 28.778599 |
| % of time in sector with platform | 663.6967 | 708.412 | 550.421 | 781.812 | 409.2282 | 355.981 | 775.402 | 758.853 | 742.9512 | 203.7172 | 587.695 | 599.477 | 716.972 | 95.97227527 | | 149.95037 | 103.736116 | 66169.38227 | 6.897307 | 403.0187987 | 31.227192 |
| Latency on Day 3 | 138.84 | 156.864 | 120.406 | 118.198 | 162.2819 | 162.405 | 156.301 | 113.698 | 123.4409 | 161.3891 | 161.161 | 160.465 | 162.152 | 158.6872667 | 149.9503703 | | 82.1862292 | 67779.95328 | 6.800146 | 520.2629277 | 26.253141 |
| Latency on Day 4 | 79.44721 | 103.754 | 94.3608 | 76.7863 | 97.48243 | 100.599 | 94.2533 | 102.778 | 103.6549 | 102.3475 | 93.1153 | 90.1973 | 103.756 | 91.66207398 | 103.7361156 | 82.1862292 | | 14700.96181 | 5.586709 | 341.1856838 | 20.334319 |
| Distance swam (m) | 55822.09 | 68069.6 | 46933 | 62982.3 | 56243.88 | 61119.5 | 67998.2 | 62854.4 | 68517.24 | 62915.45 | 51086.4 | 50811.7 | 65121 | 53216.20793 | 66169.38227 | 67779.9533 | 14700.9618 | | 2.192179 | 168.5024331 | 18.184275 |
| Velocity (m/s) | 6.874115 | 7.047 | 4.73897 | 7.42201 | 5.746362 | 6.26209 | 7.13073 | 6.08861 | 7.318519 | 6.483088 | 5.3829 | 5.79125 | 5.84586 | 6.11321548 | 6.897306998 | 6.80014582 | 5.58670878 | 2.192179392 | | 189.1522381 | 23.906569 |
| Thigmotaxis % | 517.4122 | 540.849 | 254.196 | 510.207 | 470.9244 | 496.917 | 495.011 | 495.813 | 533.4222 | 444.5054 | 445.412 | 427.462 | 539.26 | 303.4833116 | 403.0187987 | 520.262928 | 341.185684 | 168.5024331 | 189.1522 | | 23.276919 |
| Floating % | 30.023 | 31.0072 | 34.221 | 28.279 | 23.55293 | 28.7408 | 30.6663 | 29.5032 | 27.03345 | 27.57426 | 13.6258 | 10.2777 | 31.1006 | 28.77859921 | 31.22719228 | 26.2531411 | 20.3343194 | 18.1842748 | 23.90657 | 23.27691934 | |

Table 4: R-squared and corresponding MSE for or all tested parameters for transgenic female animals. For the R-squared values red indicates values from 0-0.3, yellow =0.4-0.6 and green 0.6-1.

| Correlation Coefficient | Weight | SVP Density | ICP Density | DCP Density | Total Retina | IRL | RPE | RNFL | IPL | INL | ORL | PRC | OPL | Number of Target Zone Crossings | % of time in sector with platform | Latency on Day 3 | Latency on Day 4 | Distance swam (m) | Velocity (m/s) | Thigmotaxis % | Floating % |
|---|---|---|---|---|---|---|---|---|---|---|---|---|---|---|---|---|---|---|---|---|---|
| Weight | | 0.9647 | -0.4067 | 0.5965 | 0.0326 | 0.4723 | -0.3850 | -0.3904 | 0.6394 | 0.6159 | -0.2827 | -0.2279 | -0.3112 | 0.5181 | 0.5155 | 0.7656 | -0.1823 | -0.2081 | -0.1187 | -0.0557 | 0.0752 |
| SVP Density | 0.9647 | | -0.6232 | 0.6485 | -0.4845 | 0.2209 | -0.4744 | -0.6945 | 0.8426 | 0.7571 | -0.7987 | -0.8585 | -0.4867 | 0.2974 | 0.5438 | 0.8488 | 0.0277 | -0.0233 | 0.0591 | 0.0822 | -0.1988 |
| ICP Density | -0.4067 | -0.6232 | | -0.1271 | 0.5943 | 0.1868 | 0.7312 | 0.7280 | -0.5855 | -0.8241 | 0.6971 | 0.5460 | 0.8340 | 0.3921 | -0.4815 | -0.6633 | -0.4138 | -0.3699 | -0.2085 | -0.3128 | 0.5402 |
| DCP Density | 0.5965 | 0.6485 | -0.1271 | | -0.0410 | 0.2447 | 0.3480 | -0.0164 | 0.1853 | 0.1276 | -0.2373 | -0.3515 | 0.2403 | 0.4879 | 0.7163 | 0.7737 | 0.0411 | -0.0864 | 0.0314 | -0.0572 | -0.2498 |
| Total Retina | 0.0326 | -0.4845 | 0.5943 | -0.0410 | | 0.8849 | 0.3722 | 0.8289 | -0.6481 | 0.5573 | 0.9443 | 0.9248 | 0.5277 | 0.4142 | 0.3342 | 0.0103 | -0.5052 | -0.5152 | -0.6389 | -0.5411 | 0.5608 |
| IRL | 0.4723 | 0.2209 | 0.1868 | 0.2447 | 0.8849 | | 0.1553 | 0.5426 | -0.2570 | 0.7656 | 0.6823 | 0.6861 | 0.3251 | 0.4957 | 0.5448 | 0.3818 | -0.5866 | -0.6184 | -0.7014 | -0.5919 | 0.5590 |
| RPE | -0.3850 | -0.4744 | 0.7312 | 0.3480 | 0.3722 | 0.1553 | | 0.7836 | -0.7025 | -0.4165 | 0.4746 | 0.2444 | 0.9548 | 0.2263 | 0.1053 | -0.1922 | -0.2113 | -0.2872 | -0.2084 | -0.3589 | 0.1652 |
| RNFL | -0.3904 | -0.6945 | 0.7280 | -0.0164 | 0.8289 | 0.5426 | 0.7836 | | -0.9139 | 0.0476 | 0.9177 | 0.7927 | 0.8445 | 0.2858 | 0.1550 | -0.2609 | -0.3512 | -0.3836 | -0.4537 | -0.4669 | 0.4018 |
| IPL | 0.6394 | 0.8426 | -0.5855 | 0.1853 | -0.6481 | -0.2570 | -0.7025 | -0.9139 | | 0.0704 | -0.8357 | -0.7510 | -0.6783 | -0.1509 | -0.0855 | 0.3657 | -0.0049 | 0.0187 | 0.1465 | 0.1501 | -0.0860 |
| INL | 0.6159 | 0.7571 | -0.8241 | 0.1276 | 0.5573 | 0.7656 | -0.4165 | 0.0476 | 0.0704 | | 0.3338 | 0.4903 | -0.3249 | 0.3791 | 0.6529 | 0.6175 | -0.1198 | -0.1344 | -0.3131 | -0.1136 | 0.1069 |
| ORL | -0.2827 | -0.7987 | 0.6971 | -0.2373 | 0.9443 | 0.6823 | 0.4746 | 0.9177 | -0.8357 | 0.3338 | | 0.9667 | 0.5985 | 0.2998 | 0.1395 | -0.2537 | -0.3784 | -0.3716 | -0.5072 | -0.4311 | 0.4852 |
| PRC | -0.2279 | -0.8585 | 0.5460 | -0.3515 | 0.9248 | 0.6861 | 0.2444 | 0.7927 | -0.7510 | 0.4903 | 0.9667 | | 0.3735 | 0.2632 | 0.1606 | -0.2038 | -0.2913 | -0.2680 | -0.4552 | -0.3262 | 0.4205 |
| OPL | -0.3112 | -0.4867 | 0.8340 | 0.2403 | 0.5277 | 0.3251 | 0.9548 | 0.8445 | -0.6783 | -0.3249 | 0.5985 | 0.3735 | | 0.2628 | 0.0029 | -0.2815 | -0.4596 | -0.5079 | -0.4133 | -0.5414 | 0.4424 |
| Number of Target Zone Crossings | 0.5181 | 0.2974 | 0.3921 | 0.4879 | 0.4142 | 0.4957 | 0.2263 | 0.2858 | -0.1509 | 0.3791 | 0.2998 | 0.2632 | 0.2628 | | 0.4235 | 0.3377 | 0.0582 | 0.0533 | 0.1648 | 0.1935 | -0.0067 |
| % of time in sector with platform | 0.5155 | 0.5438 | -0.4815 | 0.7163 | 0.3342 | 0.5448 | 0.1053 | 0.1550 | -0.0855 | 0.6529 | 0.1395 | 0.1606 | 0.0029 | 0.4235 | | 0.8954 | 0.1668 | 0.0403 | -0.0850 | -0.0222 | -0.3268 |
| Latency on Day 3 | 0.7656 | 0.8488 | -0.6633 | 0.7737 | 0.0103 | 0.3818 | -0.1922 | -0.2609 | 0.3657 | 0.6175 | -0.2537 | -0.2038 | -0.2815 | 0.3377 | 0.8954 | | 0.1582 | 0.0484 | -0.0027 | 0.0500 | -0.3515 |
| Latency on Day 4 | -0.1823 | 0.0277 | -0.4138 | 0.0411 | -0.5052 | -0.5866 | -0.2113 | -0.3512 | -0.0049 | -0.1198 | -0.3784 | -0.2913 | -0.4596 | 0.0582 | 0.1668 | 0.1582 | | 0.9865 | 0.9190 | 0.9336 | -0.9608 |
| Distance swam (m) | -0.2081 | -0.0233 | -0.3699 | -0.0864 | -0.5152 | -0.6184 | -0.2872 | -0.3836 | 0.0187 | -0.1344 | -0.3716 | -0.2680 | -0.5079 | 0.0533 | 0.0403 | 0.0484 | 0.9865 | | 0.9420 | 0.9687 | -0.9080 |
| Velocity (m/s) | -0.1187 | 0.0591 | -0.2085 | 0.0314 | -0.6389 | -0.7014 | -0.2084 | -0.4537 | 0.1465 | -0.3131 | -0.5072 | -0.4552 | -0.4133 | 0.1648 | -0.0850 | -0.0027 | 0.9190 | 0.9420 | | 0.9724 | -0.8444 |
| Thigmotaxis % | -0.0557 | 0.0822 | -0.3128 | -0.0572 | -0.5411 | -0.5919 | -0.3589 | -0.4669 | 0.1501 | -0.1136 | -0.4311 | -0.3262 | -0.5414 | 0.1935 | -0.0222 | 0.0500 | 0.9336 | 0.9687 | 0.9724 | | -0.8351 |
| Floating % | 0.0752 | -0.1988 | 0.5402 | -0.2498 | 0.5608 | 0.5590 | 0.1652 | 0.4018 | -0.0860 | 0.1069 | 0.4852 | 0.4205 | 0.4424 | -0.0067 | -0.3268 | -0.3515 | -0.9608 | -0.9080 | -0.8444 | -0.8351 | |

| P-Value | Weight | SVP Density | ICP Density | DCP Density | Total Retina | IRL | RPE | RNFL | IPL | INL | ORL | PRC | OPL | Number of Target Zone Crossings | % of time in sector with platform | Latency on Day 3 | Latency on Day 4 | Distance swam (m) | Velocity (m/s) | Thigmotaxis % | Floating % |
|---|---|---|---|---|---|---|---|---|---|---|---|---|---|---|---|---|---|---|---|---|---|
| Weight | | 0.0079 | 0.4969 | 0.2114 | 0.9512 | 0.3442 | 0.4511 | 0.4441 | 0.1716 | 0.1930 | 0.5873 | 0.6641 | 0.5483 | 0.2924 | 0.2952 | 0.0759 | 0.7296 | 0.6923 | 0.8228 | 0.9165 | 0.8874 |
| SVP Density | 0.0079 | | 0.2614 | 0.2365 | 0.4082 | 0.7211 | 0.4195 | 0.1931 | 0.0732 | 0.1383 | 0.1051 | 0.0625 | 0.4057 | 0.6270 | 0.3434 | 0.0689 | 0.9647 | 0.9703 | 0.9248 | 0.8954 | 0.7486 |
| ICP Density | 0.4969 | 0.2614 | | 0.8386 | 0.2906 | 0.7636 | 0.1604 | 0.1631 | 0.2996 | 0.0862 | 0.1907 | 0.3411 | 0.0792 | 0.5139 | 0.4115 | 0.2223 | 0.4886 | 0.5400 | 0.7365 | 0.6084 | 0.3473 |
| DCP Density | 0.2114 | 0.2365 | 0.8386 | | 0.9385 | 0.6403 | 0.4991 | 0.9754 | 0.7252 | 0.8097 | 0.6508 | 0.4944 | 0.6465 | 0.3263 | 0.1093 | 0.0710 | 0.9384 | 0.8707 | 0.9529 | 0.9143 | 0.6331 |
| Total Retina | 0.9512 | 0.4082 | 0.2906 | 0.9385 | | 0.0191 | 0.4674 | 0.0414 | 0.1639 | 0.2506 | 0.0046 | 0.0083 | 0.2820 | 0.4142 | 0.5174 | 0.9846 | 0.3067 | 0.2956 | 0.1720 | 0.2675 | 0.2470 |
| IRL | 0.3442 | 0.7211 | 0.7636 | 0.6403 | 0.0191 | | 0.7690 | 0.2659 | 0.6230 | 0.0760 | 0.1354 | 0.1323 | 0.5296 | 0.3173 | 0.2636 | 0.4552 | 0.2210 | 0.1906 | 0.1205 | 0.2158 | 0.2488 |
| RPE | 0.4511 | 0.4195 | 0.1604 | 0.4991 | 0.4674 | 0.7690 | | 0.0652 | 0.1196 | 0.4114 | 0.3416 | 0.6407 | 0.0030 | 0.6664 | 0.8427 | 0.7153 | 0.6878 | 0.5811 | 0.6919 | 0.4848 | 0.7545 |
| RNFL | 0.4441 | 0.1931 | 0.1631 | 0.9754 | 0.0414 | 0.2659 | 0.0652 | | 0.0108 | 0.9286 | 0.0099 | 0.0600 | 0.0344 | 0.5830 | 0.7694 | 0.6175 | 0.4948 | 0.4528 | 0.3661 | 0.3505 | 0.4298 |
| IPL | 0.1716 | 0.0732 | 0.2996 | 0.7252 | 0.1639 | 0.6230 | 0.1196 | 0.0108 | | 0.8946 | 0.0383 | 0.0853 | 0.1386 | 0.7753 | 0.8721 | 0.4759 | 0.9927 | 0.9719 | 0.7819 | 0.7765 | 0.8713 |
| INL | 0.1930 | 0.1383 | 0.0862 | 0.8097 | 0.2506 | 0.0760 | 0.4114 | 0.9286 | 0.8946 | | 0.5179 | 0.3234 | 0.5298 | 0.4586 | 0.1598 | 0.1915 | 0.8211 | 0.7996 | 0.5457 | 0.8303 | 0.8403 |
| ORL | 0.5873 | 0.1051 | 0.1907 | 0.6508 | 0.0046 | 0.1354 | 0.3416 | 0.0099 | 0.0383 | 0.5179 | | 0.0016 | 0.2094 | 0.5638 | 0.7921 | 0.6277 | 0.4595 | 0.4682 | 0.3045 | 0.3934 | 0.3293 |
| PRC | 0.6641 | 0.0625 | 0.3411 | 0.4944 | 0.0083 | 0.1323 | 0.6407 | 0.0600 | 0.0853 | 0.3234 | 0.0016 | | 0.4658 | 0.6143 | 0.7611 | 0.6986 | 0.5754 | 0.6076 | 0.3643 | 0.5281 | 0.4064 |
| OPL | 0.5483 | 0.4057 | 0.0792 | 0.6465 | 0.2820 | 0.5296 | 0.0030 | 0.0344 | 0.1386 | 0.5298 | 0.2094 | 0.4658 | | 0.6149 | 0.9957 | 0.5889 | 0.3591 | 0.3037 | 0.4153 | 0.2673 | 0.3796 |
| Number of Target Zone Crossings | 0.2924 | 0.6270 | 0.5139 | 0.3263 | 0.4142 | 0.3173 | 0.6664 | 0.5830 | 0.7753 | 0.4586 | 0.5638 | 0.6143 | 0.6149 | | 0.4027 | 0.5127 | 0.9128 | 0.9201 | 0.7550 | 0.7134 | 0.9899 |
| % of time in sector with platform | 0.2952 | 0.3434 | 0.4115 | 0.1093 | 0.5174 | 0.2636 | 0.8427 | 0.7694 | 0.8721 | 0.1598 | 0.7921 | 0.7611 | 0.9957 | 0.4027 | | 0.0158 | 0.7521 | 0.9396 | 0.8727 | 0.9666 | 0.5273 |
| Latency on Day 3 | 0.0759 | 0.0689 | 0.2223 | 0.0710 | 0.9846 | 0.4552 | 0.7153 | 0.6175 | 0.4759 | 0.1915 | 0.6277 | 0.6986 | 0.5889 | 0.5127 | 0.0158 | | 0.7646 | 0.9275 | 0.9960 | 0.9250 | 0.4944 |
| Latency on Day 4 | 0.7296 | 0.9647 | 0.4886 | 0.9384 | 0.3067 | 0.2210 | 0.6878 | 0.4948 | 0.9927 | 0.8211 | 0.4595 | 0.5754 | 0.3591 | 0.9128 | 0.7521 | 0.7646 | | 0.0003 | 0.0096 | 0.0065 | 0.0023 |
| Distance swam (m) | 0.6923 | 0.9703 | 0.5400 | 0.8707 | 0.2956 | 0.1906 | 0.5811 | 0.4528 | 0.9719 | 0.7996 | 0.4682 | 0.6076 | 0.3037 | 0.9201 | 0.9396 | 0.9275 | 0.0003 | | 0.0049 | 0.0014 | 0.0123 |
| Velocity (m/s) | 0.8228 | 0.9248 | 0.7365 | 0.9529 | 0.1720 | 0.1205 | 0.6919 | 0.3661 | 0.7819 | 0.5457 | 0.3045 | 0.3643 | 0.4153 | 0.7550 | 0.8727 | 0.9960 | 0.0096 | 0.0049 | | 0.0011 | 0.0344 |
| Thigmotaxis % | 0.9165 | 0.8954 | 0.6084 | 0.9143 | 0.2675 | 0.2158 | 0.4848 | 0.3505 | 0.7765 | 0.8303 | 0.3934 | 0.5281 | 0.2673 | 0.7134 | 0.9666 | 0.9250 | 0.0065 | 0.0014 | 0.0011 | | 0.0385 |
| Floating % | 0.8874 | 0.7486 | 0.3473 | 0.6331 | 0.2470 | 0.2488 | 0.7545 | 0.4298 | 0.8713 | 0.8403 | 0.3293 | 0.4064 | 0.3796 | 0.9899 | 0.5273 | 0.4944 | 0.0023 | 0.0123 | 0.0344 | 0.0385 | |

Table 5: Correlation Coefficient and corresponding p-value for or all tested parameters for transgenic male animals. For the Correlation coefficient red indicates values from 0-0.3, yellow =0.3-0.5 and green 0.5-1. For the p-value red indicates values above 0.15, yellow values from 0.15-0.1 and green values from below 0.05.

| R-Squared | Weight | SVP Density | ICP Density | DCP Density | Total Retina | IRL | RPE | RNFL | IPL | INL | ORL | PRC | OPL | Number of Target Zone Crossings | % of time in sector with platform | Latency on Day 3 | Latency on Day 4 | Distance swam (m) | Velocity (m/s) | Thigmotaxis % | Floating % |
|---|---|---|---|---|---|---|---|---|---|---|---|---|---|---|---|---|---|---|---|---|---|
| Weight | | 0.9306 | 0.1654 | 0.3558 | 0.0011 | 0.2231 | 0.1482 | 0.1524 | 0.4088 | 0.3793 | 0.0799 | 0.0519 | 0.0968 | 0.2684 | 0.2658 | 0.5862 | 0.0332 | 0.0433 | 0.0141 | 0.0031 | 0.0057 |
| SVP Density | 0.9306 | | 0.3884 | 0.4205 | 0.2347 | 0.0488 | 0.2251 | 0.4824 | 0.7099 | 0.5732 | 0.6380 | 0.7370 | 0.2369 | 0.0884 | 0.2958 | 0.7205 | 0.0008 | 0.0005 | 0.0035 | 0.0068 | 0.0395 |
| ICP Density | 0.1654 | 0.3884 | | 0.0162 | 0.3531 | 0.0349 | 0.5346 | 0.5300 | 0.3428 | 0.6792 | 0.4860 | 0.2981 | 0.6955 | 0.1537 | 0.2318 | 0.4400 | 0.1712 | 0.1368 | 0.0435 | 0.0978 | 0.2918 |
| DCP Density | 0.3558 | 0.4205 | 0.0162 | | 0.0017 | 0.0599 | 0.1211 | 0.0003 | 0.0344 | 0.0163 | 0.0563 | 0.1236 | 0.0578 | 0.2380 | 0.5131 | 0.5987 | 0.0017 | 0.0075 | 0.0010 | 0.0033 | 0.0624 |
| Total Retina | 0.0011 | 0.2347 | 0.3531 | 0.0017 | | 0.7831 | 0.1386 | 0.6871 | 0.4201 | 0.3106 | 0.8916 | 0.8552 | 0.2784 | 0.1716 | 0.1117 | 0.0001 | 0.2552 | 0.2654 | 0.4082 | 0.2928 | 0.3145 |
| IRL | 0.2231 | 0.0488 | 0.0349 | 0.0599 | 0.7831 | | 0.0241 | 0.2945 | 0.0660 | 0.5861 | 0.4655 | 0.4708 | 0.1057 | 0.2458 | 0.2968 | 0.1457 | 0.3441 | 0.3824 | 0.4919 | 0.3504 | 0.3125 |
| RPE | 0.1482 | 0.2251 | 0.5346 | 0.1211 | 0.1386 | 0.0241 | | 0.6140 | 0.4936 | 0.1735 | 0.2252 | 0.0597 | 0.9117 | 0.0512 | 0.0111 | 0.0369 | 0.0446 | 0.0825 | 0.0434 | 0.1288 | 0.0273 |
| RNFL | 0.1524 | 0.4824 | 0.5300 | 0.0003 | 0.6871 | 0.2945 | 0.6140 | | 0.8352 | 0.0023 | 0.8421 | 0.6284 | 0.7132 | 0.0817 | 0.0240 | 0.0681 | 0.1233 | 0.1471 | 0.2059 | 0.2180 | 0.1614 |
| IPL | 0.4088 | 0.7099 | 0.3428 | 0.0344 | 0.4201 | 0.0660 | 0.4936 | 0.8352 | | 0.0050 | 0.6984 | 0.5639 | 0.4601 | 0.0228 | 0.0073 | 0.1338 | 0.0000 | 0.0004 | 0.0215 | 0.0225 | 0.0074 |
| INL | 0.3793 | 0.5732 | 0.6792 | 0.0163 | 0.3106 | 0.5861 | 0.1735 | 0.0023 | 0.0050 | | 0.1114 | 0.2404 | 0.1056 | 0.1437 | 0.4263 | 0.3813 | 0.0144 | 0.0181 | 0.0980 | 0.0129 | 0.0114 |
| ORL | 0.0799 | 0.6380 | 0.4860 | 0.0563 | 0.8916 | 0.4655 | 0.2252 | 0.8421 | 0.6984 | 0.1114 | | 0.9345 | 0.3582 | 0.0899 | 0.0195 | 0.0643 | 0.1432 | 0.1381 | 0.2572 | 0.1858 | 0.2355 |
| PRC | 0.0519 | 0.7370 | 0.2981 | 0.1236 | 0.8552 | 0.4708 | 0.0597 | 0.6284 | 0.5639 | 0.2404 | 0.9345 | | 0.1395 | 0.0693 | 0.0258 | 0.0415 | 0.0849 | 0.0718 | 0.2072 | 0.1064 | 0.1768 |
| OPL | 0.0968 | 0.2369 | 0.6955 | 0.0578 | 0.2784 | 0.1057 | 0.9117 | 0.7132 | 0.4601 | 0.1056 | 0.3582 | 0.1395 | | 0.0690 | 0.0000 | 0.0792 | 0.2113 | 0.2580 | 0.1708 | 0.2931 | 0.1958 |
| Number of Target Zone Crossings | 0.2684 | 0.0884 | 0.1537 | 0.2380 | 0.1716 | 0.2458 | 0.0512 | 0.0817 | 0.0228 | 0.1437 | 0.0899 | 0.0693 | 0.0690 | | 0.1793 | 0.1140 | 0.0034 | 0.0028 | 0.0272 | 0.0374 | 0.0000 |
| % of time in sector with platform | 0.2658 | 0.2958 | 0.2318 | 0.5131 | 0.1117 | 0.2968 | 0.0111 | 0.0240 | 0.0073 | 0.4263 | 0.0195 | 0.0258 | 0.0000 | 0.1793 | | 0.8017 | 0.0278 | 0.0016 | 0.0072 | 0.0005 | 0.1068 |
| Latency on Day 3 | 0.5862 | 0.7205 | 0.4400 | 0.5987 | 0.0001 | 0.1457 | 0.0369 | 0.0681 | 0.1338 | 0.3813 | 0.0643 | 0.0415 | 0.0792 | 0.1140 | 0.8017 | | 0.0250 | 0.0023 | 0.0000 | 0.0025 | 0.1236 |
| Latency on Day 4 | 0.0332 | 0.0008 | 0.1712 | 0.0017 | 0.2552 | 0.3441 | 0.0446 | 0.1233 | 0.0000 | 0.0144 | 0.1432 | 0.0849 | 0.2113 | 0.0034 | 0.0278 | 0.0250 | | 0.9732 | 0.8446 | 0.8716 | 0.9231 |
| Distance swam (m) | 0.0433 | 0.0005 | 0.1368 | 0.0075 | 0.2654 | 0.3824 | 0.0825 | 0.1471 | 0.0004 | 0.0181 | 0.1381 | 0.0718 | 0.2580 | 0.0028 | 0.0016 | 0.0023 | 0.9732 | | 0.8874 | 0.9385 | 0.8245 |
| Velocity (m/s) | 0.0141 | 0.0035 | 0.0435 | 0.0010 | 0.4082 | 0.4919 | 0.0434 | 0.2059 | 0.0215 | 0.0980 | 0.2572 | 0.2072 | 0.1708 | 0.0272 | 0.0072 | 0.0000 | 0.8446 | 0.8874 | | 0.9456 | 0.7130 |
| Thigmotaxis % | 0.0031 | 0.0068 | 0.0978 | 0.0033 | 0.2928 | 0.3504 | 0.1288 | 0.2180 | 0.0225 | 0.0129 | 0.1858 | 0.1064 | 0.2931 | 0.0374 | 0.0005 | 0.0025 | 0.8716 | 0.9385 | 0.9456 | | 0.6974 |
| Floating % | 0.0057 | 0.0395 | 0.2918 | 0.0624 | 0.3145 | 0.3125 | 0.0273 | 0.1614 | 0.0074 | 0.0114 | 0.2355 | 0.1768 | 0.1958 | 0.0000 | 0.1068 | 0.1236 | 0.9231 | 0.8245 | 0.7130 | 0.6974 | |

| Mean Squared Error | Weight | SVP Density | ICP Density | DCP Density | Total Retina | IRL | RPE | RNFL | IPL | INL | ORL | PRC | OPL | Number of Target Zone Crossings | % of time in sector with platform | Latency on Day 3 | Latency on Day 4 | Distance swam (m) | Velocity (m/s) | Thigmotaxis % | Floating % |
|---|---|---|---|---|---|---|---|---|---|---|---|---|---|---|---|---|---|---|---|---|---|
| Weight | | 1.86E-05 | 0.0001814 | 0.0001418 | 29.675997 | 4.67889 | 0.57557 | 10.6256 | 3.68647 | 1.46096 | 11.0936 | 8.52552 | 0.73806 | 2.113372093 | 148.5595 | 53.7531001 | 277.082605 | 160362.109 | 14.94103 | 644.8711687 | 87.1992 |
| SVP Density | 1.86E-05 | | 0.0001329 | 0.00015148 | 12.070748 | 3.05926 | 0.60711 | 6.76741 | 1.98446 | 0.36865 | 3.05505 | 1.16954 | 0.74562 | 1.96895531 | 143.6274541 | 41.4134741 | 341.79219 | 200736.477 | 17.54716 | 770.8909933 | 97.2778 |
| ICP Density | 0.000181 | 0.000133 | | 0.00025719 | 10.202998 | 3.10396 | 0.36462 | 6.1447 | 4.49588 | 0.27712 | 4.33778 | 3.12139 | 0.29754 | 1.827987981 | 156.6650566 | 82.984163 | 283.490503 | 173361.784 | 16.84326 | 700.2202774 | 71.7211 |
| DCP Density | 0.000142 | 0.000151 | 0.0002572 | | 29.657571 | 5.66201 | 0.59388 | 12.5333 | 6.02154 | 2.31553 | 11.3783 | 7.88127 | 0.76999 | 2.201339193 | 98.5180404 | 52.1375152 | 286.124339 | 166371.285 | 15.13955 | 644.7653171 | 82.2229 |
| Total Retina | 29.676 | 12.07075 | 10.202998 | 29.6575714 | | 1.30648 | 0.58209 | 3.92219 | 3.61628 | 1.62266 | 1.30648 | 1.30189 | 0.58966 | 2.393276457 | 179.7347383 | 129.892608 | 213.458238 | 123132.714 | 8.968528 | 457.456661 | 60.1131 |
| IRL | 4.678894 | 3.05926 | 3.1039553 | 5.66201169 | 1.3064792 | | 0.65943 | 8.8452 | 5.8239 | 0.97431 | 6.44454 | 4.75888 | 0.73083 | 2.178935698 | 142.2770154 | 110.974347 | 187.984069 | 103519.511 | 7.700075 | 420.2360444 | 60.2932 |
| RPE | 0.575567 | 0.607106 | 0.3646172 | 0.59388267 | 0.5820928 | 0.65943 | | 4.83937 | 3.15797 | 1.94548 | 9.34182 | 8.45545 | 0.07216 | 2.741004972 | 200.0895209 | 125.108555 | 273.812468 | 153798.782 | 14.49617 | 563.5627017 | 85.3031 |
| RNFL | 10.62555 | 6.767406 | 6.1447009 | 12.533276 | 3.9221925 | 8.8452 | 4.83937 | | 1.02757 | 2.34849 | 1.90376 | 3.3414 | 0.23439 | 2.65294271 | 197.4717367 | 121.063281 | 251.255589 | 142959.446 | 12.03451 | 505.8657561 | 73.5405 |
| IPL | 3.68647 | 1.984461 | 4.4958789 | 6.02153635 | 3.6162756 | 5.8239 | 3.15797 | 1.02757 | | 2.34216 | 3.636 | 3.92129 | 0.4412 | 2.82306419 | 200.8532532 | 112.529389 | 286.601204 | 167564.831 | 14.8294 | 632.3029515 | 87.0469 |
| INL | 1.46096 | 0.368649 | 0.2771236 | 2.31552883 | 1.6226644 | 0.97431 | 1.94548 | 2.34849 | 2.34216 | | 10.7138 | 6.83052 | 0.73091 | 2.473701295 | 116.0822197 | 80.3759017 | 282.492828 | 164594.753 | 13.6689 | 638.5284435 | 86.6931 |
| ORL | 11.09362 | 3.055052 | 4.3377794 | 11.3783118 | 1.3064793 | 6.44454 | 9.34182 | 1.90376 | 3.636 | 10.7138 | | 0.58929 | 0.52447 | 2.629246299 | 198.392309 | 121.547922 | 245.56624 | 144474.11 | 11.25616 | 526.6759691 | 67.0461 |
| PRC | 8.525522 | 1.169543 | 3.1213921 | 7.88126787 | 1.3018927 | 4.75888 | 8.45545 | 3.3414 | 3.92129 | 6.83052 | 0.58929 | | 0.7032 | 2.688798348 | 197.110215 | 124.512447 | 262.28635 | 155582.686 | 12.01415 | 578.0664446 | 72.1889 |
| OPL | 0.738057 | 0.745616 | 0.2975387 | 0.76999256 | 0.5896645 | 0.73083 | 0.07216 | 0.23439 | 0.4412 | 0.73091 | 0.52447 | 0.7032 | | 2.689434791 | 202.3293902 | 119.612939 | 226.060635 | 124382.756 | 12.56574 | 457.2888631 | 70.5284 |
| Number of Target Zone Crossings | 2.113372 | 1.968955 | 1.827988 | 2.20133919 | 2.3932765 | 2.17894 | 2.741 | 2.65294 | 2.82306 | 2.4737 | 2.62925 | 2.6888 | 2.68943 | | 166.0452846 | 115.093324 | 285.63714 | 167147.277 | 14.74294 | 622.6715478 | 87.6912 |
| % of time in sector with platform | 148.5595 | 143.6275 | 156.66506 | 98.5180404 | 179.73474 | 142.277 | 200.09 | 197.472 | 200.853 | 116.082 | 198.392 | 197.11 | 202.329 | 166.0452846 | | 25.7624278 | 278.634773 | 167351.38 | 15.04489 | 646.5601852 | 78.3311 |
| Latency on Day 3 | 53.7531 | 41.41347 | 82.984163 | 52.1375152 | 129.89261 | 110.974 | 125.109 | 121.063 | 112.529 | 80.3759 | 121.548 | 124.512 | 119.613 | 115.0933241 | 25.76242782 | | 279.432413 | 167231.47 | 15.15438 | 645.2622457 | 76.8585 |
| Latency on Day 4 | 277.0826 | 341.7922 | 283.4905 | 286.124339 | 213.45824 | 187.984 | 273.812 | 251.256 | 286.601 | 282.493 | 245.566 | 262.286 | 226.061 | 285.6371396 | 278.6347729 | 279.432413 | | 4493.33792 | 2.354646 | 83.05890433 | 6.74114 |
| Distance swam (m) | 160362.1 | 200736.5 | 173361.78 | 166371.285 | 123132.71 | 103520 | 153799 | 142959 | 167565 | 164595 | 144474 | 155583 | 124383 | 167147.2765 | 167351.3802 | 167231.47 | 4493.33792 | | 1.706364 | 39.80308986 | 15.3945 |
| Velocity (m/s) | 14.94103 | 17.54716 | 16.843255 | 15.13955 | 8.9685275 | 7.70007 | 14.4962 | 12.0345 | 14.8294 | 13.6689 | 11.2562 | 12.0141 | 12.5657 | 14.74294464 | 15.04488511 | 15.15438 | 2.35464646 | 1.70636392 | | 35.21030704 | 25.1707 |
| Thigmotaxis % | 644.8712 | 770.891 | 700.22028 | 644.765317 | 457.45666 | 420.236 | 563.563 | 505.866 | 632.303 | 638.528 | 526.676 | 578.066 | 457.289 | 622.6715478 | 646.5601852 | 645.262246 | 83.0589043 | 39.8030899 | 35.21031 | | 26.5353 |
| Floating % | 87.1992 | 97.27783 | 71.721079 | 82.2229083 | 60.113118 | 60.2932 | 85.3031 | 73.5405 | 87.0469 | 86.6931 | 67.0461 | 72.1889 | 70.5284 | 87.69124506 | 78.33110984 | 76.8585355 | 6.74114249 | 15.3944811 | 25.1707 | 26.53533702 | |

Table 6: R-squared and corresponding MSE for or all tested parameters for transgenic male animals. For the R-squared values red indicates values from 0-0.3, yellow =0.4-0.6 and green 0.6-1.

| Correlation Coefficient | Weight | SVP Density | ICP Density | DCP Density | Total Retina | IRL | RPE | RNFL | IPL | INL | ORL | PRC | OPL | Number of Target Zone Crossings | % of time in sector with platform | Latency on Day 3 | Latency on Day 4 | Distance swam (m) | Velocity (m/s) | Thigmotaxis % | Floating % |
|---|---|---|---|---|---|---|---|---|---|---|---|---|---|---|---|---|---|---|---|---|---|
| Weight | | -0.3831 | -0.5340 | 0.0413 | 0.1072 | 0.1121 | -0.0039 | 0.1159 | -0.0147 | -0.0704 | 0.0925 | 0.0465 | 0.2940 | 0.1706 | 0.0857 | -0.3058 | 0.4759 | 0.2756 | -0.2061 | 0.4310 | 0.1287 |
| SVP Density | -0.3831 | | 0.1815 | 0.0434 | 0.0866 | 0.2714 | 0.2938 | 0.1296 | 0.0984 | 0.0465 | -0.0590 | -0.0872 | 0.1353 | 0.1617 | -0.0057 | -0.2025 | -0.1877 | 0.1667 | 0.5825 | -0.0843 | -0.0599 |
| ICP Density | -0.5340 | 0.1815 | | 0.4147 | -0.0548 | 0.0730 | -0.0416 | -0.1645 | 0.1633 | 0.3737 | -0.1436 | -0.0655 | -0.4934 | -0.3925 | -0.5363 | 0.2955 | -0.3655 | -0.2691 | 0.1663 | -0.4188 | -0.0170 |
| DCP Density | 0.0413 | 0.0434 | 0.4147 | | 0.4611 | 0.5228 | 0.2156 | 0.2735 | -0.0844 | 0.5307 | 0.3676 | 0.3873 | 0.0317 | -0.2399 | -0.2850 | 0.2752 | -0.1121 | 0.0897 | 0.3523 | -0.2443 | -0.3839 |
| Total Retina | 0.1072 | 0.0866 | -0.0548 | 0.4611 | | 0.9183 | 0.5124 | 0.8852 | -0.6970 | 0.7088 | 0.9563 | 0.9412 | 0.4546 | -0.3674 | -0.3715 | 0.0340 | 0.0041 | -0.0350 | -0.1097 | -0.2272 | -0.0200 |
| IRL | 0.1121 | 0.2714 | 0.0730 | 0.5228 | 0.9183 | | 0.5685 | 0.8547 | -0.5797 | 0.7732 | 0.7624 | 0.7461 | 0.3860 | -0.3093 | -0.4169 | 0.1028 | 0.0605 | -0.0266 | -0.1175 | -0.0994 | 0.0268 |
| RPE | -0.0039 | 0.2938 | -0.0416 | 0.2156 | 0.5124 | 0.5685 | | 0.7362 | -0.5423 | 0.0632 | 0.4176 | 0.3033 | 0.8027 | -0.5218 | -0.5653 | 0.0265 | 0.3210 | 0.3660 | 0.0960 | 0.1963 | -0.2810 |
| RNFL | 0.1159 | 0.1296 | -0.1645 | 0.2735 | 0.8852 | 0.8547 | 0.7362 | | -0.8845 | 0.5565 | 0.8155 | 0.7661 | 0.5927 | -0.4615 | -0.4508 | 0.0660 | 0.2554 | 0.1002 | -0.2650 | -0.0162 | 0.0952 |
| IPL | -0.0147 | 0.0984 | 0.1633 | -0.0844 | -0.6970 | ##### | -0.5423 | -0.8845 | | -0.4899 | -0.7112 | -0.6911 | -0.3879 | 0.5236 | 0.3813 | -0.0943 | -0.2372 | 0.0076 | 0.4621 | 0.1244 | -0.2338 |
| INL | -0.0704 | 0.0465 | 0.3737 | 0.5307 | 0.7088 | 0.7732 | 0.0632 | 0.5565 | -0.4899 | | 0.5874 | 0.6651 | -0.2080 | -0.2753 | -0.2920 | 0.2153 | -0.2106 | -0.3955 | -0.3260 | -0.4231 | 0.2108 |
| ORL | 0.0925 | -0.0590 | -0.1436 | 0.3676 | 0.9563 | 0.7624 | 0.4176 | 0.8155 | -0.7112 | 0.5874 | | 0.9874 | 0.4579 | -0.3721 | -0.2993 | -0.0217 | -0.0400 | -0.0392 | -0.0966 | -0.3120 | -0.0551 |
| PRC | 0.0465 | -0.0872 | -0.0655 | 0.3873 | 0.9412 | 0.7461 | 0.3033 | 0.7661 | -0.6911 | 0.6651 | 0.9874 | | 0.3112 | -0.3362 | -0.2688 | -0.0078 | -0.1361 | -0.1609 | -0.1444 | -0.4084 | 0.0226 |
| OPL | 0.2940 | 0.1353 | -0.4934 | 0.0317 | 0.4546 | 0.3860 | 0.8027 | 0.5927 | -0.3879 | -0.2080 | 0.4579 | 0.3112 | | -0.3452 | -0.2871 | -0.0819 | 0.4950 | 0.6312 | 0.2176 | 0.3947 | -0.4336 |
| Number of Target Zone Crossings | 0.1706 | 0.1617 | -0.3925 | -0.2399 | -0.3674 | ##### | -0.5218 | -0.4615 | 0.5236 | -0.2753 | -0.3721 | -0.3362 | -0.3452 | | 0.8461 | -0.4649 | -0.3560 | -0.3593 | 0.0193 | 0.0159 | 0.2382 |
| % of time in sector with platform | 0.0857 | -0.0057 | -0.5363 | -0.2850 | -0.3715 | ##### | -0.5653 | -0.4508 | 0.3813 | -0.2920 | -0.2993 | -0.2688 | -0.2871 | 0.8461 | | -0.3971 | -0.3912 | -0.3051 | 0.0829 | -0.1560 | -0.0158 |
| Latency on Day 3 | -0.3058 | -0.2025 | 0.2955 | 0.2752 | 0.0340 | 0.1028 | 0.0265 | 0.0660 | -0.0943 | 0.2153 | -0.0217 | -0.0078 | -0.0819 | -0.4649 | -0.3971 | | 0.4265 | 0.3401 | -0.0829 | 0.3573 | -0.0443 |
| Latency on Day 4 | 0.4759 | -0.1877 | -0.3655 | -0.1121 | 0.0041 | 0.0605 | 0.3210 | 0.2554 | -0.2372 | -0.2106 | -0.0400 | -0.1361 | 0.4950 | -0.3560 | -0.3912 | 0.4265 | | 0.7887 | -0.2114 | 0.8487 | 0.0952 |
| Distance swam (m) | 0.2756 | 0.1667 | -0.2691 | 0.0897 | -0.0350 | ##### | 0.3660 | 0.1002 | 0.0076 | -0.3955 | -0.0392 | -0.1609 | 0.6312 | -0.3593 | -0.3051 | 0.3401 | 0.7887 | | 0.4166 | 0.6472 | -0.3781 |
| Velocity (m/s) | -0.2061 | 0.5825 | 0.1663 | 0.3523 | -0.1097 | ##### | 0.0960 | -0.2650 | 0.4621 | -0.3260 | -0.0966 | -0.1444 | 0.2176 | 0.0193 | 0.0829 | -0.0829 | -0.2114 | 0.4166 | | -0.1815 | -0.6838 |
| Thigmotaxis % | 0.4310 | -0.0843 | -0.4188 | -0.2443 | -0.2272 | ##### | 0.1963 | -0.0162 | 0.1244 | -0.4231 | -0.3120 | -0.4084 | 0.3947 | 0.0159 | -0.1560 | 0.3573 | 0.8487 | 0.6472 | -0.1815 | | 0.0983 |
| Floating % | 0.1287 | -0.0599 | -0.0170 | -0.3839 | -0.0200 | 0.0268 | -0.2810 | 0.0952 | -0.2338 | 0.2108 | -0.0551 | 0.0226 | -0.4336 | 0.2382 | -0.0158 | -0.0443 | 0.0952 | -0.3781 | -0.6838 | 0.0983 | |

| P-Value | Weight | SVP Density | ICP Density | DCP Density | Total Retina | IRL | RPE | RNFL | IPL | INL | ORL | PRC | OPL | Number of Target Zone Crossings | % of time in sector with platform | Latency on Day 3 | Latency on Day 4 | Distance swam (m) | Velocity (m/s) | Thigmotaxis % | Floating % |
|---|---|---|---|---|---|---|---|---|---|---|---|---|---|---|---|---|---|---|---|---|---|
| Weight | | 0.1963 | 0.0601 | 0.8935 | 0.7273 | 0.7154 | 0.9899 | 0.7061 | 0.9620 | 0.8193 | 0.7637 | 0.8802 | 0.3295 | 0.5775 | 0.7806 | 0.3338 | 0.1179 | 0.3859 | 0.5205 | 0.1618 | 0.6903 |
| SVP Density | 0.1963 | | 0.5530 | 0.8879 | 0.7785 | 0.3697 | 0.3299 | 0.6729 | 0.7492 | 0.8802 | 0.8482 | 0.7770 | 0.6593 | 0.5976 | 0.9853 | 0.5279 | 0.5591 | 0.6045 | 0.0469 | 0.7945 | 0.8534 |
| ICP Density | 0.0601 | 0.5530 | | 0.1588 | 0.8588 | 0.8127 | 0.8926 | 0.5912 | 0.5940 | 0.2084 | 0.6398 | 0.8317 | 0.0866 | 0.1846 | 0.0588 | 0.3511 | 0.2427 | 0.3977 | 0.6056 | 0.1754 | 0.9582 |
| DCP Density | 0.8935 | 0.8879 | 0.1588 | | 0.1128 | 0.0668 | 0.4792 | 0.3659 | 0.7839 | 0.0621 | 0.2166 | 0.1911 | 0.9182 | 0.4298 | 0.3452 | 0.3866 | 0.7286 | 0.7815 | 0.2614 | 0.4441 | 0.2179 |
| Total Retina | 0.7273 | 0.7785 | 0.8588 | 0.1128 | | 0.0000 | 0.0734 | 0.0001 | 0.0081 | 0.0067 | 0.0000 | 0.0000 | 0.1186 | 0.2168 | 0.2114 | 0.9165 | 0.9899 | 0.9141 | 0.7344 | 0.4777 | 0.9508 |
| IRL | 0.7154 | 0.3697 | 0.8127 | 0.0668 | 0.0000 | | 0.0426 | 0.0002 | 0.0379 | 0.0019 | 0.0024 | 0.0034 | 0.1927 | 0.3037 | 0.1564 | 0.7505 | 0.8518 | 0.9346 | 0.7162 | 0.7585 | 0.9342 |
| RPE | 0.9899 | 0.3299 | 0.8926 | 0.4792 | 0.0734 | 0.0426 | | 0.0041 | 0.0556 | 0.8376 | 0.1556 | 0.3137 | 0.0010 | 0.0674 | 0.0441 | 0.9349 | 0.3091 | 0.2420 | 0.7665 | 0.5408 | 0.3763 |
| RNFL | 0.7061 | 0.6729 | 0.5912 | 0.3659 | 0.0001 | 0.0002 | 0.0041 | | 0.0001 | 0.0482 | 0.0007 | 0.0023 | 0.0328 | 0.1124 | 0.1221 | 0.8384 | 0.4231 | 0.7566 | 0.4052 | 0.9600 | 0.7685 |
| IPL | 0.9620 | 0.7492 | 0.5940 | 0.7839 | 0.0081 | 0.0379 | 0.0556 | 0.0001 | | 0.0893 | 0.0064 | 0.0089 | 0.1903 | 0.0663 | 0.1986 | 0.7708 | 0.4579 | 0.9812 | 0.1304 | 0.7001 | 0.4645 |
| INL | 0.8193 | 0.8802 | 0.2084 | 0.0621 | 0.0067 | 0.0019 | 0.8376 | 0.0482 | 0.0893 | | 0.0348 | 0.0131 | 0.4952 | 0.3626 | 0.3330 | 0.5015 | 0.5112 | 0.2032 | 0.3010 | 0.1706 | 0.5107 |
| ORL | 0.7637 | 0.8482 | 0.6398 | 0.2166 | 0.0000 | 0.0024 | 0.1556 | 0.0007 | 0.0064 | 0.0348 | | 0.0000 | 0.1156 | 0.2106 | 0.3205 | 0.9466 | 0.9017 | 0.9036 | 0.7653 | 0.3235 | 0.8649 |
| PRC | 0.8802 | 0.7770 | 0.8317 | 0.1911 | 0.0000 | 0.0034 | 0.3137 | 0.0023 | 0.0089 | 0.0131 | 0.0000 | | 0.3007 | 0.2614 | 0.3746 | 0.9808 | 0.6733 | 0.6175 | 0.6544 | 0.1875 | 0.9444 |
| OPL | 0.3295 | 0.6593 | 0.0866 | 0.9182 | 0.1186 | 0.1927 | 0.0010 | 0.0328 | 0.1903 | 0.4952 | 0.1156 | 0.3007 | | 0.2480 | 0.3416 | 0.8002 | 0.1018 | 0.0277 | 0.4968 | 0.2042 | 0.1591 |
| Number of Target Zone Crossings | 0.5775 | 0.5976 | 0.1846 | 0.4298 | 0.2168 | 0.3037 | 0.0674 | 0.1124 | 0.0663 | 0.3626 | 0.2106 | 0.2614 | 0.2480 | | 0.0003 | 0.1278 | 0.2560 | 0.2514 | 0.9525 | 0.9608 | 0.4559 |
| % of time in sector with platform | 0.7806 | 0.9853 | 0.0588 | 0.3452 | 0.2114 | 0.1564 | 0.0441 | 0.1221 | 0.1986 | 0.3330 | 0.3205 | 0.3746 | 0.3416 | 0.0003 | | 0.2012 | 0.2086 | 0.3349 | 0.7979 | 0.6282 | 0.9612 |
| Latency on Day 3 | 0.3338 | 0.5279 | 0.3511 | 0.3866 | 0.9165 | 0.7505 | 0.9349 | 0.8384 | 0.7708 | 0.5015 | 0.9466 | 0.9808 | 0.8002 | 0.1278 | 0.2012 | | 0.1668 | 0.2793 | 0.7979 | 0.2542 | 0.8913 |
| Latency on Day 4 | 0.1179 | 0.5591 | 0.2427 | 0.7286 | 0.9899 | 0.8518 | 0.3091 | 0.4231 | 0.4579 | 0.5112 | 0.9017 | 0.6733 | 0.1018 | 0.2560 | 0.2086 | 0.1668 | | 0.0023 | 0.5096 | 0.0005 | 0.7685 |
| Distance swam (m) | 0.3859 | 0.6045 | 0.3977 | 0.7815 | 0.9141 | 0.9346 | 0.2420 | 0.7566 | 0.9812 | 0.2032 | 0.9036 | 0.6175 | 0.0277 | 0.2514 | 0.3349 | 0.2793 | 0.0023 | | 0.1779 | 0.0229 | 0.2256 |
| Velocity (m/s) | 0.5205 | 0.0469 | 0.6056 | 0.2614 | 0.7344 | 0.7162 | 0.7665 | 0.4052 | 0.1304 | 0.3010 | 0.7653 | 0.6544 | 0.4968 | 0.9525 | 0.7979 | 0.7979 | 0.5096 | 0.1779 | | 0.5724 | 0.0142 |
| Thigmotaxis % | 0.1618 | 0.7945 | 0.1754 | 0.4441 | 0.4777 | 0.7585 | 0.5408 | 0.9600 | 0.7001 | 0.1706 | 0.3235 | 0.1875 | 0.2042 | 0.9608 | 0.6282 | 0.2542 | 0.0005 | 0.0229 | 0.5724 | | 0.7611 |
| Floating % | 0.6903 | 0.8534 | 0.9582 | 0.2179 | 0.9508 | 0.9342 | 0.3763 | 0.7685 | 0.4645 | 0.5107 | 0.8649 | 0.9444 | 0.1591 | 0.4559 | 0.9612 | 0.8913 | 0.7685 | 0.2256 | 0.0142 | 0.7611 | |

Table 7: Correlation Coefficient and corresponding p-value for or all tested parameters for all non-transgenic animals. For the Correlation coefficient red indicates values from 0-0.3, yellow =0.3-0.5 and green 0.5-1. For the p-value red indicates values above 0.15, yellow values from 0.15-0.1 and green values from below 0.05.

| R-Squared | Weight | SVP Density | ICP Density | DCP Density | Total Retina | IRL | RPE | RNFL | IPL | INL | ORL | PRC | OPL | Number of Target Zone Crossings | % of time in sector with platform | Latency on Day 3 | Latency on Day 4 | Distance swam (m) | Velocity (m/s) | Thigmotaxis % | Floating % |
|---|---|---|---|---|---|---|---|---|---|---|---|---|---|---|---|---|---|---|---|---|---|
| Weight | | 0.1468 | 0.2852 | 0.0017 | 0.0115 | 0.0126 | 0.0000 | 0.0134 | 0.0002 | 0.0050 | 0.0086 | 0.0022 | 0.0864 | 0.0291 | 0.0074 | 0.0935 | 0.2265 | 0.0760 | 0.0425 | 0.1858 | 0.0166 |
| SVP Density | 0.1468 | | 0.0329 | 0.0019 | 0.0075 | 0.0737 | 0.0863 | 0.0168 | 0.0097 | 0.0022 | 0.0035 | 0.0076 | 0.0183 | 0.0262 | 0.0000 | 0.0410 | 0.0352 | 0.0278 | 0.3393 | 0.0071 | 0.0036 |
| ICP Density | 0.2852 | 0.0329 | | 0.1720 | 0.0030 | 0.0053 | 0.0017 | 0.0271 | 0.0267 | 0.1397 | 0.0206 | 0.0043 | 0.2435 | 0.1541 | 0.2876 | 0.0873 | 0.1336 | 0.0724 | 0.0276 | 0.1754 | 0.0003 |
| DCP Density | 0.0017 | 0.0019 | 0.1720 | | 0.2126 | 0.2733 | 0.0465 | 0.0748 | 0.0071 | 0.2816 | 0.1351 | 0.1500 | 0.0010 | 0.0576 | 0.0812 | 0.0758 | 0.0126 | 0.0081 | 0.1241 | 0.0597 | 0.1474 |
| Total Retina | 0.0115 | 0.0075 | 0.0030 | 0.2126 | | 0.8433 | 0.2626 | 0.7835 | 0.4859 | 0.5024 | 0.9144 | 0.8859 | 0.2066 | 0.1350 | 0.1380 | 0.0012 | 0.0000 | 0.0012 | 0.0120 | 0.0516 | 0.0004 |
| IRL | 0.0126 | 0.0737 | 0.0053 | 0.2733 | 0.8433 | | 0.3232 | 0.7305 | 0.3360 | 0.5979 | 0.5812 | 0.5567 | 0.1490 | 0.0957 | 0.1738 | 0.0106 | 0.0037 | 0.0007 | 0.0138 | 0.0099 | 0.0007 |
| RPE | 0.0000 | 0.0863 | 0.0017 | 0.0465 | 0.2626 | 0.3232 | | 0.5420 | 0.2940 | 0.0040 | 0.1744 | 0.0920 | 0.6443 | 0.2723 | 0.3196 | 0.0007 | 0.1030 | 0.1340 | 0.0092 | 0.0386 | 0.0790 |
| RNFL | 0.0134 | 0.0168 | 0.0271 | 0.0748 | 0.7835 | 0.7305 | 0.5420 | | 0.7824 | 0.3097 | 0.6651 | 0.5869 | 0.3512 | 0.2130 | 0.2032 | 0.0044 | 0.0652 | 0.0100 | 0.0702 | 0.0003 | 0.0091 |
| IPL | 0.0002 | 0.0097 | 0.0267 | 0.0071 | 0.4859 | 0.3360 | 0.2940 | 0.7824 | | 0.2400 | 0.5058 | 0.4777 | 0.1505 | 0.2742 | 0.1454 | 0.0089 | 0.0563 | 0.0001 | 0.2135 | 0.0155 | 0.0547 |
| INL | 0.0050 | 0.0022 | 0.1397 | 0.2816 | 0.5024 | 0.5979 | 0.0040 | 0.3097 | 0.2400 | | 0.3451 | 0.4423 | 0.0433 | 0.0758 | 0.0853 | 0.0464 | 0.0444 | 0.1564 | 0.1063 | 0.1790 | 0.0444 |
| ORL | 0.0086 | 0.0035 | 0.0206 | 0.1351 | 0.9144 | 0.5812 | 0.1744 | 0.6651 | 0.5058 | 0.3451 | | 0.9749 | 0.2097 | 0.1385 | 0.0896 | 0.0005 | 0.0016 | 0.0015 | 0.0093 | 0.0973 | 0.0030 |
| PRC | 0.0022 | 0.0076 | 0.0043 | 0.1500 | 0.8859 | 0.5567 | 0.0920 | 0.5869 | 0.4777 | 0.4423 | 0.9749 | | 0.0968 | 0.1130 | 0.0722 | 0.0001 | 0.0185 | 0.0259 | 0.0208 | 0.1668 | 0.0005 |
| OPL | 0.0864 | 0.0183 | 0.2435 | 0.0010 | 0.2066 | 0.1490 | 0.6443 | 0.3512 | 0.1505 | 0.0433 | 0.2097 | 0.0968 | | 0.1192 | 0.0824 | 0.0067 | 0.2451 | 0.3984 | 0.0474 | 0.1558 | 0.1880 |
| Number of Target Zone Crossings | 0.0291 | 0.0262 | 0.1541 | 0.0576 | 0.1350 | 0.0957 | 0.2723 | 0.2130 | 0.2742 | 0.0758 | 0.1385 | 0.1130 | 0.1192 | | 0.7160 | 0.2161 | 0.1268 | 0.1291 | 0.0004 | 0.0003 | 0.0567 |
| % of time in sector with platform | 0.0074 | 0.0000 | 0.2876 | 0.0812 | 0.1380 | 0.1738 | 0.3196 | 0.2032 | 0.1454 | 0.0853 | 0.0896 | 0.0722 | 0.0824 | 0.7160 | | 0.1577 | 0.1530 | 0.0931 | 0.0069 | 0.0243 | 0.0002 |
| Latency on Day 3 | 0.0935 | 0.0410 | 0.0873 | 0.0758 | 0.0012 | 0.0106 | 0.0007 | 0.0044 | 0.0089 | 0.0464 | 0.0005 | 0.0001 | 0.0067 | 0.2161 | 0.1577 | | 0.1819 | 0.1157 | 0.0069 | 0.1276 | 0.0020 |
| Latency on Day 4 | 0.2265 | 0.0352 | 0.1336 | 0.0126 | 0.0000 | 0.0037 | 0.1030 | 0.0652 | 0.0563 | 0.0444 | 0.0016 | 0.0185 | 0.2451 | 0.1268 | 0.1530 | 0.1819 | | 0.6221 | 0.0447 | 0.7202 | 0.0091 |
| Distance swam (m) | 0.0760 | 0.0278 | 0.0724 | 0.0081 | 0.0012 | 0.0007 | 0.1340 | 0.0100 | 0.0001 | 0.1564 | 0.0015 | 0.0259 | 0.3984 | 0.1291 | 0.0931 | 0.1157 | 0.6221 | | 0.1736 | 0.4188 | 0.1430 |
| Velocity (m/s) | 0.0425 | 0.3393 | 0.0276 | 0.1241 | 0.0120 | 0.0138 | 0.0092 | 0.0702 | 0.2135 | 0.1063 | 0.0093 | 0.0208 | 0.0474 | 0.0004 | 0.0069 | 0.0069 | 0.0447 | 0.1736 | | 0.0329 | 0.4676 |
| Thigmotaxis % | 0.1858 | 0.0071 | 0.1754 | 0.0597 | 0.0516 | 0.0099 | 0.0386 | 0.0003 | 0.0155 | 0.1790 | 0.0973 | 0.1668 | 0.1558 | 0.0003 | 0.0243 | 0.1276 | 0.7202 | 0.4188 | 0.0329 | | 0.0097 |
| Floating % | 0.0166 | 0.0036 | 0.0003 | 0.1474 | 0.0004 | 0.0007 | 0.0790 | 0.0091 | 0.0547 | 0.0444 | 0.0030 | 0.0005 | 0.1880 | 0.0567 | 0.0002 | 0.0020 | 0.0091 | 0.1430 | 0.4676 | 0.0097 | |

| Mean Squared Error | Weight | SVP Density | ICP Density | DCP Density | Total Retina | IRL | RPE | RNFL | IPL | INL | ORL | PRC | OPL | Number of Target Zone Crossings | % of time in sector with platform | Latency on Day 3 | Latency on Day 4 | Distance swam (m) | Velocity (m/s) | Thigmotaxis % | Floating % |
|---|---|---|---|---|---|---|---|---|---|---|---|---|---|---|---|---|---|---|---|---|---|
| Weight | | 0.00016 | 0.00041 | 0.000274 | 76.5299 | 15.617 | 0.59757 | 27.5526 | 10.2599 | 2.88649 | 28.7205 | 25.2942 | 0.6554 | 4.986682692 | 479.5229385 | 148.80482 | 98.401248 | 28030.8751 | 7.132828 | 115.0455861 | 75.2779 |
| SVP Density | 0.00016 | | 0.00056 | 0.000274 | 76.8398 | 14.651 | 0.546 | 27.4586 | 10.1628 | 2.89459 | 28.8676 | 25.1562 | 0.70427 | 5.001757528 | 483.0586565 | 157.42194 | 122.72754 | 29491.6385 | 4.921511 | 140.2907414 | 76.2708 |
| ICP Density | 0.000412 | 0.00056 | | 0.000228 | 77.1875 | 15.731 | 0.59655 | 27.1721 | 9.98856 | 2.4957 | 28.3712 | 25.2402 | 0.54275 | 4.344833467 | 344.1300649 | 149.81971 | 110.21547 | 28138.5552 | 7.243281 | 116.5125935 | 76.5229 |
| DCP Density | 0.000274 | 0.00027 | 0.00023 | | 60.9594 | 11.494 | 0.56979 | 25.8384 | 10.189 | 2.08385 | 25.0549 | 21.5471 | 0.7167 | 4.840483864 | 443.831838 | 151.71823 | 125.60926 | 30090.7867 | 6.524524 | 132.8605262 | 65.2613 |
| Total Retina | 76.52993 | 76.8398 | 77.1875 | 60.95938 | | 2.4783 | 0.44066 | 6.04653 | 5.27599 | 1.44341 | 2.47829 | 2.89334 | 0.56917 | 4.44273019 | 416.4012432 | 163.96332 | 127.20634 | 30298.0036 | 7.359605 | 134.0028339 | 76.5144 |
| IRL | 15.61694 | 14.6506 | 15.7314 | 11.49356 | 2.47829 | | 0.40442 | 7.52585 | 6.81394 | 1.16649 | 12.1317 | 11.2362 | 0.61052 | 4.644610404 | 399.1150958 | 162.41745 | 126.7427 | 30313.6313 | 7.346396 | 139.8980756 | 76.4901 |
| RPE | 0.597574 | 0.546 | 0.59655 | 0.569793 | 0.44066 | 0.4044 | | 12.7915 | 7.24461 | 2.88927 | 23.9155 | 23.0166 | 0.25519 | 3.737598327 | 328.7024015 | 164.03791 | 114.10466 | 26271.1053 | 7.380469 | 135.8473944 | 70.5007 |
| RNFL | 27.55263 | 27.4586 | 27.1721 | 25.83842 | 6.04653 | 7.5258 | 12.7915 | | 2.23342 | 2.00236 | 9.70263 | 10.4711 | 0.46543 | 4.042320943 | 384.9198214 | 163.43699 | 118.9118 | 30030.3375 | 6.926043 | 141.2572568 | 75.8509 |
| IPL | 10.25993 | 10.1628 | 9.98856 | 10.189 | 5.27599 | 6.8139 | 7.24461 | 2.23342 | | 2.20472 | 14.3148 | 13.2403 | 0.60947 | 3.727811545 | 412.8319826 | 162.69445 | 120.05226 | 30333.3043 | 5.858556 | 139.1080189 | 72.3603 |
| INL | 2.886489 | 2.89459 | 2.4957 | 2.083855 | 1.44341 | 1.1665 | 2.88927 | 2.00236 | 2.20472 | | 18.9718 | 14.1362 | 0.68636 | 4.746789699 | 441.8813846 | 156.54088 | 121.56586 | 25590.0178 | 6.657296 | 116.0025838 | 73.1428 |
| ORL | 28.72048 | 28.8676 | 28.3712 | 25.05489 | 2.47829 | 12.132 | 23.9155 | 9.70263 | 14.3148 | 18.9718 | | 0.63706 | 0.56698 | 4.424992004 | 439.7997696 | 164.07567 | 127.00467 | 30288.3834 | 7.379705 | 127.54 | 76.3124 |
| PRC | 25.29419 | 25.1562 | 25.2402 | 21.54708 | 2.89334 | 11.236 | 23.0166 | 10.4711 | 13.2403 | 14.1362 | 0.63706 | | 0.64794 | 4.555529353 | 448.1789808 | 164.143 | 124.85342 | 29550.2089 | 7.293909 | 117.7318306 | 76.5059 |
| OPL | 0.655398 | 0.70427 | 0.54275 | 0.716697 | 0.56917 | 0.6105 | 0.25519 | 0.46543 | 0.60947 | 0.68636 | 0.56698 | 0.64794 | | 4.523987687 | 443.2676434 | 163.05148 | 96.034962 | 18249.189 | 7.096365 | 119.2821626 | 62.156 |
| Number of Target Zone Crossings | 4.986683 | 5.00176 | 4.34483 | 4.840484 | 4.44273 | 4.6446 | 3.7376 | 4.04232 | 3.72781 | 4.74679 | 4.42499 | 4.55553 | 4.52399 | | 137.2105843 | 128.67644 | 111.08245 | 26419.6133 | 7.446406 | 141.2586586 | 72.2013 |
| % of time in sector with platform | 479.5229 | 483.059 | 344.13 | 443.8318 | 416.401 | 399.12 | 328.702 | 384.92 | 412.832 | 441.881 | 439.8 | 448.179 | 443.268 | 137.2105843 | | 138.26403 | 107.74328 | 27511.4879 | 7.398029 | 137.8551199 | 76.526 |
| Latency on Day 3 | 148.8048 | 157.422 | 149.82 | 151.7182 | 163.963 | 162.42 | 164.038 | 163.437 | 162.694 | 156.541 | 164.076 | 164.143 | 163.051 | 128.6764417 | 138.2640283 | | 104.074 | 26825.2639 | 7.39804 | 123.2594164 | 76.3949 |
| Latency on Day 4 | 98.40125 | 122.728 | 110.215 | 125.6093 | 127.206 | 126.74 | 114.105 | 118.912 | 120.052 | 121.566 | 127.005 | 124.853 | 96.035 | 111.0824495 | 107.7432762 | 104.074 | | 11465.0993 | 7.116407 | 39.53110378 | 75.851 |
| Distance swam (m) | 28030.88 | 29491.6 | 28138.6 | 30090.79 | 30298 | 30314 | 26271.1 | 30030.3 | 30333.3 | 25590 | 30288.4 | 29550.2 | 18249.2 | 26419.61325 | 27511.48794 | 26825.264 | 11465.099 | | 6.156232 | 82.11457998 | 65.6024 |
| Velocity (m/s) | 7.132828 | 4.92151 | 7.24328 | 6.524524 | 7.35961 | 7.3464 | 7.38047 | 6.92604 | 5.85856 | 6.6573 | 7.3797 | 7.29391 | 7.09636 | 7.446405878 | 7.398028872 | 7.3980398 | 7.116407 | 6.1562316 | | 136.6407555 | 40.7526 |
| Thigmotaxis % | 115.0456 | 140.291 | 116.513 | 132.8605 | 134.003 | 139.9 | 135.847 | 141.257 | 139.108 | 116.003 | 127.54 | 117.732 | 119.282 | 141.2586586 | 137.8551199 | 123.25942 | 39.531104 | 82.11458 | 136.6408 | | 75.8049 |
| Floating % | 75.27794 | 76.2708 | 76.5229 | 65.26129 | 76.5144 | 76.49 | 70.5007 | 75.8509 | 72.3603 | 73.1428 | 76.3124 | 76.5059 | 62.156 | 72.2013215 | 76.52596058 | 76.394895 | 75.851 | 65.602432 | 40.75259 | 75.80488212 | |

Table 8: R-squared and corresponding MSE for or all tested parameters for all non-transgenic animals. For the R-squared values red indicates values from 0-0.3, yellow =0.4-0.6 and green 0.6-1.

| Correlation Coefficient | Weight | SVP Density | ICP Density | DCP Density | Total Retina | IRL | RPE | RNFL | IPL | INL | ORL | PRC | OPL | Number of Target Zone Crossings | % of time in sector with platform | Latency on Day 3 | Latency on Day 4 | Distance swam (m) | Velocity (m/s) | Thigmotaxis % | Floating % |
|---|---|---|---|---|---|---|---|---|---|---|---|---|---|---|---|---|---|---|---|---|---|
| Weight | | -0.0697 | -0.4180 | 0.0742 | 0.1425 | 0.1284 | 0.0354 | 0.1234 | 0.0021 | -0.0673 | 0.1506 | 0.0821 | 0.4116 | -0.1308 | -0.1546 | -0.2628 | 0.7773 | 0.6827 | 0.3230 | 0.4677 | -0.0677 |
| SVP Density | -0.0697 | | -0.6629 | -0.1896 | 0.4870 | 0.4458 | 0.4984 | 0.5684 | -0.3825 | -0.0220 | 0.5093 | 0.4241 | 0.6147 | 0.3256 | 0.3603 | -0.4007 | -0.1175 | 0.0338 | 0.2928 | -0.0184 | -0.1023 |
| ICP Density | -0.4180 | -0.6629 | | 0.4941 | ##### | 0.0603 | -0.1207 | -0.1070 | -0.0040 | 0.4627 | -0.0690 | 0.0366 | -0.5822 | -0.5783 | -0.6258 | 0.5159 | -0.4132 | -0.4658 | -0.3342 | -0.5138 | 0.0929 |
| DCP Density | 0.0742 | -0.1896 | 0.4941 | | 0.6410 | 0.7163 | 0.4235 | 0.5428 | -0.3972 | 0.7086 | 0.5654 | 0.5631 | 0.1935 | -0.7328 | -0.7500 | 0.5606 | 0.1413 | 0.1244 | 0.0604 | -0.1175 | -0.4549 |
| Total Retina | 0.1425 | 0.4870 | -0.0113 | 0.6410 | | 0.9844 | 0.5311 | 0.9322 | -0.7706 | 0.7852 | 0.9898 | 0.9723 | 0.4100 | -0.3521 | -0.3753 | -0.0338 | -0.1054 | -0.0820 | -0.0572 | -0.3684 | -0.0435 |
| IRL | 0.1284 | 0.4458 | 0.0603 | 0.7163 | 0.9844 | | 0.5426 | 0.8919 | -0.6770 | 0.7905 | 0.9493 | 0.9293 | 0.4106 | -0.4040 | -0.4332 | 0.0863 | -0.0799 | -0.0421 | 0.0443 | -0.3223 | -0.1538 |
| RPE | 0.0354 | 0.4984 | -0.1207 | 0.4235 | 0.5311 | 0.5426 | | 0.7545 | -0.6223 | 0.0401 | 0.5097 | 0.3829 | 0.8362 | -0.6014 | -0.5305 | -0.0266 | 0.2802 | 0.4003 | 0.5181 | 0.0635 | -0.5718 |
| RNFL | 0.1234 | 0.5684 | -0.1070 | 0.5428 | 0.9322 | 0.8919 | 0.7545 | | -0.8995 | 0.5760 | 0.9436 | 0.8872 | 0.6016 | -0.4456 | -0.4286 | -0.1749 | -0.0081 | 0.0369 | 0.0373 | -0.3140 | -0.1231 |
| IPL | 0.0021 | -0.3825 | -0.0040 | -0.3972 | ##### | -0.6770 | -0.6223 | -0.8995 | | -0.5319 | -0.8287 | -0.8072 | -0.3798 | 0.4342 | 0.4034 | 0.3203 | 0.1518 | 0.1609 | 0.2924 | 0.4973 | -0.1349 |
| INL | -0.0673 | -0.0220 | 0.4627 | 0.7086 | 0.7852 | 0.7905 | 0.0401 | 0.5760 | -0.5319 | | 0.7629 | 0.8479 | -0.2062 | -0.3255 | -0.3944 | 0.2035 | -0.4062 | -0.4652 | -0.4753 | -0.6241 | 0.2201 |
| ORL | 0.1506 | 0.5093 | -0.0690 | 0.5654 | 0.9898 | 0.9493 | 0.5097 | 0.9436 | -0.8287 | 0.7629 | | 0.9850 | 0.4002 | -0.3021 | -0.3198 | -0.1301 | -0.1236 | -0.1124 | -0.1381 | -0.3972 | 0.0468 |
| PRC | 0.0821 | 0.4241 | 0.0366 | 0.5631 | 0.9723 | 0.9293 | 0.3829 | 0.8872 | -0.8072 | 0.8479 | 0.9850 | | 0.2359 | -0.2696 | -0.3016 | -0.1072 | -0.2479 | -0.2564 | -0.2804 | -0.5094 | 0.1668 |
| OPL | 0.4116 | 0.6147 | -0.5822 | 0.1935 | 0.4100 | 0.4106 | 0.8362 | 0.6016 | -0.3798 | -0.2062 | 0.4002 | 0.2359 | | -0.2691 | -0.1990 | -0.1635 | 0.6199 | 0.7277 | 0.7105 | 0.4677 | -0.6218 |
| Number of Target Zone Crossings | -0.1308 | 0.3256 | -0.5783 | -0.7328 | ##### | -0.4040 | -0.6014 | -0.4456 | 0.4342 | -0.3255 | -0.3021 | -0.2696 | -0.2691 | | 0.9917 | -0.3430 | -0.2641 | -0.2474 | -0.2022 | 0.1073 | 0.3502 |
| % of time in sector with platform | -0.1546 | 0.3603 | -0.6258 | -0.7500 | ##### | -0.4332 | -0.5305 | -0.4286 | 0.4034 | -0.3944 | -0.3198 | -0.3016 | -0.1990 | 0.9917 | | -0.3648 | -0.2261 | -0.1994 | -0.1693 | 0.1481 | 0.2869 |
| Latency on Day 3 | -0.2628 | -0.4007 | 0.5159 | 0.5606 | ##### | 0.0863 | -0.0266 | -0.1749 | 0.3203 | 0.2035 | -0.1301 | -0.1072 | -0.1635 | -0.3430 | -0.3648 | | 0.0834 | 0.1175 | 0.2019 | 0.2850 | -0.4434 |
| Latency on Day 4 | 0.7773 | -0.1175 | -0.4132 | 0.1413 | ##### | -0.0799 | 0.2802 | -0.0081 | 0.1518 | -0.4062 | -0.1236 | -0.2479 | 0.6199 | -0.2641 | -0.2261 | 0.0834 | | 0.9754 | 0.6398 | 0.8380 | -0.5558 |
| Distance swam (m) | 0.6827 | 0.0338 | -0.4658 | 0.1244 | ##### | -0.0421 | 0.4003 | 0.0369 | 0.1609 | -0.4652 | -0.1124 | -0.2564 | 0.7277 | -0.2474 | -0.1994 | 0.1175 | 0.9754 | | 0.7776 | 0.8755 | -0.6660 |
| Velocity (m/s) | 0.3230 | 0.2928 | -0.3342 | 0.0604 | ##### | 0.0443 | 0.5181 | 0.0373 | 0.2924 | -0.4753 | -0.1381 | -0.2804 | 0.7105 | -0.2022 | -0.1693 | 0.2019 | 0.6398 | 0.7776 | | 0.6960 | -0.7515 |
| Thigmotaxis % | 0.4677 | -0.0184 | -0.5138 | -0.1175 | ##### | -0.3223 | 0.0635 | -0.3140 | 0.4973 | -0.6241 | -0.3972 | -0.5094 | 0.4677 | 0.1073 | 0.1481 | 0.2850 | 0.8380 | 0.8755 | 0.6960 | | -0.5757 |
| Floating % | -0.0677 | -0.1023 | 0.0929 | -0.4549 | ##### | -0.1538 | -0.5718 | -0.1231 | -0.1349 | 0.2201 | 0.0468 | 0.1668 | -0.6218 | 0.3502 | 0.2869 | -0.4434 | -0.5558 | -0.6660 | -0.7515 | -0.5757 | |

| P-Value | Weight | SVP Density | ICP Density | DCP Density | Total Retina | IRL | RPE | RNFL | IPL | INL | ORL | PRC | OPL | Number of Target Zone Crossings | % of time in sector with platform | Latency on Day 3 | Latency on Day 4 | Distance swam (m) | Velocity (m/s) | Thigmotaxis % | Floating % |
|---|---|---|---|---|---|---|---|---|---|---|---|---|---|---|---|---|---|---|---|---|---|
| Weight | | 0.8698 | 0.3027 | 0.8613 | 0.7365 | 0.7618 | 0.9336 | 0.7710 | 0.9961 | 0.8742 | 0.7219 | 0.8468 | 0.3110 | 0.7575 | 0.7148 | 0.5294 | 0.0232 | 0.0621 | 0.4352 | 0.2425 | 0.8734 |
| SVP Density | 0.8698 | | 0.0732 | 0.6529 | 0.2210 | 0.2683 | 0.2087 | 0.1415 | 0.3497 | 0.9588 | 0.1974 | 0.2950 | 0.1048 | 0.4313 | 0.3806 | 0.3252 | 0.7817 | 0.9366 | 0.4816 | 0.9654 | 0.8094 |
| ICP Density | 0.3027 | 0.0732 | | 0.2133 | 0.9788 | 0.8871 | 0.7758 | 0.8009 | 0.9925 | 0.2483 | 0.8710 | 0.9315 | 0.1300 | 0.1332 | 0.0970 | 0.1906 | 0.3089 | 0.2447 | 0.4184 | 0.1927 | 0.8269 |
| DCP Density | 0.8613 | 0.6529 | 0.2133 | | 0.0868 | 0.0456 | 0.2958 | 0.1645 | 0.3298 | 0.0491 | 0.1441 | 0.1461 | 0.6461 | 0.0387 | 0.0321 | 0.1483 | 0.7385 | 0.7691 | 0.8870 | 0.7818 | 0.2574 |
| Total Retina | 0.7365 | 0.2210 | 0.9788 | 0.0868 | | 0.0000 | 0.1756 | 0.0007 | 0.0252 | 0.0210 | 0.0000 | 0.0001 | 0.3131 | 0.3923 | 0.3596 | 0.9368 | 0.8038 | 0.8469 | 0.8929 | 0.3693 | 0.9186 |
| IRL | 0.7618 | 0.2683 | 0.8871 | 0.0456 | 0.0000 | | 0.1646 | 0.0029 | 0.0651 | 0.0195 | 0.0003 | 0.0008 | 0.3123 | 0.3209 | 0.2836 | 0.8390 | 0.8508 | 0.9211 | 0.9170 | 0.4362 | 0.7162 |
| RPE | 0.9336 | 0.2087 | 0.7758 | 0.2958 | 0.1756 | 0.1646 | | 0.0305 | 0.0994 | 0.9249 | 0.1969 | 0.3492 | 0.0097 | 0.1147 | 0.1762 | 0.9501 | 0.5014 | 0.3257 | 0.1884 | 0.8812 | 0.1386 |
| RNFL | 0.7710 | 0.1415 | 0.8009 | 0.1645 | 0.0007 | 0.0029 | 0.0305 | | 0.0023 | 0.1351 | 0.0004 | 0.0033 | 0.1146 | 0.2685 | 0.2893 | 0.6787 | 0.9849 | 0.9308 | 0.9301 | 0.4489 | 0.7715 |
| IPL | 0.9961 | 0.3497 | 0.9925 | 0.3298 | 0.0252 | 0.0651 | 0.0994 | 0.0023 | | 0.1748 | 0.0110 | 0.0154 | 0.3534 | 0.2824 | 0.3216 | 0.4393 | 0.7197 | 0.7034 | 0.4822 | 0.2098 | 0.7502 |
| INL | 0.8742 | 0.9588 | 0.2483 | 0.0491 | 0.0210 | 0.0195 | 0.9249 | 0.1351 | 0.1748 | | 0.0277 | 0.0078 | 0.6241 | 0.4314 | 0.3336 | 0.6288 | 0.3180 | 0.2455 | 0.2340 | 0.0981 | 0.6004 |
| ORL | 0.7219 | 0.1974 | 0.8710 | 0.1441 | 0.0000 | 0.0003 | 0.1969 | 0.0004 | 0.0110 | 0.0277 | | 0.0000 | 0.3259 | 0.4671 | 0.4400 | 0.7587 | 0.7706 | 0.7910 | 0.7443 | 0.3298 | 0.9124 |
| PRC | 0.8468 | 0.2950 | 0.9315 | 0.1461 | 0.0001 | 0.0008 | 0.3492 | 0.0033 | 0.0154 | 0.0078 | 0.0000 | | 0.5739 | 0.5185 | 0.4678 | 0.8006 | 0.5538 | 0.5400 | 0.5012 | 0.1972 | 0.6930 |
| OPL | 0.3110 | 0.1048 | 0.1300 | 0.6461 | 0.3131 | 0.3123 | 0.0097 | 0.1146 | 0.3534 | 0.6241 | 0.3259 | 0.5739 | | 0.5193 | 0.6365 | 0.6988 | 0.1011 | 0.0407 | 0.0482 | 0.2425 | 0.0998 |
| Number of Target Zone Crossings | 0.7575 | 0.4313 | 0.1332 | 0.0387 | 0.3923 | 0.3209 | 0.1147 | 0.2685 | 0.2824 | 0.4314 | 0.4671 | 0.5185 | 0.5193 | | 0.0000 | 0.4055 | 0.5274 | 0.5547 | 0.6311 | 0.8003 | 0.3951 |
| % of time in sector with platform | 0.7148 | 0.3806 | 0.0970 | 0.0321 | 0.3596 | 0.2836 | 0.1762 | 0.2893 | 0.3216 | 0.3336 | 0.4400 | 0.4678 | 0.6365 | 0.0000 | | 0.3743 | 0.5903 | 0.6358 | 0.6886 | 0.7263 | 0.4908 |
| Latency on Day 3 | 0.5294 | 0.3252 | 0.1906 | 0.1483 | 0.9368 | 0.8390 | 0.9501 | 0.6787 | 0.4393 | 0.6288 | 0.7587 | 0.8006 | 0.6988 | 0.4055 | 0.3743 | | 0.8444 | 0.7817 | 0.6316 | 0.4938 | 0.2711 |
| Latency on Day 4 | 0.0232 | 0.7817 | 0.3089 | 0.7385 | 0.8038 | 0.8508 | 0.5014 | 0.9849 | 0.7197 | 0.3180 | 0.7706 | 0.5538 | 0.1011 | 0.5274 | 0.5903 | 0.8444 | | 0.0000 | 0.0876 | 0.0094 | 0.1526 |
| Distance swam (m) | 0.0621 | 0.9366 | 0.2447 | 0.7691 | 0.8469 | 0.9211 | 0.3257 | 0.9308 | 0.7034 | 0.2455 | 0.7910 | 0.5400 | 0.0407 | 0.5547 | 0.6358 | 0.7817 | 0.0000 | | 0.0231 | 0.0044 | 0.0714 |
| Velocity (m/s) | 0.4352 | 0.4816 | 0.4184 | 0.8870 | 0.8929 | 0.9170 | 0.1884 | 0.9301 | 0.4822 | 0.2340 | 0.7443 | 0.5012 | 0.0482 | 0.6311 | 0.6886 | 0.6316 | 0.0876 | 0.0231 | | 0.0552 | 0.0316 |
| Thigmotaxis % | 0.2425 | 0.9654 | 0.1927 | 0.7818 | 0.3693 | 0.4362 | 0.8812 | 0.4489 | 0.2098 | 0.0981 | 0.3298 | 0.1972 | 0.2425 | 0.8003 | 0.7263 | 0.4938 | 0.0094 | 0.0044 | 0.0552 | | 0.1353 |
| Floating % | 0.8734 | 0.8094 | 0.8269 | 0.2574 | 0.9186 | 0.7162 | 0.1386 | 0.7715 | 0.7502 | 0.6004 | 0.9124 | 0.6930 | 0.0998 | 0.3951 | 0.4908 | 0.2711 | 0.1526 | 0.0714 | 0.0316 | 0.1353 | |

Table 9: Correlation Coefficient and corresponding p-value for or all tested parameters for female non-transgenic animals. For the Correlation coefficient red indicates values from 0-0.3, yellow =0.3-0.5 and green 0.5-1. For the p-value red indicates values above 0.15, yellow values from 0.15-0.1 and green values from below 0.05.

| R-Squared | Weight | SVP Density | ICP Density | DCP Density | Total Retina | IRL | RPE | RNFL | IPL | INL | ORL | PRC | OPL | Number of Target Zone Crossings | % of time in sector with platform | Latency on Day 3 | Latency on Day 4 | Distance swam (m) | Velocity (m/s) | Thigmotaxis % | Floating % |
|---|---|---|---|---|---|---|---|---|---|---|---|---|---|---|---|---|---|---|---|---|---|
| Weight | | 0.0049 | 0.1748 | 0.0055 | 0.0203 | 0.0165 | 0.0013 | 0.0152 | 0.0000 | 0.0045 | 0.0227 | 0.0067 | 0.1694 | 0.0171 | 0.0239 | 0.0691 | 0.6042 | 0.4661 | 0.1043 | 0.2187 | 0.0046 |
| SVP Density | 0.0049 | | 0.4395 | 0.0360 | 0.2372 | 0.1987 | 0.2484 | 0.3231 | 0.1463 | 0.0005 | 0.2594 | 0.1799 | 0.3779 | 0.1060 | 0.1298 | 0.1606 | 0.0138 | 0.0011 | 0.0857 | 0.0003 | 0.0105 |
| ICP Density | 0.1748 | 0.4395 | | 0.2441 | 0.0001 | 0.0036 | 0.0146 | 0.0114 | 0.0000 | 0.2141 | 0.0048 | 0.0013 | 0.3390 | 0.3344 | 0.3916 | 0.2661 | 0.1707 | 0.2170 | 0.1117 | 0.2640 | 0.0086 |
| DCP Density | 0.0055 | 0.0360 | 0.2441 | | 0.4109 | 0.5131 | 0.1793 | 0.2947 | 0.1578 | 0.5021 | 0.3197 | 0.3171 | 0.0375 | 0.5369 | 0.5624 | 0.3143 | 0.0200 | 0.0155 | 0.0037 | 0.0138 | 0.2070 |
| Total Retina | 0.0203 | 0.2372 | 0.0001 | 0.4109 | | 0.9690 | 0.2821 | 0.8690 | 0.5938 | 0.6165 | 0.9797 | 0.9454 | 0.1681 | 0.1240 | 0.1408 | 0.0011 | 0.0111 | 0.0067 | 0.0033 | 0.1357 | 0.0019 |
| IRL | 0.0165 | 0.1987 | 0.0036 | 0.5131 | 0.9690 | | 0.2944 | 0.7954 | 0.4584 | 0.6249 | 0.9012 | 0.8636 | 0.1686 | 0.1632 | 0.1877 | 0.0074 | 0.0064 | 0.0018 | 0.0020 | 0.1039 | 0.0236 |
| RPE | 0.0013 | 0.2484 | 0.0146 | 0.1793 | 0.2821 | 0.2944 | | 0.5693 | 0.3872 | 0.0016 | 0.2598 | 0.1466 | 0.6992 | 0.3617 | 0.2814 | 0.0007 | 0.0785 | 0.1603 | 0.2684 | 0.0040 | 0.3270 |
| RNFL | 0.0152 | 0.3231 | 0.0114 | 0.2947 | 0.8690 | 0.7954 | 0.5693 | | 0.8091 | 0.3318 | 0.8904 | 0.7871 | 0.3619 | 0.1986 | 0.1837 | 0.0306 | 0.0001 | 0.0014 | 0.0014 | 0.0986 | 0.0151 |
| IPL | 0.0000 | 0.1463 | 0.0000 | 0.1578 | 0.5938 | 0.4584 | 0.3872 | 0.8091 | | 0.2829 | 0.6868 | 0.6516 | 0.1442 | 0.1885 | 0.1628 | 0.1026 | 0.0231 | 0.0259 | 0.0855 | 0.2474 | 0.0182 |
| INL | 0.0045 | 0.0005 | 0.2141 | 0.5021 | 0.6165 | 0.6249 | 0.0016 | 0.3318 | 0.2829 | | 0.5820 | 0.7189 | 0.0425 | 0.1060 | 0.1556 | 0.0414 | 0.1650 | 0.2164 | 0.2259 | 0.3895 | 0.0485 |
| ORL | 0.0227 | 0.2594 | 0.0048 | 0.3197 | 0.9797 | 0.9012 | 0.2598 | 0.8904 | 0.6868 | 0.5820 | | 0.9702 | 0.1602 | 0.0913 | 0.1023 | 0.0169 | 0.0153 | 0.0126 | 0.0191 | 0.1578 | 0.0022 |
| PRC | 0.0067 | 0.1799 | 0.0013 | 0.3171 | 0.9454 | 0.8636 | 0.1466 | 0.7871 | 0.6516 | 0.7189 | 0.9702 | | 0.0556 | 0.0727 | 0.0910 | 0.0115 | 0.0615 | 0.0657 | 0.0786 | 0.2595 | 0.0278 |
| OPL | 0.1694 | 0.3779 | 0.3390 | 0.0375 | 0.1681 | 0.1686 | 0.6992 | 0.3619 | 0.1442 | 0.0425 | 0.1602 | 0.0556 | | 0.0724 | 0.0396 | 0.0267 | 0.3843 | 0.5295 | 0.5049 | 0.2188 | 0.3866 |
| Number of Target Zone Crossings | 0.0171 | 0.1060 | 0.3344 | 0.5369 | 0.1240 | 0.1632 | 0.3617 | 0.1986 | 0.1885 | 0.1060 | 0.0913 | 0.0727 | 0.0724 | | 0.9834 | 0.1177 | 0.0697 | 0.0612 | 0.0409 | 0.0115 | 0.1226 |
| % of time in sector with platform | 0.0239 | 0.1298 | 0.3916 | 0.5624 | 0.1408 | 0.1877 | 0.2814 | 0.1837 | 0.1628 | 0.1556 | 0.1023 | 0.0910 | 0.0396 | 0.9834 | | 0.1331 | 0.0511 | 0.0398 | 0.0287 | 0.0219 | 0.0823 |
| Latency on Day 3 | 0.0691 | 0.1606 | 0.2661 | 0.3143 | 0.0011 | 0.0074 | 0.0007 | 0.0306 | 0.1026 | 0.0414 | 0.0169 | 0.0115 | 0.0267 | 0.1177 | 0.1331 | | 0.0069 | 0.0138 | 0.0408 | 0.0812 | 0.1966 |
| Latency on Day 4 | 0.6042 | 0.0138 | 0.1707 | 0.0200 | 0.0111 | 0.0064 | 0.0785 | 0.0001 | 0.0231 | 0.1650 | 0.0153 | 0.0615 | 0.3843 | 0.0697 | 0.0511 | 0.0069 | | 0.9515 | 0.4093 | 0.7022 | 0.3089 |
| Distance swam (m) | 0.4661 | 0.0011 | 0.2170 | 0.0155 | 0.0067 | 0.0018 | 0.1603 | 0.0014 | 0.0259 | 0.2164 | 0.0126 | 0.0657 | 0.5295 | 0.0612 | 0.0398 | 0.0138 | 0.9515 | | 0.6047 | 0.7665 | 0.4436 |
| Velocity (m/s) | 0.1043 | 0.0857 | 0.1117 | 0.0037 | 0.0033 | 0.0020 | 0.2684 | 0.0014 | 0.0855 | 0.2259 | 0.0191 | 0.0786 | 0.5049 | 0.0409 | 0.0287 | 0.0408 | 0.4093 | 0.6047 | | 0.4844 | 0.5647 |
| Thigmotaxis % | 0.2187 | 0.0003 | 0.2640 | 0.0138 | 0.1357 | 0.1039 | 0.0040 | 0.0986 | 0.2474 | 0.3895 | 0.1578 | 0.2595 | 0.2188 | 0.0115 | 0.0219 | 0.0812 | 0.7022 | 0.7665 | 0.4844 | | 0.3314 |
| Floating % | 0.0046 | 0.0105 | 0.0086 | 0.2070 | 0.0019 | 0.0236 | 0.3270 | 0.0151 | 0.0182 | 0.0485 | 0.0022 | 0.0278 | 0.3866 | 0.1226 | 0.0823 | 0.1966 | 0.3089 | 0.4436 | 0.5647 | 0.3314 | |

| Mean Squared Error | Weight | SVP Density | ICP Density | DCP Density | Total Retina | IRL | RPE | RNFL | IPL | INL | ORL | PRC | OPL | Number of Target Zone Crossings | % of time in sector with platform | Latency on Day 3 | Latency on Day 4 | Distance swam (m) | Velocity (m/s) | Thigmotaxis % | Floating % |
|---|---|---|---|---|---|---|---|---|---|---|---|---|---|---|---|---|---|---|---|---|---|
| Weight | | 6.8E-05 | 0.00054 | 0.00033 | 114.86 | 23.6588 | 0.862778 | 37.7809 | 11.3469 | 4.39535 | 35.9048 | 32.4513 | 0.85832 | 4.530467446 | 430.5474464 | 128.59837 | 38.259146 | 19101.294 | 2.49839 | 96.23562141 | 61.83143 |
| SVP Density | 6.85E-05 | | 0.00037 | 0.00032 | 89.431 | 19.2754 | 0.64927 | 25.9686 | 9.68704 | 4.41323 | 27.2098 | 26.794 | 0.64283 | 4.120843712 | 383.8181899 | 115.96273 | 95.333736 | 35736.384 | 2.55027 | 123.1391888 | 61.46589 |
| ICP Density | 0.000542 | 0.00037 | | 0.00025 | 117.22 | 23.9679 | 0.851272 | 37.9259 | 11.3468 | 3.47007 | 36.5629 | 32.6278 | 0.68307 | 3.068006653 | 268.3711291 | 101.3796 | 80.162828 | 28013.128 | 2.4777 | 90.65708748 | 61.58069 |
| DCP Density | 0.000329 | 0.00032 | 0.00025 | | 69.066 | 11.713 | 0.708946 | 27.0601 | 9.55656 | 2.19835 | 24.9916 | 22.3113 | 0.99463 | 2.134470562 | 192.9999314 | 94.72908 | 94.738303 | 35223.393 | 2.77915 | 121.481706 | 49.26072 |
| Total Retina | 114.8568 | 89.4308 | 117.221 | 69.0662 | | 0.74461 | 0.620175 | 5.02456 | 4.60875 | 1.69346 | 0.74461 | 1.78401 | 0.85964 | 4.037901445 | 378.9651257 | 137.98483 | 95.595171 | 35536.73 | 2.7802 | 106.46692 | 61.99917 |
| IRL | 23.65875 | 19.2754 | 23.9679 | 11.713 | 0.7446 | | 0.6095 | 7.84878 | 6.14572 | 1.65607 | 3.62893 | 4.4579 | 0.85916 | 3.857033267 | 358.2993982 | 137.11364 | 96.051403 | 35713.807 | 2.78385 | 110.3853575 | 60.64799 |
| RPE | 0.862778 | 0.64927 | 0.85127 | 0.70895 | 0.6202 | 0.6095 | | 16.5255 | 6.95318 | 4.40826 | 27.1937 | 27.8827 | 0.3108 | 2.942012419 | 316.9430122 | 138.04439 | 89.077646 | 30043.586 | 2.04072 | 122.6840153 | 41.80575 |
| RNFL | 37.78088 | 25.9686 | 37.9259 | 27.0601 | 5.0246 | 7.84878 | 16.52546 | | 2.16601 | 2.95049 | 4.02623 | 6.95467 | 0.65936 | 3.69414333 | 360.0412102 | 133.91594 | 96.662672 | 35728.482 | 2.78545 | 111.0396532 | 61.17548 |
| IPL | 11.34693 | 9.68704 | 11.3468 | 9.55656 | 4.6087 | 6.14572 | 6.953178 | 2.16601 | | 3.16619 | 11.5057 | 11.3829 | 0.88432 | 3.74039742 | 369.2950716 | 123.97091 | 94.440352 | 34850.829 | 2.55087 | 92.71210358 | 60.98671 |
| INL | 4.395346 | 4.41323 | 3.47007 | 2.19835 | 1.6935 | 1.65607 | 4.408257 | 2.95049 | 3.16619 | | 15.3562 | 9.18525 | 0.98939 | 4.120863508 | 372.4667858 | 132.41869 | 80.717873 | 28035.837 | 2.15932 | 75.19652745 | 59.10667 |
| ORL | 35.90483 | 27.2098 | 36.5629 | 24.9916 | 0.7446 | 3.62893 | 27.19375 | 4.02623 | 11.5057 | 15.3562 | | 0.97511 | 0.86784 | 4.188746574 | 395.9665208 | 135.80297 | 95.192208 | 35325.326 | 2.73613 | 103.7445024 | 61.98061 |
| PRC | 32.45129 | 26.794 | 32.6278 | 22.3113 | 1.784 | 4.4579 | 27.8827 | 6.95467 | 11.3829 | 9.18525 | 0.97511 | | 0.97586 | 4.274332485 | 400.9559016 | 136.55584 | 90.726905 | 33425.959 | 2.57004 | 91.21784972 | 60.38803 |
| OPL | 0.858319 | 0.64283 | 0.68307 | 0.99463 | 0.8596 | 0.85916 | 0.310799 | 0.65936 | 0.88432 | 0.98939 | 0.86784 | 0.97586 | | 4.275618656 | 423.6114164 | 134.44798 | 59.518746 | 16832.84 | 1.3811 | 96.23367655 | 38.10244 |
| Number of Target Zone Crossings | 4.530467 | 4.12084 | 3.06801 | 2.13447 | 4.0379 | 3.85703 | 2.942012 | 3.69414 | 3.7404 | 4.12086 | 4.18875 | 4.27433 | 4.27562 | | 7.324923099 | 121.88852 | 89.928769 | 33587.657 | 2.67528 | 121.7619291 | 54.49908 |
| % of time in sector with platform | 430.5474 | 383.818 | 268.371 | 193 | 378.97 | 358.299 | 316.943 | 360.041 | 369.295 | 372.467 | 395.967 | 400.956 | 423.611 | 7.324923099 | | 119.76032 | 91.727196 | 34354.089 | 2.70941 | 120.4783246 | 57.00264 |
| Latency on Day 3 | 128.5984 | 115.963 | 101.38 | 94.7291 | 137.98 | 137.114 | 138.0444 | 133.916 | 123.971 | 132.419 | 135.803 | 136.556 | 134.448 | 121.888523 | 119.7603249 | | 95.997246 | 35283.369 | 2.67561 | 113.1748526 | 49.90308 |
| Latency on Day 4 | 38.25915 | 95.3337 | 80.1628 | 94.7383 | 95.595 | 96.0514 | 89.07765 | 96.6627 | 94.4404 | 80.7179 | 95.1922 | 90.7269 | 59.5187 | 89.92876873 | 91.72719565 | 95.997246 | | 1736.6452 | 1.64765 | 36.6882226 | 42.92695 |
| Distance swam (m) | 19101.29 | 35736.4 | 28013.1 | 35223.4 | 35537 | 35713.8 | 30043.59 | 35728.5 | 34850.8 | 28035.8 | 35325.3 | 33426 | 16832.8 | 33587.65701 | 34354.08881 | 35283.369 | 1736.6452 | | 1.10268 | 28.76638276 | 34.56133 |
| Velocity (m/s) | 2.498387 | 2.55027 | 2.4777 | 2.77915 | 2.7802 | 2.78385 | 2.040716 | 2.78545 | 2.55087 | 2.15932 | 2.73613 | 2.57004 | 1.3811 | 2.675278376 | 2.709410625 | 2.6756122 | 1.6476509 | 1.1026779 | | 63.5169528 | 27.04015 |
| Thigmotaxis % | 96.23562 | 123.139 | 90.6571 | 121.482 | 106.47 | 110.385 | 122.684 | 111.04 | 92.7121 | 75.1965 | 103.745 | 91.2178 | 96.2337 | 121.7619291 | 120.4783246 | 113.17485 | 36.688223 | 28.766383 | 63.517 | | 41.52832 |
| Floating % | 61.83143 | 61.4659 | 61.5807 | 49.2607 | 61.999 | 60.648 | 41.80575 | 61.1755 | 60.9867 | 59.1067 | 61.9806 | 60.388 | 38.1024 | 54.49908376 | 57.00263726 | 49.903081 | 42.926953 | 34.56133 | 27.0401 | 41.52832057 | |

Table 10: R-squared and corresponding MSE for or all tested parameters all for female non-transgenic animals. For the R-squared values red indicates values from 0-0.3, yellow =0.4-0.6 and green 0.6-1.

| Correlation Coefficient | Weight | SVP Density | ICP Density | DCP Density | Total Retina | IRL | RPE | RNFL | IPL | INL | ORL | PRC | OPL | Number of Target Zone Crossings | % of time in sector with platform | Latency on Day 3 | Latency on Day 4 | Distance swam (m) | Velocity (m/s) | Thigmotaxis % | Floating % |
|---|---|---|---|---|---|---|---|---|---|---|---|---|---|---|---|---|---|---|---|---|---|
| Weight | | 0.0375 | 0.0803 | -0.0469 | -0.0082 | 0.5634 | 0.4989 | 0.0221 | 0.3618 | -0.2907 | -0.2262 | -0.2608 | 0.0826 | 0.5533 | -0.2867 | -0.4670 | 0.0655 | -0.6240 | -0.4541 | 0.5547 | 0.5589 |
| SVP Density | 0.0375 | | 0.8573 | 0.4484 | -0.9840 | 0.1649 | -0.2584 | -0.4144 | 0.4652 | 0.4884 | -0.9652 | -0.9434 | -0.9253 | 0.6570 | 0.6595 | -0.2450 | -0.3491 | -0.1391 | 0.5610 | -0.1718 | 0.0950 |
| ICP Density | 0.0803 | 0.8573 | | 0.6203 | -0.9320 | -0.1980 | -0.4974 | -0.8020 | 0.8186 | 0.1006 | -0.7768 | -0.7467 | -0.8393 | 0.7940 | 0.8823 | -0.6844 | -0.7983 | -0.5478 | 0.7877 | -0.5662 | -0.3706 |
| DCP Density | -0.0469 | 0.4484 | 0.6203 | | -0.5366 | -0.5903 | -0.8587 | -0.7840 | 0.7165 | -0.4641 | -0.2624 | -0.1927 | -0.7326 | 0.7196 | 0.7850 | -0.2945 | -0.5762 | -0.1203 | 0.8301 | -0.5198 | -0.2686 |
| Total Retina | -0.0082 | -0.9840 | -0.9320 | -0.5366 | | -0.0150 | 0.3805 | 0.5661 | -0.5891 | -0.3771 | 0.9217 | 0.8938 | 0.9369 | -0.7070 | -0.7748 | 0.3938 | 0.5395 | 0.2613 | -0.7024 | 0.3626 | 0.1092 |
| IRL | 0.5634 | 0.1649 | -0.1980 | -0.5903 | -0.0150 | | 0.8835 | 0.6741 | -0.3767 | 0.5494 | -0.4018 | -0.4597 | 0.1197 | -0.0220 | -0.6135 | 0.2590 | 0.7437 | 0.0454 | -0.9069 | 0.9749 | 0.9207 |
| RPE | 0.4989 | -0.2584 | -0.4974 | -0.8587 | 0.3805 | 0.8835 | | 0.7812 | -0.5292 | 0.3862 | 0.0057 | -0.0676 | 0.5572 | -0.3410 | -0.8352 | 0.1908 | 0.6730 | -0.0360 | -0.9741 | 0.8360 | 0.6626 |
| RNFL | 0.0221 | -0.4144 | -0.8020 | -0.7840 | 0.5661 | 0.6741 | 0.7812 | | -0.9212 | 0.4426 | 0.2568 | 0.2092 | 0.5618 | -0.6912 | -0.9062 | 0.7454 | 0.9481 | 0.5799 | -0.8971 | 0.8028 | 0.6655 |
| IPL | 0.3618 | 0.4652 | 0.8186 | 0.7165 | -0.5891 | -0.3767 | -0.5292 | -0.9212 | | -0.4654 | -0.3933 | -0.3624 | -0.5437 | 0.8797 | 0.7561 | -0.8608 | -0.8485 | -0.7652 | 0.6672 | -0.5240 | -0.3848 |
| INL | -0.2907 | 0.4884 | 0.1006 | -0.4641 | -0.3771 | 0.5494 | 0.3862 | 0.4426 | -0.4654 | | -0.5585 | -0.5860 | -0.2340 | -0.2807 | -0.0884 | 0.9799 | 0.9227 | 0.9087 | -0.5764 | 0.6123 | 0.5804 |
| ORL | -0.2262 | -0.9652 | -0.7768 | -0.2624 | 0.9217 | -0.4018 | 0.0057 | 0.2568 | -0.3933 | -0.5585 | | 0.9970 | 0.8116 | -0.6390 | -0.4716 | 0.2815 | 0.1874 | 0.2513 | -0.2753 | -0.1163 | -0.3547 |
| PRC | -0.2608 | -0.9434 | -0.7467 | -0.1927 | 0.8938 | -0.4597 | -0.0676 | 0.2092 | -0.3624 | -0.5860 | 0.9970 | | 0.7639 | -0.6117 | -0.4164 | 0.2989 | 0.1482 | 0.2908 | -0.1922 | -0.1887 | -0.4128 |
| OPL | 0.0826 | -0.9253 | -0.8393 | -0.7326 | 0.9369 | 0.1197 | 0.5572 | 0.5618 | -0.5437 | -0.2340 | 0.8116 | 0.7639 | | -0.7102 | -0.7885 | 0.1353 | 0.3638 | -0.0083 | -0.6727 | 0.2965 | 0.0244 |
| Number of Target Zone Crossings | 0.5533 | 0.6570 | 0.7940 | 0.7196 | -0.7070 | -0.0220 | -0.3410 | -0.6912 | 0.8797 | -0.2807 | -0.6390 | -0.6117 | -0.7102 | | 0.6066 | -0.6195 | -0.5064 | -0.5727 | 0.4285 | -0.1304 | 0.0705 |
| % of time in sector with platform | -0.2867 | 0.6595 | 0.8823 | 0.7850 | -0.7748 | -0.6135 | -0.8352 | -0.9062 | 0.7561 | -0.0884 | -0.4716 | -0.4164 | -0.7885 | 0.6066 | | -0.4883 | -0.8297 | -0.2860 | 0.9743 | -0.8128 | -0.6290 |
| Latency on Day 3 | -0.4670 | -0.2450 | -0.6844 | -0.2945 | 0.3938 | 0.2590 | 0.1908 | 0.7454 | -0.8608 | 0.9799 | 0.2815 | 0.2989 | 0.1353 | -0.6195 | -0.4883 | | 0.8328 | 0.9736 | -0.4019 | 0.4640 | 0.4643 |
| Latency on Day 4 | 0.0655 | -0.3491 | -0.7983 | -0.5762 | 0.5395 | 0.7437 | 0.6730 | 0.9481 | -0.8485 | 0.9227 | 0.1874 | 0.1482 | 0.3638 | -0.5064 | -0.8297 | 0.8328 | | 0.6869 | -0.8227 | 0.8664 | 0.8124 |
| Distance swam (m) | -0.6240 | -0.1391 | -0.5478 | -0.1203 | 0.2613 | 0.0454 | -0.0360 | 0.5799 | -0.7652 | 0.9087 | 0.2513 | 0.2908 | -0.0083 | -0.5727 | -0.2860 | 0.9736 | 0.6869 | | -0.1832 | 0.2655 | 0.2992 |
| Velocity (m/s) | -0.4541 | 0.5610 | 0.7877 | 0.8301 | -0.7024 | -0.9069 | -0.9741 | -0.8971 | 0.6672 | -0.5764 | -0.2753 | -0.1922 | -0.6727 | 0.4285 | 0.9743 | -0.4019 | -0.8227 | -0.1832 | | -0.9059 | -0.7554 |
| Thigmotaxis % | 0.5547 | -0.1718 | -0.5662 | -0.5198 | 0.3626 | 0.9749 | 0.8360 | 0.8028 | -0.5240 | 0.6123 | -0.1163 | -0.1887 | 0.2965 | -0.1304 | -0.8128 | 0.4640 | 0.8664 | 0.2655 | -0.9059 | | 0.9618 |
| Floating % | 0.5589 | 0.0950 | -0.3706 | -0.2686 | 0.1092 | 0.9207 | 0.6626 | 0.6655 | -0.3848 | 0.5804 | -0.3547 | -0.4128 | 0.0244 | 0.0705 | -0.6290 | 0.4643 | 0.8124 | 0.2992 | -0.7554 | 0.9618 | |

| P-Value | Weight | SVP Density | ICP Density | DCP Density | Total Retina | IRL | RPE | RNFL | IPL | INL | ORL | PRC | OPL | Number of Target Zone Crossings | % of time in sector with platform | Latency on Day 3 | Latency on Day 4 | Distance swam (m) | Velocity (m/s) | Thigmotaxis % | Floating % |
|---|---|---|---|---|---|---|---|---|---|---|---|---|---|---|---|---|---|---|---|---|---|
| Weight | | 0.9522 | 0.8979 | 0.9403 | 0.9895 | 0.3227 | 0.3922 | 0.9719 | 0.5496 | 0.6351 | 0.7145 | 0.6718 | 0.8949 | 0.3333 | 0.6400 | 0.5330 | 0.9345 | 0.3760 | 0.5459 | 0.4453 | 0.4411 |
| SVP Density | 0.9522 | | 0.0633 | 0.4488 | 0.0024 | 0.7910 | 0.6746 | 0.4879 | 0.4298 | 0.4038 | 0.0078 | 0.0160 | 0.0242 | 0.2283 | 0.2259 | 0.7550 | 0.6509 | 0.8609 | 0.4390 | 0.8282 | 0.9050 |
| ICP Density | 0.8979 | 0.0633 | | 0.2642 | 0.0211 | 0.7495 | 0.3939 | 0.1026 | 0.0902 | 0.8721 | 0.1223 | 0.1471 | 0.0754 | 0.1087 | 0.0476 | 0.3156 | 0.2017 | 0.4522 | 0.2123 | 0.4338 | 0.6294 |
| DCP Density | 0.9403 | 0.4488 | 0.2642 | | 0.3512 | 0.2947 | 0.0624 | 0.1165 | 0.1733 | 0.4311 | 0.6698 | 0.7562 | 0.1591 | 0.1706 | 0.1157 | 0.7055 | 0.4238 | 0.8797 | 0.1699 | 0.4802 | 0.7314 |
| Total Retina | 0.9895 | 0.0024 | 0.0211 | 0.3512 | | 0.9809 | 0.5274 | 0.3199 | 0.2959 | 0.5315 | 0.0260 | 0.0409 | 0.0189 | 0.1818 | 0.1238 | 0.6062 | 0.4605 | 0.7387 | 0.2976 | 0.6374 | 0.8908 |
| IRL | 0.3227 | 0.7910 | 0.7495 | 0.2947 | 0.9809 | | 0.0469 | 0.2120 | 0.5319 | 0.3374 | 0.5026 | 0.4361 | 0.8480 | 0.9720 | 0.2711 | 0.7410 | 0.2563 | 0.9546 | 0.0931 | 0.0251 | 0.0793 |
| RPE | 0.3922 | 0.6746 | 0.3939 | 0.0624 | 0.5274 | 0.0469 | | 0.1188 | 0.3591 | 0.5207 | 0.9928 | 0.9139 | 0.3292 | 0.5743 | 0.0783 | 0.8092 | 0.3270 | 0.9640 | 0.0259 | 0.1640 | 0.3374 |
| RNFL | 0.9719 | 0.4879 | 0.1026 | 0.1165 | 0.3199 | 0.2120 | 0.1188 | | 0.0262 | 0.4555 | 0.6766 | 0.7356 | 0.3244 | 0.1962 | 0.0340 | 0.2546 | 0.0519 | 0.4201 | 0.1029 | 0.1972 | 0.3345 |
| IPL | 0.5496 | 0.4298 | 0.0902 | 0.1733 | 0.2959 | 0.5319 | 0.3591 | 0.0262 | | 0.4296 | 0.5124 | 0.5489 | 0.3435 | 0.0491 | 0.1392 | 0.1392 | 0.1515 | 0.2348 | 0.3328 | 0.4760 | 0.6152 |
| INL | 0.6351 | 0.4038 | 0.8721 | 0.4311 | 0.5315 | 0.3374 | 0.5207 | 0.4555 | 0.4296 | | 0.3278 | 0.2992 | 0.7049 | 0.6473 | 0.8875 | 0.0201 | 0.0773 | 0.0913 | 0.4236 | 0.3877 | 0.4196 |
| ORL | 0.7145 | 0.0078 | 0.1223 | 0.6698 | 0.0260 | 0.5026 | 0.9928 | 0.6766 | 0.5124 | 0.3278 | | 0.0002 | 0.0954 | 0.2458 | 0.4226 | 0.7185 | 0.8126 | 0.7487 | 0.7247 | 0.8837 | 0.6453 |
| PRC | 0.6718 | 0.0160 | 0.1471 | 0.7562 | 0.0409 | 0.4361 | 0.9139 | 0.7356 | 0.5489 | 0.2992 | 0.0002 | | 0.1327 | 0.2729 | 0.4856 | 0.7011 | 0.8518 | 0.7092 | 0.8078 | 0.8113 | 0.5872 |
| OPL | 0.8949 | 0.0242 | 0.0754 | 0.1591 | 0.0189 | 0.8480 | 0.3292 | 0.3244 | 0.3435 | 0.7049 | 0.0954 | 0.1327 | | 0.1789 | 0.1130 | 0.8647 | 0.6362 | 0.9917 | 0.3273 | 0.7035 | 0.9756 |
| Number of Target Zone Crossings | 0.3333 | 0.2283 | 0.1087 | 0.1706 | 0.1818 | 0.9720 | 0.5743 | 0.1962 | 0.0491 | 0.6473 | 0.2458 | 0.2729 | 0.1789 | | 0.2781 | 0.3805 | 0.4936 | 0.4273 | 0.5715 | 0.8696 | 0.9295 |
| % of time in sector with platform | 0.6400 | 0.2259 | 0.0476 | 0.1157 | 0.1238 | 0.2711 | 0.0783 | 0.0340 | 0.1392 | 0.8875 | 0.4226 | 0.4856 | 0.1130 | 0.2781 | | 0.5117 | 0.1703 | 0.7140 | 0.0257 | 0.1872 | 0.3710 |
| Latency on Day 3 | 0.5330 | 0.7550 | 0.3156 | 0.7055 | 0.6062 | 0.7410 | 0.8092 | 0.2546 | 0.1392 | 0.0201 | 0.7185 | 0.7011 | 0.8647 | 0.3805 | 0.5117 | | 0.1672 | 0.0264 | 0.5981 | 0.5360 | 0.5357 |
| Latency on Day 4 | 0.9345 | 0.6509 | 0.2017 | 0.4238 | 0.4605 | 0.2563 | 0.3270 | 0.0519 | 0.1515 | 0.0773 | 0.8126 | 0.8518 | 0.6362 | 0.4936 | 0.1703 | 0.1672 | | 0.3131 | 0.1773 | 0.1336 | 0.1876 |
| Distance swam (m) | 0.3760 | 0.8609 | 0.4522 | 0.8797 | 0.7387 | 0.9546 | 0.9640 | 0.4201 | 0.2348 | 0.0913 | 0.7487 | 0.7092 | 0.9917 | 0.4273 | 0.7140 | 0.0264 | 0.3131 | | 0.8168 | 0.7345 | 0.7008 |
| Velocity (m/s) | 0.5459 | 0.4390 | 0.2123 | 0.1699 | 0.2976 | 0.0931 | 0.0259 | 0.1029 | 0.3328 | 0.4236 | 0.7247 | 0.8078 | 0.3273 | 0.5715 | 0.0257 | 0.5981 | 0.1773 | 0.8168 | | 0.0941 | 0.2446 |
| Thigmotaxis % | 0.4453 | 0.8282 | 0.4338 | 0.4802 | 0.6374 | 0.0251 | 0.1640 | 0.1972 | 0.4760 | 0.3877 | 0.8837 | 0.8113 | 0.7035 | 0.8696 | 0.1872 | 0.5360 | 0.1336 | 0.7345 | 0.0941 | | 0.0382 |
| Floating % | 0.4411 | 0.9050 | 0.6294 | 0.7314 | 0.8908 | 0.0793 | 0.3374 | 0.3345 | 0.6152 | 0.4196 | 0.6453 | 0.5872 | 0.9756 | 0.9295 | 0.3710 | 0.5357 | 0.1876 | 0.7008 | 0.2446 | 0.0382 | |

Table 11: Correlation Coefficient and corresponding p-value for or all tested parameters for male non-transgenic animals. For the Correlation coefficient red indicates values from 0-0.3, yellow =0.3-0.5 and green 0.5-1. For the p-value red indicates values above 0.15, yellow values from 0.15-0.1 and green values from below 0.05.

| R-Squared | Weight | SVP Density | ICP Density | DCP Density | Total Retina | IRL | RPE | RNFL | IPL | INL | ORL | PRC | OPL | Number of Target Zone Crossings | % of time in sector with platform | Latency on Day 3 | Latency on Day 4 | Distance swam (m) | Velocity (m/s) | Thigmotaxis % | Floating % |
|---|---|---|---|---|---|---|---|---|---|---|---|---|---|---|---|---|---|---|---|---|---|
| Weight | | 0.0014 | 0.0064 | 0.0022 | 0.0001 | 0.3174 | 0.2489 | 0.0005 | 0.1309 | 0.0845 | 0.0511 | 0.0680 | 0.0068 | 0.3062 | 0.0822 | 0.2181 | 0.0043 | 0.3894 | 0.2062 | 0.3076 | 0.3124 |
| SVP Density | 0.0014 | | 0.7350 | 0.2011 | 0.9682 | 0.0272 | 0.0668 | 0.1717 | 0.2164 | 0.2386 | 0.9316 | 0.8901 | 0.8561 | 0.4317 | 0.4349 | 0.0600 | 0.1219 | 0.0193 | 0.3147 | 0.0295 | 0.0090 |
| ICP Density | 0.0064 | 0.7350 | | 0.3848 | 0.8687 | 0.0392 | 0.2474 | 0.6432 | 0.6701 | 0.0101 | 0.6033 | 0.5576 | 0.7045 | 0.6304 | 0.7785 | 0.4685 | 0.6372 | 0.3001 | 0.6204 | 0.3205 | 0.1373 |
| DCP Density | 0.0022 | 0.2011 | 0.3848 | | 0.2879 | 0.3484 | 0.7373 | 0.6147 | 0.5133 | 0.2153 | 0.0688 | 0.0371 | 0.5367 | 0.5178 | 0.6162 | 0.0867 | 0.3319 | 0.0145 | 0.6891 | 0.2702 | 0.0721 |
| Total Retina | 0.0001 | 0.9682 | 0.8687 | 0.2879 | | 0.0002 | 0.1448 | 0.3204 | 0.3470 | 0.1422 | 0.8495 | 0.7990 | 0.8777 | 0.4999 | 0.6004 | 0.1551 | 0.2910 | 0.0683 | 0.4934 | 0.1315 | 0.0119 |
| IRL | 0.3174 | 0.0272 | 0.0392 | 0.3484 | 0.0002 | | 0.7806 | 0.4545 | 0.1419 | 0.3019 | 0.1614 | 0.2113 | 0.0143 | 0.0005 | 0.3763 | 0.0671 | 0.5531 | 0.0021 | 0.8224 | 0.9503 | 0.8477 |
| RPE | 0.2489 | 0.0668 | 0.2474 | 0.7373 | 0.1448 | 0.7806 | | 0.6102 | 0.2801 | 0.1492 | 0.0000 | 0.0046 | 0.3105 | 0.1163 | 0.6976 | 0.0364 | 0.4529 | 0.0013 | 0.9489 | 0.6989 | 0.4390 |
| RNFL | 0.0005 | 0.1717 | 0.6432 | 0.6147 | 0.3204 | 0.4545 | 0.6102 | | 0.8486 | 0.1959 | 0.0660 | 0.0438 | 0.3156 | 0.4777 | 0.8212 | 0.5556 | 0.8989 | 0.3363 | 0.8049 | 0.6445 | 0.4429 |
| IPL | 0.1309 | 0.2164 | 0.6701 | 0.5133 | 0.3470 | 0.1419 | 0.2801 | 0.8486 | | 0.2166 | 0.1547 | 0.1313 | 0.2956 | 0.7740 | 0.5717 | 0.7410 | 0.7199 | 0.5856 | 0.4451 | 0.2746 | 0.1480 |
| INL | 0.0845 | 0.2386 | 0.0101 | 0.2153 | 0.1422 | 0.3019 | 0.1492 | 0.1959 | 0.2166 | | 0.3120 | 0.3433 | 0.0547 | 0.0788 | 0.0078 | 0.9602 | 0.8513 | 0.8258 | 0.3323 | 0.3749 | 0.3369 |
| ORL | 0.0511 | 0.9316 | 0.6033 | 0.0688 | 0.8495 | 0.1614 | 0.0000 | 0.0660 | 0.1547 | 0.3120 | | 0.9940 | 0.6587 | 0.4083 | 0.2224 | 0.0792 | 0.0351 | 0.0632 | 0.0758 | 0.0135 | 0.1258 |
| PRC | 0.0680 | 0.8901 | 0.5576 | 0.0371 | 0.7990 | 0.2113 | 0.0046 | 0.0438 | 0.1313 | 0.3433 | 0.9940 | | 0.5835 | 0.3742 | 0.1734 | 0.0893 | 0.0220 | 0.0845 | 0.0369 | 0.0356 | 0.1704 |
| OPL | 0.0068 | 0.8561 | 0.7045 | 0.5367 | 0.8777 | 0.0143 | 0.3105 | 0.3156 | 0.2956 | 0.0547 | 0.6587 | 0.5835 | | 0.5043 | 0.6217 | 0.0183 | 0.1323 | 0.0001 | 0.4525 | 0.0879 | 0.0006 |
| Number of Target Zone Crossings | 0.3062 | 0.4317 | 0.6304 | 0.5178 | 0.4999 | 0.0005 | 0.1163 | 0.4777 | 0.7740 | 0.0788 | 0.4083 | 0.3742 | 0.5043 | | 0.3680 | 0.3838 | 0.2565 | 0.3280 | 0.1836 | 0.0170 | 0.0050 |
| % of time in sector with platform | 0.0822 | 0.4349 | 0.7785 | 0.6162 | 0.6004 | 0.3763 | 0.6976 | 0.8212 | 0.5717 | 0.0078 | 0.2224 | 0.1734 | 0.6217 | 0.3680 | | 0.2385 | 0.6884 | 0.0818 | 0.9493 | 0.6607 | 0.3957 |
| Latency on Day 3 | 0.2181 | 0.0600 | 0.4685 | 0.0867 | 0.1551 | 0.0671 | 0.0364 | 0.5556 | 0.7410 | 0.9602 | 0.0792 | 0.0893 | 0.0183 | 0.3838 | 0.2385 | | 0.6935 | 0.9480 | 0.1615 | 0.2153 | 0.2156 |
| Latency on Day 4 | 0.0043 | 0.1219 | 0.6372 | 0.3319 | 0.2910 | 0.5531 | 0.4529 | 0.8989 | 0.7199 | 0.8513 | 0.0351 | 0.0220 | 0.1323 | 0.2565 | 0.6884 | 0.6935 | | 0.4718 | 0.6768 | 0.7507 | 0.6599 |
| Distance swam (m) | 0.3894 | 0.0193 | 0.3001 | 0.0145 | 0.0683 | 0.0021 | 0.0013 | 0.3363 | 0.5856 | 0.8258 | 0.0632 | 0.0845 | 0.0001 | 0.3280 | 0.0818 | 0.9480 | 0.4718 | | 0.0336 | 0.0705 | 0.0895 |
| Velocity (m/s) | 0.2062 | 0.3147 | 0.6204 | 0.6891 | 0.4934 | 0.8224 | 0.9489 | 0.8049 | 0.4451 | 0.3323 | 0.0758 | 0.0369 | 0.4525 | 0.1836 | 0.9493 | 0.1615 | 0.6768 | 0.0336 | | 0.8206 | 0.5706 |
| Thigmotaxis % | 0.3076 | 0.0295 | 0.3205 | 0.2702 | 0.1315 | 0.9503 | 0.6989 | 0.6445 | 0.2746 | 0.3749 | 0.0135 | 0.0356 | 0.0879 | 0.0170 | 0.6607 | 0.2153 | 0.7507 | 0.0705 | 0.8206 | | 0.9251 |
| Floating % | 0.3124 | 0.0090 | 0.1373 | 0.0721 | 0.0119 | 0.8477 | 0.4390 | 0.4429 | 0.1480 | 0.3369 | 0.1258 | 0.1704 | 0.0006 | 0.0050 | 0.3957 | 0.2156 | 0.6599 | 0.0895 | 0.5706 | 0.9251 | |

| Mean Squared Error | Weight | SVP Density | ICP Density | DCP Density | Total Retina | IRL | RPE | RNFL | IPL | INL | ORL | PRC | OPL | Number of Target Zone Crossings | % of time in sector with platform | Latency on Day 3 | Latency on Day 4 | Distance swam (m) | Velocity (m/s) | Thigmotaxis % | Floating % |
|---|---|---|---|---|---|---|---|---|---|---|---|---|---|---|---|---|---|---|---|---|---|
| Weight | | 0.00014 | 5.09E-05 | 0.0001851 | 13.7135 | 1.68062 | 0.10007 | 11.0619 | 6.78803 | 0.4363 | 15.5142 | 12.489 | 0.20826 | 3.607894737 | 285.0386308 | 166.606 | 187.1345 | 7851.7885 | 9.61774 | 122.5694663 | 71.65259 |
| SVP Density | 0.000136 | | 1.36E-05 | 0.0001482 | 0.43617 | 2.3951 | 0.12432 | 9.16705 | 6.12029 | 0.36288 | 1.1191 | 1.47274 | 0.03017 | 2.955200041 | 175.4849022 | 200.2779 | 165.0327 | 12609.977 | 8.303371 | 171.8052101 | 103.2611 |
| ICP Density | 5.09E-05 | 1.4E-05 | | 0.0001141 | 1.80086 | 2.36552 | 0.10027 | 3.94932 | 2.57671 | 0.47175 | 6.48554 | 5.92849 | 0.06197 | 1.921974568 | 68.795818 | 113.2558 | 68.18276 | 8999.8975 | 4.599234 | 120.2854882 | 89.89221 |
| DCP Density | 0.000185 | 0.00015 | 0.000114 | | 9.76581 | 1.6042 | 0.035 | 4.26436 | 3.80103 | 0.37395 | 15.225 | 12.9028 | 0.09714 | 2.507679877 | 119.1871817 | 194.5872 | 125.5532 | 12672.459 | 3.766591 | 129.1923886 | 96.68517 |
| Total Retina | 13.71354 | 0.43617 | 1.800865 | 9.7658052 | | 2.46153 | 0.11393 | 7.52111 | 5.10023 | 0.40881 | 2.46153 | 2.694 | 0.02564 | 2.600506435 | 124.1114943 | 180.0258 | 133.2438 | 11980.449 | 6.138515 | 153.7604597 | 102.9595 |
| IRL | 1.680616 | 2.3951 | 2.365516 | 1.6042044 | 2.46153 | | 0.02922 | 6.03769 | 6.70204 | 0.33272 | 13.7114 | 10.569 | 0.20669 | 5.197490939 | 193.6867397 | 198.7718 | 83.99595 | 12832.235 | 2.151644 | 8.792553498 | 15.8656 |
| RPE | 0.100066 | 0.12432 | 0.100266 | 0.0349955 | 0.11393 | 0.02922 | | 4.31401 | 5.62316 | 0.40548 | 16.35 | 13.3389 | 0.14458 | 4.595163707 | 93.90947675 | 205.3091 | 102.8289 | 12842.044 | 0.61887 | 53.30400895 | 58.4559 |
| RNFL | 11.06188 | 9.16705 | 3.949321 | 4.2643623 | 7.52111 | 6.03769 | 4.31401 | | 1.18249 | 0.38322 | 15.272 | 12.8138 | 0.14352 | 2.715896664 | 55.54191597 | 94.68775 | 19.00193 | 8534.4454 | 2.364227 | 62.93140751 | 58.05304 |
| IPL | 6.788029 | 6.12029 | 2.576708 | 3.801028 | 5.10023 | 6.70204 | 5.62316 | 1.18249 | | 0.37336 | 13.8212 | 11.6407 | 0.1477 | 1.175433716 | 133.0194146 | 55.1854 | 52.63283 | 5329.023 | 6.723252 | 128.4159372 | 88.77589 |
| INL | 0.436296 | 0.36288 | 0.471754 | 0.3739473 | 0.40881 | 0.33272 | 0.40548 | 0.38322 | 0.37336 | | 11.2496 | 8.79943 | 0.19822 | 4.79025151 | 308.1344424 | 8.470873 | 27.93961 | 2240.4087 | 8.09033 | 110.6616624 | 69.09802 |
| ORL | 15.51421 | 1.1191 | 6.485539 | 15.224985 | 2.46153 | 13.7114 | 16.35 | 15.272 | 13.8212 | 11.2496 | | 0.08042 | 0.07158 | 3.076624377 | 241.4989548 | 196.1867 | 181.3422 | 12046.608 | 11.19804 | 174.6384606 | 91.08837 |
| PRC | 12.48899 | 1.47274 | 5.928493 | 12.902799 | 2.694 | 10.569 | 13.3389 | 12.8138 | 11.6407 | 8.79943 | 0.08042 | | 0.08733 | 3.254308248 | 256.7272547 | 194.035 | 183.8103 | 11771.587 | 11.66862 | 170.7251365 | 86.44578 |
| OPL | 0.208265 | 0.03017 | 0.061974 | 0.0971426 | 0.02564 | 0.20669 | 0.14458 | 0.14352 | 0.1477 | 0.19822 | 0.07158 | 0.08733 | | 2.577425163 | 117.475555 | 209.1671 | 163.0667 | 12857.815 | 6.633848 | 161.4655513 | 104.1394 |
| Number of Target Zone Crossings | 3.607895 | 2.9552 | 1.921975 | 2.5076799 | 2.60051 | 5.19749 | 4.59516 | 2.7159 | 1.17543 | 4.79025 | 3.07662 | 3.25431 | 2.57743 | | 196.2893941 | 131.2954 | 139.7386 | 8641.0985 | 9.891138 | 174.0216512 | 103.6843 |
| % of time in sector with platform | 285.0386 | 175.485 | 68.79582 | 119.18718 | 124.111 | 193.687 | 93.9095 | 55.5419 | 133.019 | 308.134 | 241.499 | 256.727 | 117.476 | 196.2893941 | | 162.2582 | 58.5615 | 11807.12 | 0.613792 | 60.07134553 | 62.96898 |
| Latency on Day 3 | 166.606 | 200.278 | 113.2558 | 194.58723 | 180.026 | 198.772 | 205.309 | 94.6877 | 55.1854 | 8.47087 | 196.187 | 194.035 | 209.167 | 131.2954487 | 162.258232 | | 57.59861 | 668.99503 | 10.1587 | 138.9239041 | 81.73392 |
| Latency on Day 4 | 187.1345 | 165.033 | 68.18276 | 125.55321 | 133.244 | 83.9959 | 102.829 | 19.0019 | 52.6328 | 27.9396 | 181.342 | 183.81 | 163.067 | 139.7386266 | 58.56149891 | 57.59861 | | 6791.6285 | 3.915639 | 44.13628579 | 35.43657 |
| Distance swam (m) | 7851.789 | 12610 | 8999.898 | 12672.459 | 11980.4 | 12832.2 | 12842 | 8534.45 | 5329.02 | 2240.41 | 12046.6 | 11771.6 | 12857.8 | 8641.098471 | 11807.12008 | 668.995 | 6791.628 | | 11.70951 | 164.5501029 | 94.87433 |
| Velocity (m/s) | 9.61774 | 8.30337 | 4.599234 | 3.7665908 | 6.13851 | 2.15164 | 0.61887 | 2.36423 | 6.72325 | 8.09033 | 11.198 | 11.6686 | 6.63385 | 9.891138456 | 0.613792428 | 10.1587 | 3.915639 | 11.709514 | | 31.75477952 | 44.74547 |
| Thigmotaxis % | 122.5695 | 171.805 | 120.2855 | 129.19239 | 153.76 | 8.79255 | 53.304 | 62.9314 | 128.416 | 110.662 | 174.638 | 170.725 | 161.466 | 174.0216512 | 60.07134553 | 138.9239 | 44.13629 | 164.5501 | 31.75478 | | 7.809298 |
| Floating % | 71.65259 | 103.261 | 89.89221 | 96.685174 | 102.96 | 15.8656 | 58.4559 | 58.053 | 88.7759 | 69.098 | 91.0884 | 86.4458 | 104.139 | 103.6843105 | 62.96897569 | 81.73392 | 35.43657 | 94.874331 | 44.74547 | 7.809298045 | |

Table 12: R-squared and corresponding MSE for or all tested parameters all for male non-transgenic animals. For the R-squared values red indicates values from 0-0.3, yellow =0.4-0.6 and green 0.6-1.
